# CAM evolution is associated with gene family expansion in an explosive bromeliad radiation

**DOI:** 10.1101/2023.02.01.526631

**Authors:** Clara Groot Crego, Jaqueline Hess, Gil Yardeni, Marylaure de La Harpe, Clara Priemer, Francesca Beclin, Sarah Saadain, Luiz A. Cauz-Santos, Eva M. Temsch, Hanna Weiss-Schneeweiss, Michael H.J. Barfuss, Walter Till, Wolfram Weckwerth, Karolina Heyduk, Christian Lexer, Ovidiu Paun, Thibault Leroy

**Affiliations:** Department of Botany and Biodiversity Research, University of Vienna, Vienna, Austria; Vienna Graduate School of Population Genetics, Vienna, Austria; Cambrium GmbH, Max-Urich-Str. 3, 13055 Berlin, Germany; Institute of Computational Biology, Department of Biotechnology, University of Life Sciences and Natural Resources (BOKU), Muthgasse 18, 1190 Vienna, Austria; Office for Nature and Environment, Canton of Grisons, Chur, Switzerland; Department of Functional and Evolutionary Ecology, Molecular Systems Biology (MOSYS), University of Vienna, Vienna, Austria; Gregor Mendel Institute, Austrian Academy of Sciences, Vienna BioCenter, Vienna, Austria; Vienna Metabolomics Center (VIME), University of Vienna, Vienna, Austria; Ecology and Evolutionary Biology, University of Connecticut, Storrs, CT USA; GenPhySE, Université de Toulouse, INRAE, ENVT, Castanet Tolosan, France

## Abstract

The subgenus *Tillandsia* (Bromeliaceae) belongs to one of the fastest radiating clades in the plant kingdom and is characterised by the repeated evolution of Crassulacean Acid Metabolism (CAM). Despite its complex genetic basis, this water-conserving trait has evolved independently across many plant families and is regarded as a key innovation trait and driver of ecological diversification in Bromeliaceae. By producing high-quality genome assemblies of a *Tillandsia* species pair displaying divergent photosynthetic phenotypes, and combining genome-wide investigations of synteny, TE dynamics, sequence evolution, gene family evolution and temporal differential expression, we were able to pinpoint the genomic drivers of CAM evolution in *Tillandsia*. Several large-scale rearrangements associated with karyotype changes between the two genomes and a highly dynamic TE landscape shaped the genomes of *Tillandsia*. However, our analyses show that rewiring of photosynthetic metabolism is mainly obtained through regulatory evolution rather than coding sequence evolution, as CAM-related genes are differentially expressed across a 24-hour cycle between the two species, but are no candidates of positive selection. Gene orthology analyses reveal that CAM-related gene families manifesting differential expression underwent accelerated gene family expansion in the constitutive CAM species, further supporting the view of gene family evolution as a driver of CAM evolution.

## 2. Introduction

Crassulacean Acid Metabolism (CAM) is a photosynthetic phenotype playing a major role in plant adaptation to arid environments and the epiphytic lifeform (Winter and Smith, 2012; Cushman, 2001; Silvera et al., 2010), and has been described as a key innovation trait driving plant diversification and speciation in several plant lineages (Quezada and Gianoli, 2011; Silvera et al., 2009; Ogburn and Edwards, 2009). CAM functions as a carbon concentrating mechanism by assimilating CO_2_ overnight and storing it as malate in the vacuole, which greatly enhances the efficiency of Rubisco, the first enzyme of the Calvin cycle (Osmond, 1978). This also has the secondary effect of improving the plant’s overall water use efficiency by reducing evapotranspiration, as stomata can remain closed during the day (Borland et al., 2014). Though often presented as a discrete trait, CAM actually encompasses a large spectrum of photosynthetic phenotypes including intermediate and facultative forms (Edwards, 2023). Phenotypes from this CAM continuum have evolved repeatedly in at least 37 plant families (Winter et al., 2021), yet the underlying evolutionary mechanisms allowing this complex and diverse trait to emerge multiple times throughout plant history are not fully understood.

Due to the sparse availability of CAM plant genomes, most studies on CAM evolution have focussed on transcription levels and sequence evolution to understand its underlying genetic drivers. However, novel variation can be generated by other mechanisms which have not been investigated thoroughly in the context of CAM evolution. For example, several studies have suggested a potential importance of gene family expansion as a driver of CAM evolution (Cai et al., 2015; Silvera et al., 2014). In C4 plants, duplicated gene copies tend to be more often retained compared to closely related C3 lineages (Hoang et al., 2023). Gene duplication occurs at higher rates than point mutation in many lineages (Katju and Bergthorsson, 2013) and can lead to novel functional variation through dosage effects, neo-functionalization, or subfunctionalization (Ohno, 1970), as observed in teleost fish (Arnegard et al., 2010; Moriyama et al., 2016) and orchids (Mondragón-Palomino and Theissen, 2009). Another form of structural variation that can contribute to the evolution of complex traits is TE insertion in and around genes, which has been shown to play a role in local adaptation, for example in *Arabidopsis* (Baduel et al., 2019). Finally, large-scale rearrangements such as chromosomal fusions, inversions or translocations can increase linkage between co-adapted alleles and generate reproductive barriers (Luo et al., 2018; Lowry and Willis, 2010).

The adaptive radiation of *Tillandsia* subgenus *Tillandsia* (Bromeliaceae) is part of one of the fastest diversifying clades known in the plant kingdom (Tillandsioideae) (Givnish et al., 2014) and is characterised by a number of key innovation traits such as the epiphytic lifestyle, absorptive trichomes, water-impounding tanks and photosynthetic metabolism driving extraordinary diversity both on the taxonomic and ecological level (Barfuss et al., 2016). The group displays a broad range of phenotypes of the CAM continuum, resulting from repeated evolution of constitutive CAM (Crayn et al., 2015; De La Harpe et al., 2020). CAM evolution has been described as an ecological driver of diversification in the subgenus *Tillandsia* (Barfuss et al., 2016; Crayn et al., 2004), and across Bromeliaceae in general (Givnish et al., 2014; Crayn et al., 2004; Benzing and Bennett, 2000). This renders the radiation a fascinating system both for studies on speciation and rapid adaptation generally, and for studies on CAM evolution specifically, as comparative investigations between recently diverged species with contrasting phenotypes prevent the overestimation of evolutionary changes needed to evolve adaptations such as CAM (Heyduk et al., 2019a).

While Bromeliaceae is generally regarded as a homoploid radiation with conserved chromosome counts and little genome size variation (Gitaí et al., 2014), more recent work has pointed at a high “genomic potential” of the subgenus *Tillandsia*, notably from elevated gene loss and duplication rates (De La Harpe et al., 2020), providing an exemplary system to study the role of genome evolution and structural variation in CAM evolution. Not only are adaptive radiations like *Tillandsia* characterised by repeated evolution of key innovation traits, the short timescales at which novel variation arises in these systems challenge classical views of adaptive evolution, stimulating a range of studies pointing at the potential importance of genome evolution as a genomic driver of diversification (McGee et al., 2020; Brawand et al., 2015; Cicconardi et al., 2021).

In this study, we comparatively investigated *de novo* assembled genomes of two ecologically divergent members of the subgenus *Tillandsia* to further our understanding of genome evolution in this recent radiation and its link to CAM evolution as a key innovation trait. *Tillandsia fasciculata* (Fig. 1a) displays a set of phenotypes typically described as “grey” or “atmospheric” *Tillandsia* (Benzing and Bennett, 2000): a dense layer of absorptive, umbrella-shaped trichomes, CAM photosynthesis and occurrence in arid places with high solar incidence and low rainfall. On the other hand, *T. leiboldiana* (Fig. 1b) represents a typical “green” *Tillandsia* displaying tank formation, C3-like leaf morphology, a sparse layer of absorptive trichomes and occurrences in cooler, wetter regions. While not sister species, *T. fasciculata* and *T. leiboldiana* belong to sister clades displaying a shift in photosynthetic metabolism (Fig. 1c), and represent phenotypic extremes within subgenus *Tillandsia*. Their photosynthetic metabolisms have been described as strong CAM for *T. fasciculata,* and C3 for *T. leiboldiana* based on carbon isotope ratios (δ^13^C_T. fasciculata_ = −11.9 / −16.1; δ^13^C_T.leiboldiana_ = −28.0 / −31.3) reported by Crayn et al., (2015) and De La Harpe et al., (2020) respectively. These are some of the most distinct values reported for the subgenus.

**Figure 1:**
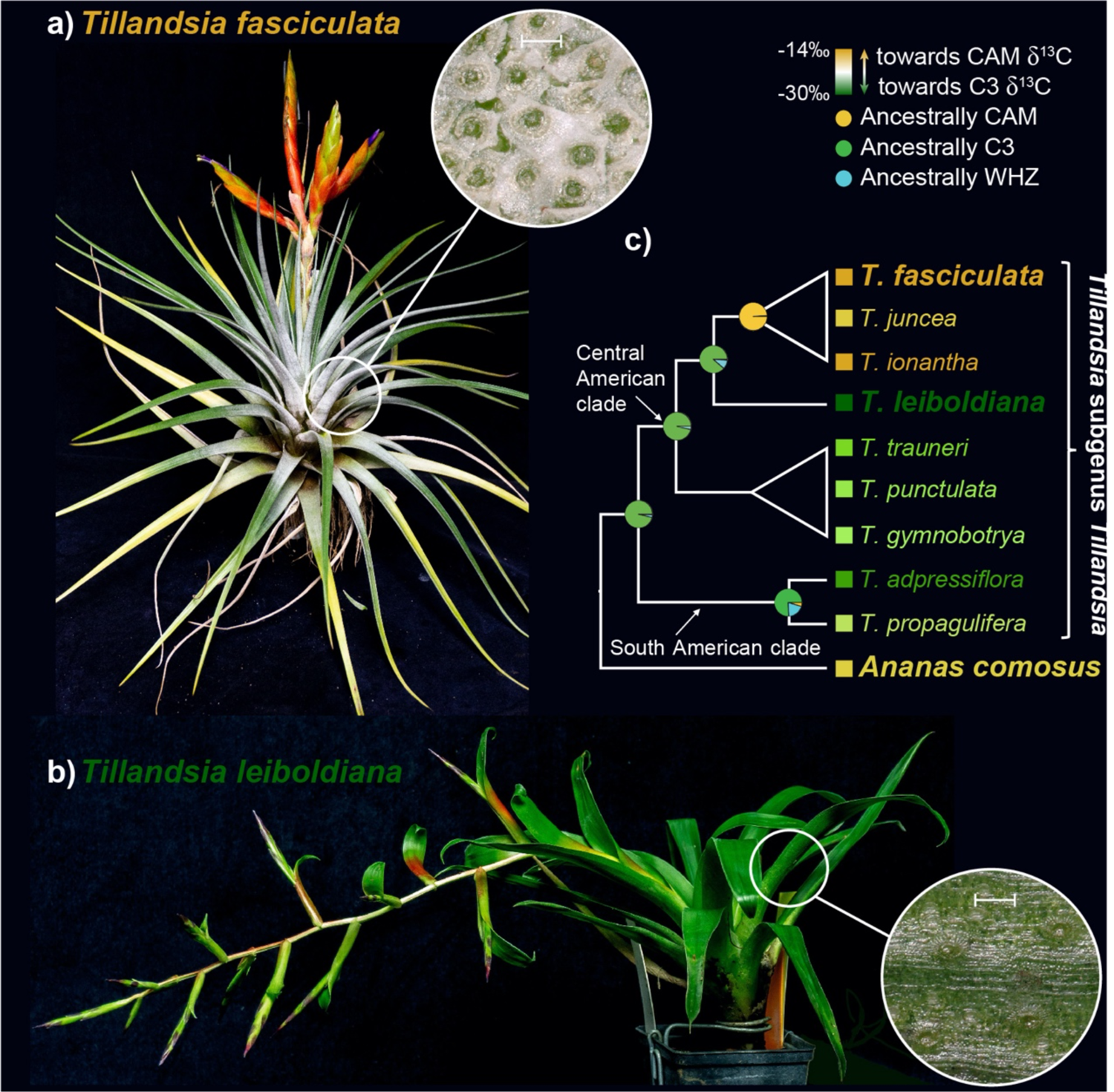
**a)** Tillandsia fasciculata, a “grey” or “atmospheric” Tillandsia with a dense layer of umbrella-shaped trichomes (inset), carbon isotope values within the CAM range, a lack of water-impounding tank and roots adapted to the epiphytic lifestyle. The leaf close-up is at a 100 μm scale (also in b). **b)** Tillandsia leiboldiana, a green Tillandsia with C3-like leaf morphology and carbon isotope values, an impounding tank and a sparse trichome layer (inset). **c)** Schematic representation of the evolutionary relationship between the two investigated species of Tillandsia within the subgenus. Colours indicate reported carbon isotope values (De La Harpe et al., 2020; Crayn et al., 2015). The average was taken when multiple values have been reported for the same species. Pie charts at internal nodes show the ancestral state of photosynthetic metabolism as reported in De La Harpe et al. 2020. WHZ stands for Winter-Holtum Zone (Males, 2018) and represents intermediate forms of the CAM continuum.

However, due to the limited ability of carbon isotope measurements in capturing intermediate CAM phenotypes and therefore representing the full CAM continuum (Messerschmid et al., 2021; Pierce et al., 2002), the exact photosynthetic phenotypes of *T. leiboldiana* and *T. fasciculata* need to be corroborated to fully understand what range of the CAM continuum is truly encompassed in the subgenus. By characterising the photosynthetic metabolisms of *T. leiboldiana* and *T. fasciculata* and investigating genomic variation between these two species on multiple levels, from karyotype, chromosomal rearrangements, to molecular evolution, gene family evolution and temporal differential gene expression, we thoroughly explore the degree of genomic divergence found within this radiation and the link between this variation and the evolution of a key innovation trait. We ascertain that the photosynthetic metabolisms of these species are clearly distinct, with *T. fasciculata* at the late stages of CAM evolution (i.e., constitutive, strong CAM), and *T. leiboldiana* likely at the very early stages (i.e., no night-time malate accumulation, but CAM-like expression profiles of certain enzymes). We further document karyotype differences, multiple chromosomal rearrangements, distinct TE landscapes and gene family evolution rates between the two species. We find evidence that molecular variation underlying the difference in phenotype is largely found at the transcriptomic level. We also find a clear association between CAM-related temporal gene expression differences and both gene family expansion in the constitutive CAM plant and pre-existing gene duplications shared between both species.

## 3. Results

### 3.1. Photosynthetic phenotypes of *T. fasciculata* and *T. leiboldiana*

To better understand the difference in photosynthetic metabolism between *T. fasciculata* and *T. leiboldiana*, we measured metabolite abundances with GC-MS for six samples per species at six timepoints across a 24-hour cycle. The overall composition of 49 metabolite abundances separate samples of both species within the first two principal components (Fig. 2a), which combined explain 41.5 % of variance. This suggests a pronounced general metabolic differentiation between *T. fasciculata* and *T. leiboldiana*. Besides amino acids, the organic acids malate, citrate and gluconic acid contribute most to this differentiation along PC1 (Fig. S1), which is a common pattern for species with diverging photosynthetic metabolism (De La Harpe et al., 2020; Benzing and Bennett, 2000; Popp et al., 2003). Sugars appear large contributors to differentiation along PC2 (Fig. S1).

**Figure 2:**
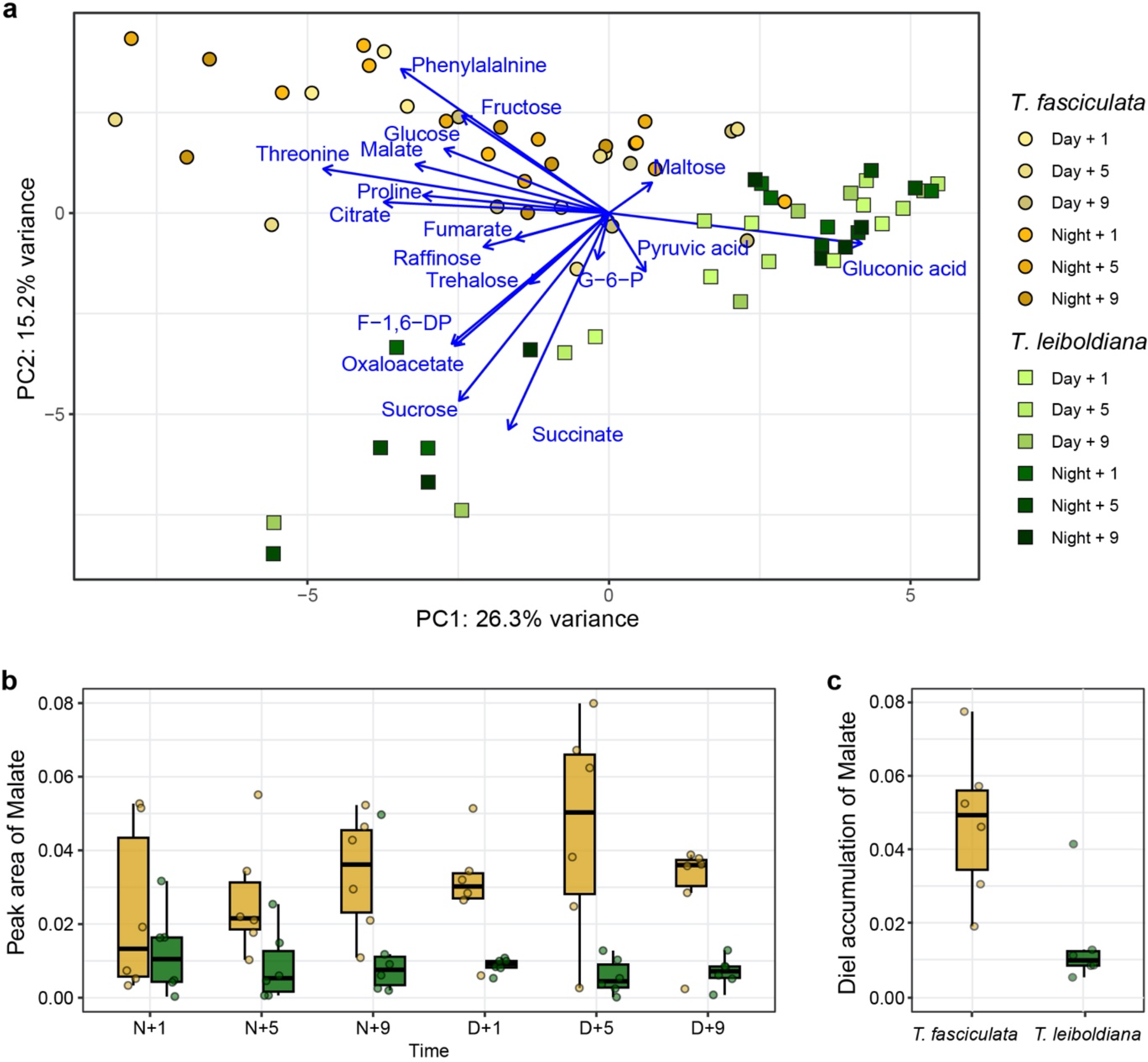
Metabolomic analyses of T. fasciculata and T. leiboldiana leaf material throughout a 24-hour cycle. Abundances of individual metabolites were measured with GC-MS and normalised against the Main Total Ion Count (MTIC). **a)** Principal component analysis of metabolic composition of 72 leaf samples based on 77 metabolic compounds including soluble sugars, amino acids, and organic acids. The first and second principal components are displayed. Blue arrows show the loadings of a subset of metabolites relevant for photosynthetic metabolism. **b)** Malate abundance in leaf material of T. fasciculata and T. leiboldiana at six timepoints across a 24-hour cycle. Dots represent individual observations across timepoints. Timepoints are noted as hours into the day (D) or into the night (N). **c)** Distribution of per-individual accumulation of malate per species over a 24-hour cycle. Accumulations were obtained by taking the difference in malate abundance between the highest and lowest reported abundances across time for each accession.

Overnight malate accumulation is a core feature of CAM photosynthesis and therefore an indicator of the respective photosynthetic phenotypes of *T. fasciculata* and *T. leiboldiana*. Malate abundances in the leaf fluctuated strongly in *T. fasciculata* over 24 hours, with highest median abundances around midday (D+5) and lowest abundances in the early night (N+1), representing a 3.8-fold difference (Fig. 2b). In comparison, malate abundances were overall lower for *T. leiboldiana* and fluctuated less. The highest median abundance in the latter species was found at N+1, while the lowest was found at D+5, representing a 2.2-fold difference. Interestingly, the accumulation times of malate seem reversed in the two species, with the highest abundances in *T. fasciculata* found at the time of lowest abundances in *T. leiboldiana* and vice versa. The reversed timing of malate accumulation has been described as a key difference between C3 and CAM metabolisms ((Winter and Smith, 2022), but see also (Bräutigam et al., 2017)). The median accumulation in malate within 24-hours differs significantly between, with a 3.2-fold higher value in *T. fasciculata* than in *T. leiboldiana* (Wilcoxon rank-sum test p-value = 8.6E-03, Fig. 2c).

Overall, the malate accumulation curves suggest distinct photosynthetic phenotypes for *T. fasciculata* and *T. leiboldiana*. *T. fasciculata* appears to behave as a constitutive CAM plant in standard conditions, accumulating malate from the early night until the early day, while *T. leiboldiana*’s flux is more C3-like, without a clear accumulation overnight.

### 3.2. Genome assembly and annotation

We constructed *de novo* haploid genome assemblies for both species (Table S1) using a combination of long-read (PacBio), short read (Illumina) and chromosome conformation capture (Hi-C) data. This resulted in assemblies of 838 Mb and 1,198 Mb with an N50 of 23.6 and 43.3 Mb in *T. fasciculata* and *T. leiboldiana* respectively. The assembly sizes closely match the estimated genome size of each species based on flow cytometry and k-mer analysis (Table S2, SI Notes 1-2). The 25 and respectively 26 longest scaffolds (hereafter referred to as ‘main scaffolds’) contain 72 % and 75.5 % of the full assembly, after which scaffold sizes steeply decline (SI Note 3, Fig. S2). The number of main scaffolds corresponds with the species karyotype in *T. fasciculata*, but deviates from the *T. leiboldiana* karyotype (SI Note 1), suggesting that a few fragmented chromosome sequences remain in the latter assembly.

Structural gene annotation resulted in a total of 34,886 and 38,180 gene models in *T. fasciculata* and *T. leiboldiana* respectively, of which 92.6 % and 71.9 % are considered robust based on additional curation (Methods, Section 5). Annotation completeness was evaluated with BUSCO using the Liliopsida dataset resulting in a score of 89.7 % complete genes in *T. fasciculata* and 85.3 % in *T. leiboldiana* (Table S2).

### 3.3. Genic, repetitive and GC content

TE annotation performed with EDTA (Ou et al., 2019) revealed a total repetitive content of 65.5 % and 77.1 % in *T. fasciculata* and *T. leiboldiana* respectively. This closely matches estimates derived from k-mer analyses (66 % and 75 %, SI Note 2). Compared to *T. fasciculata*, the repetitive content in *T. leiboldiana* is enriched for Gypsy LTR retrotransposon and *Mutator* DNA transposon content, with a 1.7-fold and 4.2-fold increase in total covered genomic length, respectively (Table S3).

Repetitive content per scaffold is negatively correlated with gene count in both assemblies (Kendall’s correlation coefficient: −0.79 in *T. fasciculata*, −0.82 in *T. leiboldiana,* p-values < 2.2E-16), with gene-rich regions in distal positions (Fig. 3a, green track) and repetitive regions primarily in median positions (Fig. 3a, yellow track). This pattern is accentuated in *T. leiboldiana*: on average, the repetitive-to-exonic content per scaffold is 1.6 times larger compared to *T. fasciculata* (Mann Whitney U, p-value = 4.3E-04). The genome size difference between the two assemblies is therefore mostly explained by differential accumulation of TE content, mostly in heterochromatic regions.

**Figure 3:**
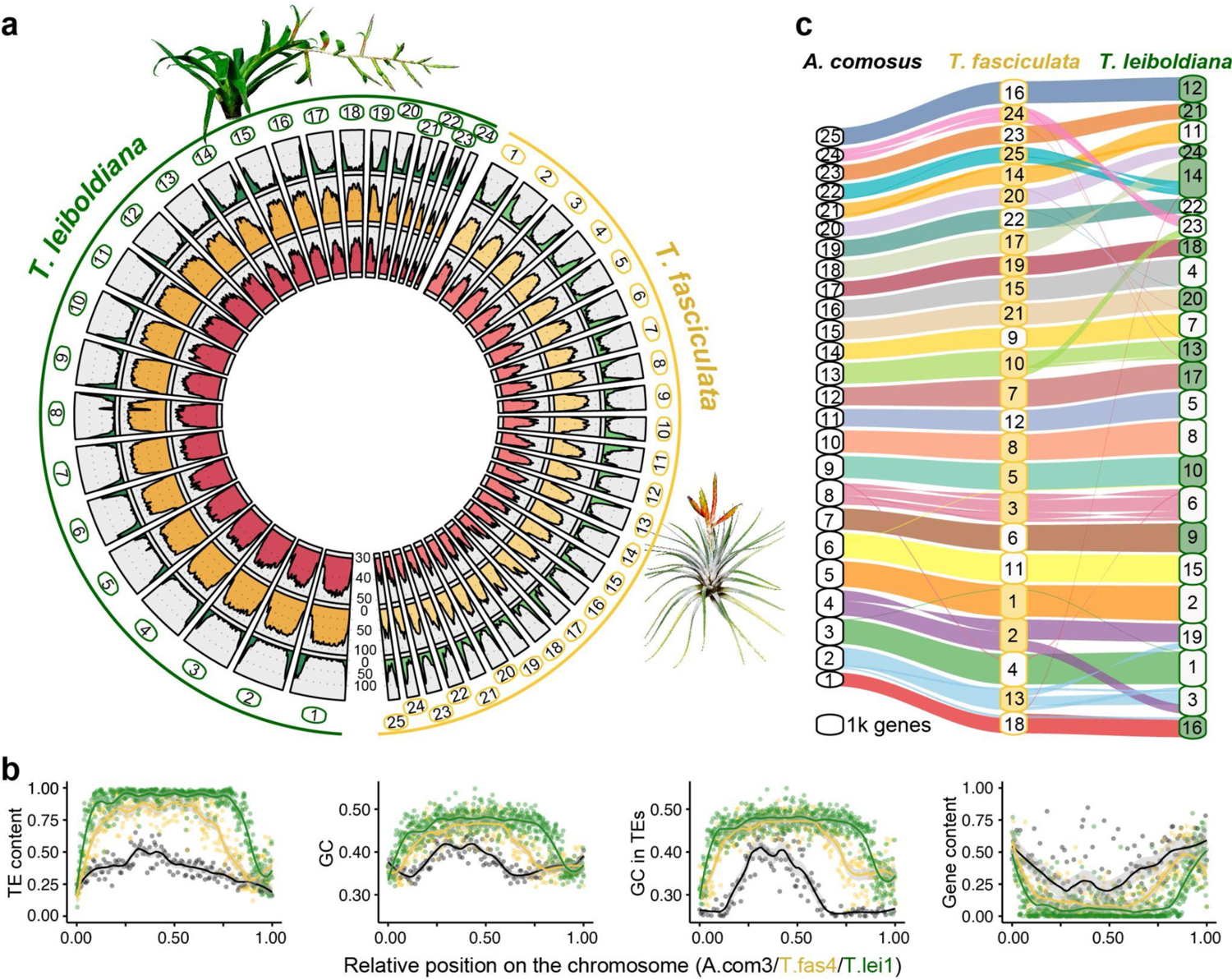
**a)** Circular overview of the main scaffolds of the T. fasciculata (right) and T. leiboldiana (left) genome assemblies. Scaffolds 25 and 26 of T. leiboldiana are not shown due to their reduced size. Going inwards, the tracks show: (1, green) gene count; (2, yellow) proportion of repetitive content; (3, red), and GC content per 1-Mb windows. **b)** TE and GC content, GC content exclusively in TEs, and genic content in a triplet of syntenic scaffolds between Ananas comosus (LG3, black), T. fasciculata (scaffold 4, grey) and T. leiboldiana (scaffold 1, green; see Fig. S3 for other syntenic chromosomes). **c)** Syntenic plot linking blocks of orthologous genes between A. comosus, T. fasciculata and T. leiboldiana. The size of each scaffold on the y-axis is proportional to genic content and therefore does not represent the true scaffold size. Colour-filled boxes indicate scaffolds with reversed coordinates as compared to the sequences in A. comosus.

GC content is negatively correlated with gene content in both species (Kendall’s correlation coefficient: −0.68 in *T. fasciculata*, −0.71 in *T. leiboldiana,* p-values < 2.2E-16, red track in Fig. 3a, detailed in Fig. 3b). By visualising GC and TE content across a syntenic chromosome triplet of *A. comosus*, *T. fasciculata* and *T. leiboldiana,* we show that this relationship can be mostly explained by elevated GC content in repetitive regions (Fig. 3b). TE-rich regions indeed exhibit a much higher GC content than TE-poor regions, a pattern which is exacerbated as the overall TE content per species increases (Fig. 3b, SI Note 4).

### 3.4. Synteny and chromosomal evolution

Cytogenetic karyotyping (SI Note 1) revealed a difference of six chromosome pairs between *T. fasciculata* (2*n* = 50) and *T. leiboldiana* (2*n* = 38), which is atypical in this largely homoploid clade with generally constant karyotype (Gitaí et al., 2014; Brown and Gilmartin, 1989). To investigate orthology and synteny, we inferred orthogroups between protein sequences of *Ananas comosus* (Ming et al., 2015) (pineapple), *T. fasciculata* and *T. leiboldiana* using Orthofinder (Emms and Kelly, 2019). This resulted in 21,045 (78 %), 26,325 (87.5 %) and 23,584 (75 %) gene models assigned to orthogroups respectively, of which 10,021 were single-copy orthologs between all three species (Table S4).

Syntenic blocks were then defined across all three assemblies using Genespace (Lovell et al., 2022) (Fig. 3c). Remarkably, the three-way synteny analysis between *A. comosus, T. fasciculata* and *T. leiboldiana* show higher synteny between *T. fasciculata* and *A. comosus* than between the two *Tillandsia* genomes, which could be explained by *T. leiboldiana’*s diverged karyotype. While the difference in karyotype could have arisen from chromosomal loss in *T. leiboldiana*, our Genespace analysis reveals conserved synteny between the two *Tillandsia* assemblies without major orphan regions in *T. leiboldiana*. This is consistent with a scenario of chromosomal fusion, rather than loss. We found clear evidence of such a fusion on scaffold 14 in *T. leiboldiana* (Fig. 3c, Fig. S4a), which was confirmed with in-depth analyses of potential breakpoints (SI Note 5). However, chromosomal rearrangements are not limited to fusions, since we also detected two major reciprocal translocations (Fig. 3c, hereafter referred as Translocations 1 and 2, Fig. S4b-c).

### 3.5. Gene family evolution

6,261 genes in *T. fasciculata* and 4,693 genes in *T. leiboldiana* were assigned to non-unique gene families with multiple gene copies in at least one species, after correcting gene family sizes (Table S4). On average, the multicopy gene family size is 1.3x larger in *T. fasciculata* than in *T. leiboldiana* (Mann Whitney U, p-value: 8.8E-16, Fig. 4a).

**Figure 4:**
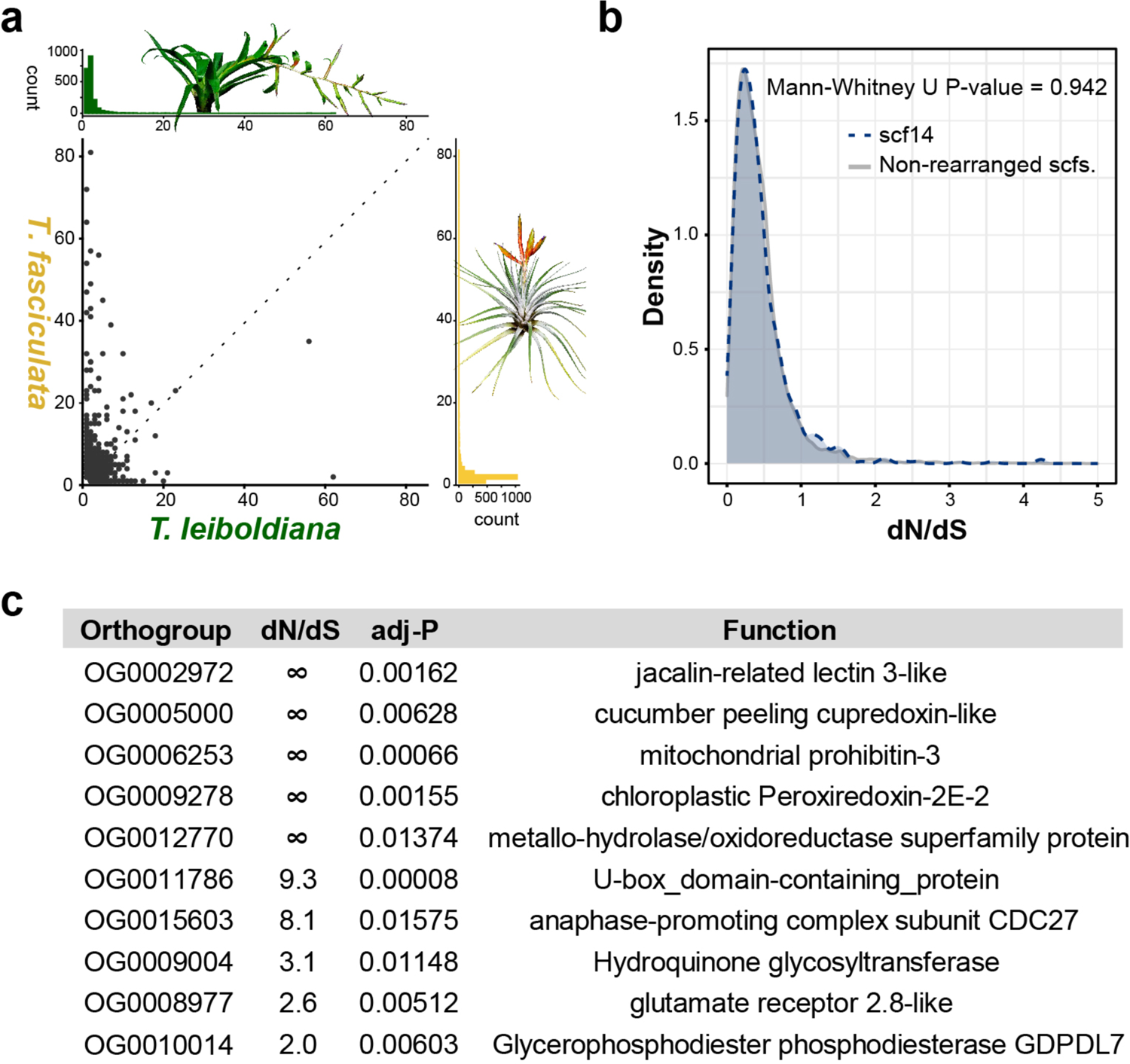
**a)** Scatterplot: composition of per-species gene counts among orthogroups. Upper histogram: distribution of per-orthogroup gene count in T. fasciculata. Lower histogram: distribution of per-orthogroup gene count in T. leiboldiana. **b)** Distribution of d_N_/d_S_ values of one-to-one orthologs across non-rearranged scaffolds (grey profile) and scaffold 14 in T. leiboldiana (blue profile), which is the result of a fusion. **c)** Single-copy orthogroups with significant d_N_/d_S_ values and their functions. Three uncharacterized genes that are excluded here are detailed in Table S6. Infinite d_N_/d_S_ values correspond to genes with d_S_=0 (no synonymous substitutions), an expected situation considering the low divergence of the two species. Further explanation about the biological significance of these functions can be found in SI Note 7.

To investigate the role of expanded gene families in CAM evolution, we combined gene ontology (GO) enrichment tests on multicopy orthogroups (SI Note 6) with a targeted search of known genes involved in the CAM pathway. This highlighted 25 multi-copy gene families with functions putatively related to CAM (Table S5), of which 17 have expanded in *T. fasciculata* and eight in *T. leiboldiana.* Families expanded in *T. fasciculata* included a malate dehydrogenase (MDH) and a β-carbonic anhydrase (CA), which are putatively involved in the carbon fixation module of CAM photosynthesis, and subunits of the two vacuolar pumps (V-ATPase and V-PPiase) known to energise the night-time transport of malate in pineapple (McRae et al., 2002) (Table S5). Additionally, two enolase families (members of the glycolysis pathway), a vacuolar acid invertase putatively involved in day-time soluble sugar accumulation in the vacuole (McRae et al., 2002; Holtum et al., 2005), and a pyrophosphate-dependent phosphofructokinase associated with night-time conversion of soluble sugars through the glycolysis to PEP (Carnal and Black, 1989) were expanded in *T. fasciculata*. Two subunits of succinate dehydrogenase, a member of the tricarboxylic acid cycle and the electron transport chain which also plays a role in stomatal opening regulation, a relevant aspect of CAM photosynthesis (Araújo et al., 2011), were also expanded in *T. fasciculata*. XAP5 CIRCADIAN TIMEKEEPER (XCT), a regulator of circadian rhythm and disease resistance (Liu et al., 2022) which was previously identified as undergoing rapid gene family evolution in *Tillandsia* (De La Harpe et al., 2020) is also expanded in *T. fasciculata*. Gene families expanded in *T. leiboldiana* contained three members of the glycolysis, two aquaporins, a member of the tricarboxylic acid cycle and a regulator of stomatal opening (Table S5).

### 3.6. Adaptive sequence evolution

Adaptive sequence evolution was evaluated in 9,077 one-to-one orthologous gene pairs using the non-synonymous to synonymous substitution ratio (ω = d_N_/d_S_). Little among-scaffold variation in d_N_/d_S_ was observed, with per-scaffold median d_N_/d_S_ values ranging from 0.32 to 0.39 in *T. fasciculata* and 0.31 to 0.4 in *T. leiboldiana* (Fig. S5a). Regions of large chromosomal rearrangement such as the fused scaffold 14 in *T. leiboldiana* do not exhibit strong signatures of fast coding sequence evolution (Fig. 4b), though for Translocation 1, d_N_/d_S_ values are slightly, yet significantly, lower for scaffold 13 in *T. fasciculata* and scaffold 19 in *T. leiboldiana* (Fig. S5b, SI Note 5).

Among the 9,077 orthologous gene pairs, 13 candidates (0.21%) exhibit a significant d_N_/d_S_ > 1 (adjusted p-value < 0.05, Fig. 4c, Table S6, SI Note 7). Notably, we recover a significant signal in a type B glycerophosphodiester phosphodiesterase (GDPDL-7). GDPDL’s are involved in cell wall cellulose accumulation and pectin linking, and play a role in trichome development (Hayashi et al., 2008), a main trait differentiating the two species and more broadly, green and grey *Tillandsia*. Additionally, GDPDL-7 may be involved in response to drought and salt stress (Cheng et al., 2011). A glutamate receptor (GLR) 2.8-like also exhibits a significant d_N_/d_S_ > 1. By mediating Ca^2+^ fluxes, GLRs act as signalling proteins and mediate a number of physiological and developmental processes in plants (Weiland et al., 2015), including stomatal movement (Kong et al., 2016). Although it is associated with drought-stress response in *Medicago trunculata* (Philippe et al., 2019), the specific function of GLR2.8 still remains unknown.

### 3.7. Gene expression analyses

To study gene expression differences linked to distinct photosynthetic phenotypes, we performed a time-series RNA-seq experiment using six plants of each species (Table S1, SI Note 8), sampled every four hours in a 24-hour period. We recovered 907 genes with a differential temporal expression (DE) profile between *T. fasciculata* and *T. leiboldiana*. Among them are 46 known CAM-related genes and 22 genes associated with starch metabolism and glycolysis/gluconeogenesis (Fig. S6). GO term enrichment of the 907 DE genes revealed many CAM-related functions such as malate and oxaloacetate transport, circadian rhythm, light response, water and proton pumps, sucrose and maltose transport and starch metabolism (Table S7; Fig. 5a). While none of the candidate genes for adaptive sequence evolution recovered in this study were differentially expressed, nine of 22 genes reported by De La Harpe et al. 2020 as candidates for adaptive sequence evolution during transitions to constitutive CAM in the wider context of the genus *Tillandsia* were also differentially expressed in this study (Table S7).

**Figure 5:**
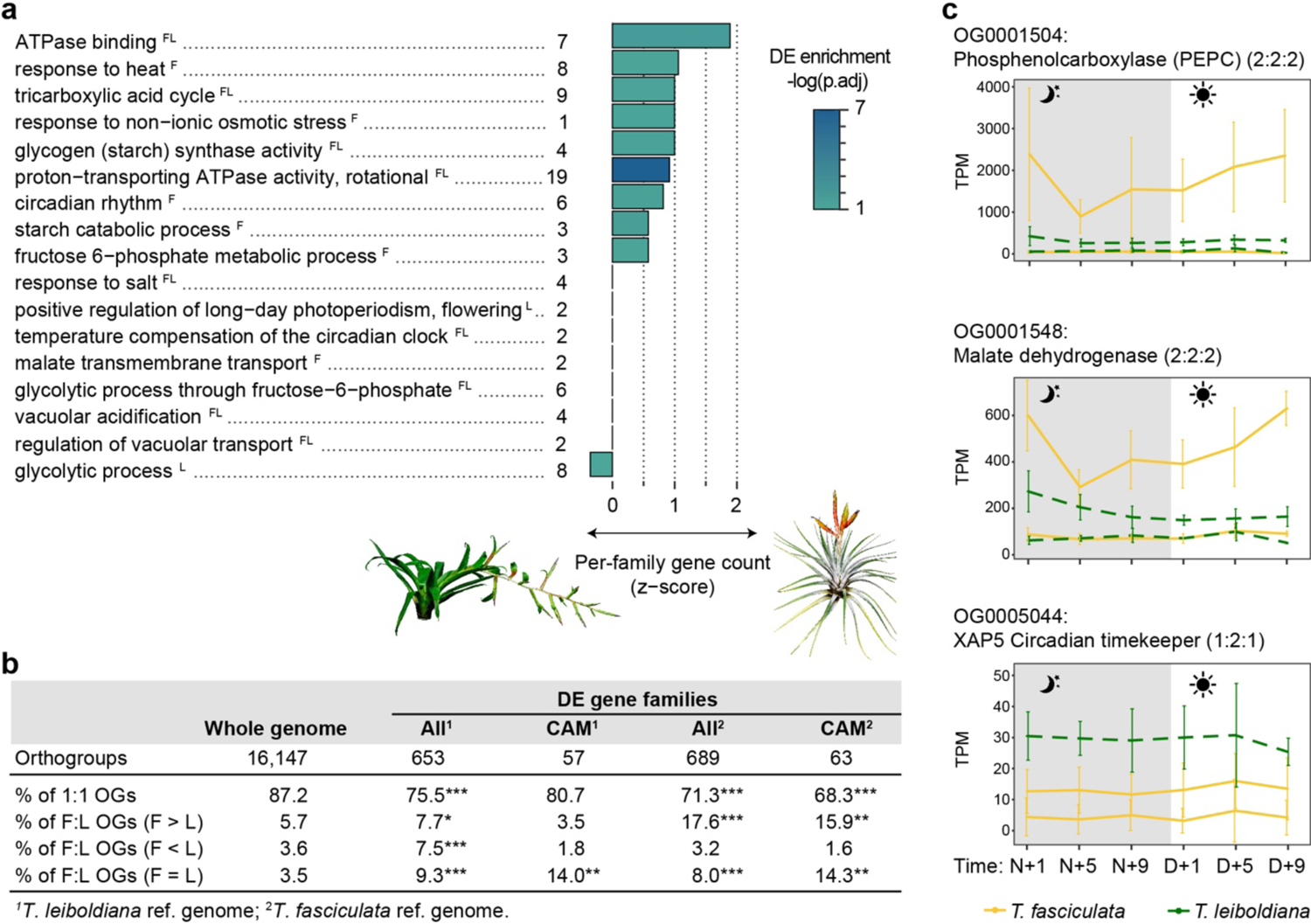
**a)** CAM-related enriched GO terms among differentially expressed (DE) genes between T. fasciculata and T. leiboldiana. The genome in which a GO term has been found to be enriched among DE genes is shown by ^F^ and ^L^ for T. fasciculata and T. leiboldiana, respectively. The family size difference for the underlying orthogroups is represented as a Z-score: a negative score indicates a tendency towards gene families with larger size in T. leiboldiana than in T. fasciculata, and vice versa. The p-value displayed represents the significance of the GO-term enrichment among DE genes in T. fasciculata unless the term was only enriched in T. leiboldiana. The number of DE genes underlying each function is shown next to the GO-term name. **b)** Composition of gene families by relative size between T. fasciculata (F) and T. leiboldiana (L) for three subsets of gene families (whole genome, DE, and CAM-related DE). Species-specific orthogroups are not included in this analysis. A chi-square test of independence was applied to test the significance of composition changes in 2×2 contingency tables for each category when testing the entire DE gene subset. For CAM DE genes, the Fisher’s exact test was applied. Significant p-values of both tests are reported as: *0.05-0.01, **0.01-0.0001, ***0.0001-0. **c)** Expression profiles in a 24-hour period of exemplary CAM-related gene families (PEPC, MDH and XCT) displayed at the orthogroup level. For each gene copy and timepoint, the average read count (in Transcripts Per Million, TPM), and the standard deviation across accessions are displayed. Read counts of each ortholog are obtained by mapping conspecific accessions to their conspecific reference genome. We show two families with older duplications preceding the split of T. fasciculata and T. leiboldiana (PEPC and MDH) and one gene family with a recent duplication in T. fasciculata (XCT).

Core CAM enzymes phosphoenolpyruvate carboxylase (PEPC) and phosphoenolpyruvate carboxylase kinase (PEPC kinase, PPCK) display clear temporal expression cycling in *T. fasciculata* (Fig. 5c, S5). PPCK also shows a night-time increase in expression in *T. leiboldiana* (Fig. S7), albeit with a milder temporal effect, a phenomenon that has been documented before in C3-assigned *Tillandsia* (De La Harpe et al., 2020) and also in other C3-like species belonging to CAM- and C4-evolving lineages (Heyduk et al., 2019b, 2019a). Clustering analysis distributed DE genes across seven clusters with sizes ranging from 209 to 38 genes (Table S7). CAM-related genes were distributed across six of seven clusters, highlighting the diversity of expression profiles associated with CAM (Fig. S8). While core CAM genes (see Fig. 6) are mainly found in cluster 5, we find malate transporters in cluster 1, circadian regulators in clusters 2 and 3, sugar transport in clusters 3 and 6, and vacuolar transport regulators in clusters 2, 4 and 6. Cluster 7, though not containing any core CAM candidate genes, was enriched for salt and heat stress response and contains a mitochondrial isocitrate dehydrogenase, which has been proposed as an alternative carbon fixator in CAM plants (Tay et al., 2021; Töpfer et al., 2020).

**Figure 6:**
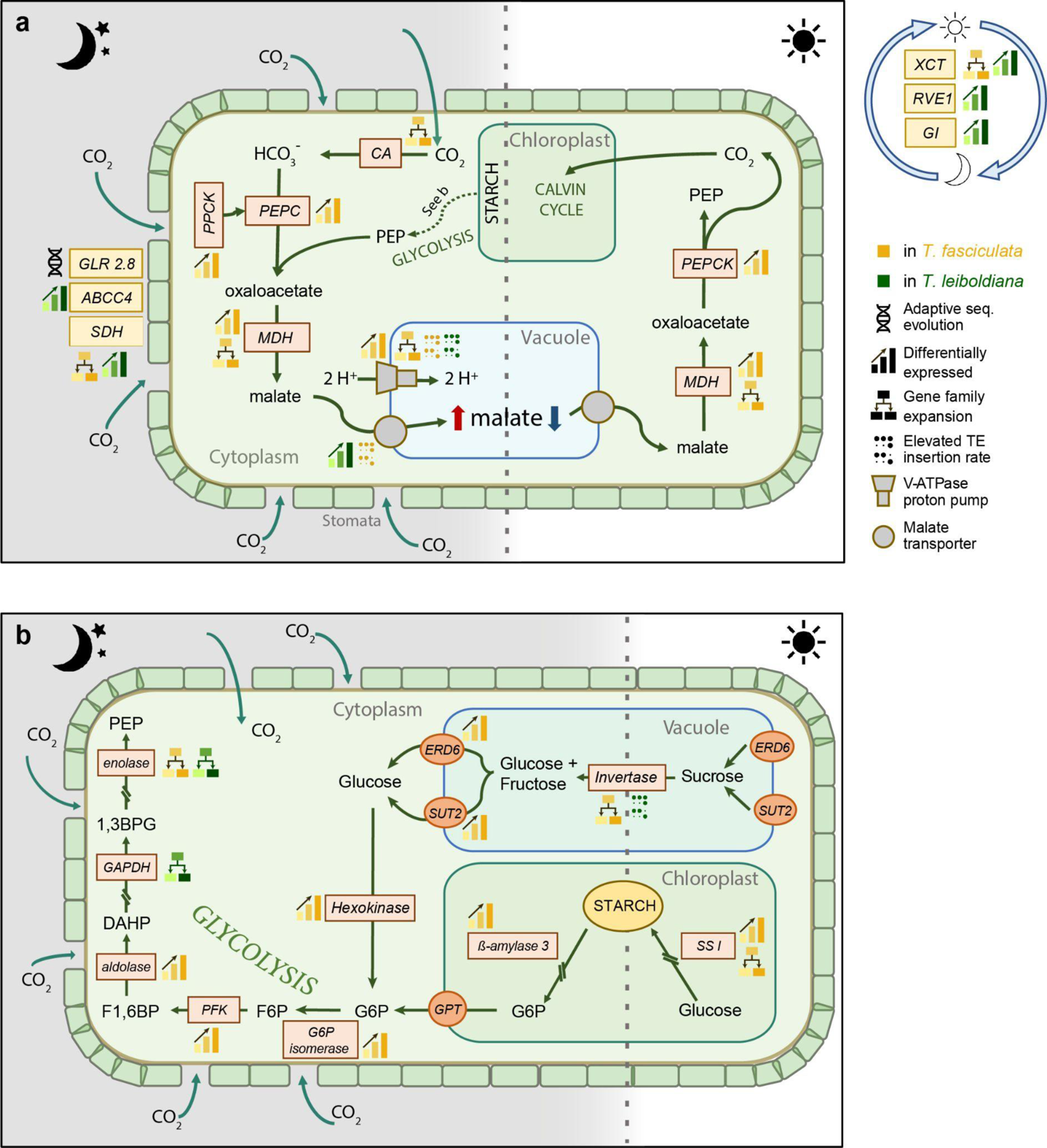
Pathway of Crassulacean Acid Metabolism (CAM), highlighting underlying genes detected in this study as differentially expressed, with gene family expansion, with signature of adaptive sequence evolution or elevated TE insertion counts. The colour of the process indicates the species showing diel or elevated expression, increased gene family size or increased TE insertion rate. Enzymes are shown in coloured boxes, while pathway products are shown in bold outside boxes. Enzymatic members of CAM metabolism pathways are shown in orange boxes, while stomatal and circadian regulators are highlighted in yellow. Stomatal regulators are shown on the left outside the cell and circadian regulators on the right. **a)** CO_2_ is absorbed at night and first converted to HCO_3_^-^ by carbonic anhydrase (CA). Then, it is converted to malate by carboxylating phosphoenolpyruvate (PEP), a key component of the glycolysis. In a first step, PEP carboxylase (PEPC) converts PEP to oxaloacetate, after being activated by PEPC kinase (PPCK). In a second step, malate dehydrogenase (MDH) converts oxaloacetate to malate. Malate is then transported into the vacuole by two possible transporters, either a tonoplast dicarboxylate transporter or an aluminium-activated malate transporter, which are assisted by V-ATPase proton pumps. During the day, the accumulated malate becomes the main source of CO_2_ for photosynthesis. This allows the stomata to remain closed, which greatly enhances the water use efficiency of the plant. Malate is again transported out of the vacuole and reconverted to oxaloacetate by MDH, and then decarboxylated to PEP and CO_2_ by PEP carboxykinase (PEPCK). The CO_2_ will cycle through the Calvin cycle and generate sugars. Abbreviations: GLR2.8 – Glutamate receptor 2.8, ABCC4 - ABC transporter C 4, SDH – Succinate Dehydrogenase, XCT – XAP5 CIRCADIAN TIMEKEEPER, GI – protein GIGANTEA, RVE1 – REVEILLE 1. **b)** Glycolysis, transitory starch, and sugar metabolism are tightly linked with the core CAM pathway as providers of starting materials such as PEP. During the day, CAM plants can store starch in the chloroplast and hexoses in the vacuole. In Bromeliaceae, the relative importance of soluble sugars versus starch as a source for PEP is variable across species^56^. At night, the stored starch and/or sugars are transported into the cytoplasm, converted to glucose or fructose, and broken down via the glycolysis to PEP. Abbreviations: G6P - glucose-6-phosphate; PGM – phosphoglucomutase; GPT - glucose-6-phosphate/phosphate translocator; PFK – phosphofructokinase; GAPDH - Glyceraldehyde-3-phosphate dehydrogenase; F6P – fructose-6-phosphate; F1,6BP – fructose-1,6-biphosphate; SSI – starch synthase I; 1,3BPG - 1,3-Bisphosphoglyceric acid; PEP – Phosphoenolpyruvate; SUT2 – Sucrose transporter 2; ERD6 - EARLY RESPONSE TO DEHYDRATION 6. For a detailed description and accompanying per-gene expression profiles, see Fig. S6 and SI Note 10.

The expression curves of the respective clusters (Fig. S8) demonstrate a complex web of expression changes between photosynthetic phenotypes. The most common expression change pattern among CAM-related genes is an overall increase in expression in the strong CAM plant (*T. fasciculata*), paired with increased diel cycling peaking in the early night (clusters 2, 5 and 6). This involved members of the night-time carbon fixating module of CAM such as PEPCK, PPCK and MDH, enzymes involved in malate transport as V-ATPase and several glycolysis enzymes such as glucose-6-phosphate isomerase, aldolase, Ppi-dependent phosphofructokinase (PFK) and enolase (Fig. 6, S6). Enzymes of both soluble sugar transport (SUT2, ERD6, cluster 6) and starch metabolism (Starch synthase I, α- and β-amylase and glucose-6-phosphate/phosphate translocator (GPT), cluster 5 and 6) are overall up-regulated and show cycling expression curves in *T. fasciculata* compared to *T. leiboldiana,* all showing highest activity in the late day. While the increased night-time expression of Ppi-dependent PFK suggests a primary role of soluble sugars as night-time source for PEP (Carnal and Black, 1989) in CAM *Tillandsia*, the simultaneously cycling expression patterns in starch metabolic enzymes point also at transitory starch as a potential source. A handful of CAM-related genes show increased expression in *T. leiboldiana*: aluminium-activated malate transporter (ALMT), secondary vacuolar proton pump AVP1, which also displays a phase shift peaking later in the night in *T. fasciculata*, and three circadian clock regulators (LHY, GI and RVE1, cluster 1), which all show similar but reduced cycling patterns in *T. fasciculata* compared to *T. leiboldiana*. We also see a phase shift in succinate dehydrogenase peaking earlier in the night in *T fasciculata* versus *T. leiboldiana*’s early morning peak.

Overall, most CAM-related DE gene expression profiles align with the view that *T. fasciculata* is a constitutive CAM plant while *T. leiboldiana* performs a C3-like metabolism in normal conditions, though showing signs of very early CAM evolution. The difference in metabolism between both plants seems to be largely attained through regulatory rewiring of functional enzymes.

### 3.8. Circadian clock-related motif enrichment in promoter sequences

We calculated the per-kb frequency of four known circadian clock-related motifs in the 2-kb upstream regions of identified DE genes, to further understand the role of circadian clock regulation in this set. We contrasted the frequencies of each motif in the set of DE genes against their frequencies in upstream regions of non-DE genes, and found that the Evening Element (EE) and CCA1-binding site (CBS) were the most enriched in this set with a frequency increase of 19% and 18 % respectively (Table S8). The difference in median per-promotor count of these motifs was however not statistically significant. Among co-expression clusters, the changes in motif frequency compared to non-DE genes varied greatly. We find a significant increase in motif frequency in cluster 1 for the G-Box motif (82 % increase), in cluster 3 for EE (207 %) and in cluster 7 for CBS (43 %).

We performed the same analysis on a set of *T. leiboldiana* genes that were temporally differentially expressed (see SI Note 10). The upstream regions of these genes also showed a small but not statistically significant increase of 16 % and 10 % in EE and CBS frequency respectively. The enrichment of circadian-clock related motifs in promotor regions of DE genes shows similarities between the two species, with comparable rates of frequency change for each specific element, though they are slightly larger in *T. fasciculata*.

When comparing the composition of circadian motifs in the upstream regions of core CAM genes and their homologs between species, we find a large diversity of motif presence among genes (Table S9), yet homologs between species tend to share the same motifs. No circadian motif appears to be present in any homolog of PEPC except for a copy in *T. leiboldiana* which was not differentially expressed between species. On the other hand, PPCK, an important regulator of PEPC, contains several circadian motifs, and shows marked differences in its composition compared to its non-DE homolog (PPCK1). The DE copy of PPCK misses two G-Box sites compared to its homolog in *T. leiboldiana*.

### 3.9. Genomic features of DE and CAM-related genes

#### 3.9.1. Genomic distribution of DE genes is not associated with rearranged regions

Differentially expressed genes are present on all major scaffolds of both genome assemblies and the total number of DE genes per scaffold is positively correlated with the scaffold size (Kendall’s correlation coefficient: 0.365 in *T. fasciculata* and 0.453 in *T. leiboldiana*, p-values < 0.015). Rearranged scaffolds in *T. leiboldiana* do not show a deviation in DE counts from other scaffolds relative to their size (Fig. S9). The density of DE genes is slightly higher in *T. fasciculata* than in *T. leiboldiana* (1.47 vs. 0.93 DE genes per 1-kb window on average). On the other hand, the average proportion of genes that are DE per 1-kb window is higher in *T. leiboldiana* (3.3 %) than in *T. fasciculata* (2.9 %), indicating that DE genes are more often located in gene-sparse regions in *T. leiboldiana* (Fig. S10).

#### 3.9.2. Differentially expressed genes belong more often to multi-copy orthogroups

To investigate the consequences of gene family evolution on gene expression, we tested whether the proportion of multi-copy orthogroups underlying DE genes was significantly elevated to that of the whole-genome set of orthogroups in both species (Fig. 5b, SI Note 9). The 907 DE genes in *T. fasciculata* are found in 738 orthogroups (hereafter called DE orthogroups) containing a total of 2,141 and 910 genes in *T. fasciculata* and *T. leiboldian*a, respectively. Genes from multi-copy orthogroups are more likely to be differentially expressed: while multi-copy orthogroups account for 24 % of all orthogroups in the genome, they represent 31 % of DE orthogroups. This difference is primarily explained by a 3.2x larger proportion of multi-copy orthogroups with a larger family size in *T. fasciculata* than in *T. leiboldiana* in the DE subset compared to the whole genome (Chi-square P = 1.59e^-66^).

Reciprocally, the DE analysis on the *T. leiboldiana* genome (SI Note 9) resulted in 836 DE genes belonging to 714 orthogroups, of which 489 overlap with the DE orthogroups resulting from the analysis on the *T. fasciculata* genome. As in the analysis on the *T. fasciculata* genome, we find that orthogroups with a larger family size in *T. leiboldiana* are enriched among DE genes. Additionally, both analyses point at a significant enrichment for multi-copy orthogroups with equal family sizes in both species, suggesting that also older duplications preceding the split of *T. fasciculata* and *T. leiboldiana* play a role in day-night regulatory evolution. This highlights the importance not only of novel, but also ancient variation in fuelling trait evolution in *Tillandsia*.

Multi-copy gene families are also enriched in a restricted subset of DE orthogroups related to CAM, starch metabolism and gluconeogenesis (68 genes in 67 orthogroups), especially gene families with equal copy number and with copy number expansion in *T. fasciculata* (Fig. 5b, Table S5). Importantly, expanded gene families in *T. leiboldiana* are not significantly enriched in this functional subset, showing that while the full set of DE orthogroups exhibit increased gene family dynamics in both lineages, CAM-related gene family expansion is only associated with *T. fasciculata* (constitutive CAM). This pattern is also reflected on the GO-term level, where enriched CAM-related biological functions appear disproportionally associated with gene family expansion in *T. fasciculata* (nine functions) than in *T. leiboldiana* (one function). Functions associated with V-ATPase proton pumps especially tend to have larger gene family size in *T. fasciculata* than in *T. leiboldiana* (ATPase binding, proton-transporting ATPase activity).

CAM-related expanded gene families often show one highly expressed copy that performs diel cycling, while the other copies are lowly or not expressed (e.g. SDH, glyceraldehyde-3-phosphate dehydrogenase (GAPDH), V-ATPase subunit H, SI Note 10), however, several gene families show diel and/or elevated expression in *T. fasciculata* in two or more gene copies (starch synthase, Ppi-dependent PFK and enolase, SI Note 10). Both copies of XCT are expressed in *T. fasciculata,* though showing no diel cycling or increased expression compared to *T. leiboldiana* (Fig. 5c), making its role in CAM photosynthesis unclear. V-ATPase subunit H has eight copies in *T. fasciculata* and three in *T. leiboldiana*. While in both species only one copy is highly expressed (with diel cycling peaking at night in *T. fasciculata*), the copies implemented for elevated expression in either species are not each other’s homologs (SI Note 10), suggesting that different copies are recruited for the distinct photosynthetic phenotypes of these species. The other copies are lowly or not expressed. A candidate gene family with larger gene family size in *T. leiboldiana* is a probable aquaporin PIP2-6 (OG0005047, Fig. S11), which is involved in water regulation and follows a diel pattern in pineapple (*A. comosus*) (Zhu and Ming, 2019). While lowly expressed in *T. fasciculata*, one of the two gene copies shows strong diel expression in *T. leiboldiana*, with highest expression in the early night. This is another indication that an early, latent CAM cycle may be present in *T. leiboldiana*.

CAM-related gene families with duplications preceding the divergence of *T. leiboldiana* and *T. fasciculata* include a malate dehydrogenase (MDH) with two copies in both species, where only one copy is highly expressed and cycling in *T. fasciculata,* and the core CAM enzyme Phosphoenolpyruvate carboxylase (PEPC), which shares an ancient duplication among monocots (Deng et al., 2016) (Fig. 5c). The widely varying expression patterns of multicopy DE CAM-related families suggest a variety of mechanisms possibly contributing to CAM regulatory evolution: dosage changes (“more of the same”), subfunctionalization and neofunctionalization.

#### 3.9.3. Differentially expressed genes have more TE insertions

To investigate whether transposable element (TE) activity and differential gene expression are associated in *Tillandsia*, we tested whether TE insertions in introns and the 3-kb upstream regions of genes are significantly enriched in DE genes in both species. Both the presence of one or more TE insertions in a gene, as well as the average number of TE insertions per gene is higher across all genes of *T. leiboldiana* compared to the *T. fasciculata* gene set, which was expected given its larger proportion of repetitive content (See Results section 3.2.). While in both genomes, the proportion of DE genes with one or more TE insertions is not significantly different to that of the full gene set, the average number of TE insertions per gene is significantly higher in DE compared to non-DE genes (Table 1).

**Table 1:** Statistical test results on TE insertions in DE versus non-DE genes, and in DE genes previously described as underlying CAM, glycolysis, or starch metabolism, versus all other genes. TE insertions were counted in intronic + 3-kb upstream regions, but insertions in introns only are also shown. P-value: *>0.05, **>0.01, ***>10-3.

| Total number of genes with one or more TE insertions |  |  |  |  |  |  |
| --- | --- | --- | --- | --- | --- | --- |
|  | *T. fasciculata* |  |  | *T. leiboldiana* |  |  |
|  | All genes | DE genes | CAM DE genes | All genes | DE genes | CAM DE genes |
| All TEs (intronic only) | 15,844 (50 %) | 473 (52 %) | 28 (41.2 %) | 10,348 (51 %) | 387 (54 %) | 21 (36.2 %)* |
| All TEs | 22,969 (90 %) | 725 (91 %) | 58 (85 %) | 18,512 (91 %) | 665 (92 %) | 50 (86 %) |
| DNA transposons | 16,857 (66 %) | 556 (70 %) | 45 (66 %) | 14,463 (71 %) | 547 (76 %)** | 42 (72 %) |
| Helitrons | 17,388 (69 %) | 561 (70 %) | 47 (69 %) | 14,701 (72 %) | 530 (74 %) | 38 (65 %) |
| LTR-Copia | 11,147 (44 %) | 353 (44 %) | 28 (41 %) | 10,487 (51 %) | 394 (55 %) | 31 (53 %) |
| LTR-Gypsy | 10,365 (41 %) | 341 (43 %) | 22 (32 %) | 5,399 (26 %) | 199 (28 %) | 10 (17 %) |
| Average and (median) TE insertion counts per gene |  |  |  |  |  |  |
|  | *T. fasciculata* |  |  | *T. leiboldiana* |  |  |
|  | non-DE genes | DE genes | CAM genes | non-DE genes | DE genes | CAM genes |
| All TEs (intronic only) | 2.90 (1) | 3.71 (1)* | 2.618 (0) | 3.24 (1) | 3.96 (1)** | 2.862 (0) |
| All TEs | 6.79 (5) | 7.83 (6)** | 6.32 (5) | 7.62 (5) | 8.75 (6)*** | 7.36 (5) |
| DNA transposons | 2.15 (1) | 2.56 (2)*** | 2.03 (1) | 2.49 (2) | 2.97 (2)*** | 2.77 (2) |
| Helitrons | 1.97 (1) | 2.27 (2)*** | 2.12 (1) | 2.43 (2) | 2.72 (2)* | 2.12 (1) |
| LTR-Copia | 1.00 (0) | 0.95 (0) | 0.87 (0) | 1.22 (1) | 1.45 (1)** | 1.25 (1) |
| LTR-Gypsy | 0.92 (0) | 1.09 (0) | 0.76 (0) | 0.61 (0) | 0.65 (0) | 0.39 (0) |

On the other hand, TE insertion rates in DE genes related to CAM, starch metabolism and glycolysis / gluconeogenesis do not significantly differ from background rates in both genomes, though they are slightly reduced (Table 1). However, the proportion of CAM-related DE genes with an intronic TE insertion is larger in *T. fasciculata* (41.2 %) than in *T. leiboldiana* (36.2 %), despite *T. leiboldiana*‘s generally elevated intronic TE insertion rate. This pattern is not discernible when including TE insertions in the 3-kb upstream region of genes.

When studying genic TE insertions across four separate TE classes, we recover a similar trend among all categories as observed across all TEs. Insertion rates around genes are the highest for DNA transposons in both species, but the TE class that is most often present around a gene are Helitrons, which occur in 69 % and 72 % of genes in *T. fasciculata* and *T. leiboldiana* respectively (Table 1). This contrasts with the small proportion of the whole genome that is covered by Helitrons – only 5.86 % and 3.7 % in *T. fasciculata* and *T. leiboldiana* respectively (Table S3). LTRs, while covering the largest proportion of the genome in both species, are the least present and show the lowest insertion rates around genes in both species.

Nine DE genes related to CAM, starch, and gluconeogenesis display more than twice the number of TE insertions as the genome-wide average in *T. fasciculata*. This includes a V-type proton ATPase subunit H (vacuolar transport and acidification), an aluminium-activated malate transporter, an ABC transporter C family member 4 (stomatal opening and circadian rhythm) and a mitochondrial isocitrate dehydrogenase subunit (Table S10), which all had more TE insertions than their orthologs in *T. leiboldiana*. On the other hand, six DE genes of interest had elevated TE insertion rates in *T. leiboldiana,* including a vacuolar acid invertase (sugar metabolism) and a sugar transporter ERD6-like. Four of these genes showed high amounts of TE insertions in both genomes, such as a glucose-1-phosphate adenylyltransferase subunit (starch synthesis), a V-type proton ATPase subunit C, and circadian clock regulator GIGANTEA (Dalchau et al., 2011), which also plays a role in stomatal opening (Ando et al., 2013).

## 4. Discussion

The sources of variation fuelling trait evolution in rapid radiations have been a long-standing topic in evolutionary biology (Simpson, 1953), and our understanding of how complex traits such as CAM evolve repeatedly is still incomplete. By showcasing a broad range of photosynthetic phenotypes and repeated evolution of constitutive CAM, the subgenus *Tillandsia* provides an excellent opportunity to study CAM evolution. The recent divergence between members of *Tillandsia* allows to pinpoint the necessary evolutionary changes to evolve a constitutive CAM phenotype. By integrating comparative genomics using *de novo* assemblies and in-depth gene expression analyses of two closely related *Tillandsia* species representing one of the most distinct photosynthetic phenotypes within the clade, we found support for regulatory evolution and gene family expansion as major features of CAM evolution (Fig. 6).

Our metabolic analyses of night-time malate accumulation and gene expression analyses provide a much more detailed understanding of the photosynthetic phenotypes present in the subgenus than the previously reported carbon isotope measurements, which do not accurately reflect all stages of the CAM continuum. For example, weak CAM phenotypes have been reported in Bromeliaceae for species with carbon isotope ratios falling in the C3 range (−26.5) (Pierce et al., 2002), indicating that the photosynthetic metabolism of *T. leiboldiana* could be different from C3 *sensu stricto* as its carbon isotope measurements suggest. Our timewise malate measurements show distinct fluxes for *T. fasciculata* and *T. leiboldiana*, with constitutive accumulation of night-time malate in *T. fasciculata* which is absent in *T. leiboldiana*. On the other hand, *T. leiboldiana* displays CAM-like temporal expression profiles for certain enzymes, such as PEPC kinase and Aquaporin PIP-6, and shares circadian clock-related cis-elements in the promotor regions of CAM homologs with *T. fasciculata*. It has been suggested that repeated evolution of CAM (similar to C4) may be facilitated in lineages where C3 species already display increased or CAM-like expression of core genes (Heyduk et al., 2019a, 2019b; Kajala et al., 2012). However, while a CAM cycle is seemingly not being expressed in *T. leiboldiana* under normal circumstances, we cannot exclude that a latent CAM cycle could become activated under certain conditions, for example under drought stress. In that case, *T. leiboldiana* would rather be at the very early stages of CAM evolution than a “pre-adapted” C3 plant (De La Harpe et al., 2020). We hope that future studies will study the drought response of *T. leiboldiana* to better understand its exact position in the CAM continuum.

On the other hand, even if *T. leiboldiana* and potentially all subgenus *Tillandsia* species previously labelled as C3 represent in fact the early stages of the CAM continuum, our analyses show widely distinct photosynthetic phenotypes within the radiation which required divergent evolution. Therefore, while this study may underestimate the total number of evolutionary changes needed to establish constitutive CAM from C3 *sensu stricto*, it highlights the evolutionary drivers underlying the least understood section of the CAM continuum: from early CAM to constitutive CAM.

Differences between the two genomes related to CAM evolution can be primarily found on the regulatory level, with CAM-related genes showing temporal differential expression between species across a 24-hour period. These reveal a complex web of underlying expression changes, as they are distributed over all inferred co-expression clusters (Fig. S8). Together with the diversity of circadian clock-related motif composition in promoter sequences (Table S8), this finding emphasises the lack of a master regulator and a clear overall direction of expression changes underlying CAM (Wickell et al., 2021; Heyduk et al., 2022).

Gene family expansion has been previously observed in CAM lineages (Cai et al., 2015; Silvera et al., 2010) and suggested as a driver of CAM evolution (Silvera et al., 2014). We witnessed an increased number of genes belonging to multi-copy families in *T. fasciculata* than in *T. leiboldiana*, consistent with a net higher rate of gene duplication in this species than in *T. leiboldiana*, as previously reported by De La Harpe et al. (2020). Strikingly, both the total subset of differentially expressed genes and a more stringent group of CAM-related DE genes was significantly enriched for gene families that have expanded in *T. fasciculata* (constitutive CAM). CAM-related functions that were enriched in DE genes show a disproportionate bias towards gene family expansion in *T. fasciculata* (circadian rhythm, vacuolar ATPase activity, tricarboxylic acid cycle and starch metabolism) compared to *T. leiboldiana* (glycolysis) (Fig. 5a). Gene duplications, preceding the split of *T. fasciculata* and *T. leiboldiana* are also significantly associated with day-night expression differences in CAM-related genes (Fig. 5b), suggesting that older, already existing gene duplications may also be recruited in CAM evolution, alongside novel duplications.

The expression curves of DE multicopy gene families with a potential link to CAM reveal a multitude of expression behaviours (e.g. Fig. 5c), which supports that complex regulatory evolution on the transcriptional level underlies CAM evolution. Our findings suggest that gene family evolution played a significant role in modulating regulatory changes underlying the evolution towards constitutive CAM in *Tillandsia*. As gene family expansion leads to increased redundancy, selection on individual gene copies and their expression relaxes, facilitating the assimilation of a constitutively expressed CAM expression profile (Ohno, 1970).

Another potential driver of trait evolution is TE insertion, though its role in CAM evolution in *Tillandsia* remains unclear. TE insertions are overall less common in CAM-related DE genes compared to all genes in both genomes, suggesting a selection pressure against TE insertions around these genes. However, the proportion of CAM-related DE genes with intronic TE insertions is greater in *T. fasciculata* than in *T. leiboldiana,* despite the overall higher genic TE insertion rate in *T. leiboldiana*. This suggests that the pressure to maintain CAM-related genes TE-free is reduced in the constitutive CAM lineage relative to *T. leiboldiana*. We detect nine and six CAM-related DE genes with more than twice the whole-genome average TE insertion count in *T. fasciculata* and *T. leiboldiana* respectively, of which four are single-copy orthologs shared between species. The high degree of sharedness of TE-rich DE genes between species rather suggests that TE insertions are not a major driver of CAM-related gene expression changes. In fact, genes with exceptionally high insertion rates in *T. fasciculata* tend to show reduced expression (ALMT, V-ATPase subunit H copy Tfasc_v1.24696, ABCC4, SI Note 10). Instead, the larger proportion of CAM-related DE genes with one or more TE insertions in *T. fasciculata* may be a consequence of higher rates of gene family expansion and eventual pseudogenization of redundant copies.

Candidate genes under positive selection underlie a broad array of functions, but had no immediate link to CAM photosynthesis. While the study of adaptive sequence evolution would greatly benefit from a broader sampling across *Tillandsia*, the lack of overlap between regulatory and adaptive sequence evolution is in line with previously proposed mechanisms of CAM evolution largely relying on regulatory changes in other systems (Deng et al., 2016). A small number of cases of convergent and adaptive sequence evolution between distantly related CAM and C3 species have been described (Yang et al., 2017), though no overlap was found between convergence in expression and sequence evolution. Our study suggests that while on larger evolutionary scales adaptive sequence evolution may play an important role, distinct photosynthetic phenotypes between closely related species may be achieved primarily with gene expression changes (but see SI Note 12), or may be especially relevant between the transition of C3 *sensu stricto* to the CAM continuum.

Though we observe a karyotype difference of six chromosome pairs between *T. fasciculata* and *T. leiboldiana* and we identified one fusion in the *T. leiboldiana* assembly, along with two reciprocal translocations, we did not find detectable consequences of large-scale rearrangements for either functional diversification or adaptation in *Tillandsia*, unlike other studies (Davey et al., 2016; Cicconardi et al., 2021) (Fig. 4b, S4, S6, but see SI Note 5 and 14). However, due to the remaining fragmentation of the *T. leiboldiana* genome, it is likely that we were not able to describe all rearrangements, and we hope that future endeavours will improve the genome assembly and make a more in-depth study of the role of large-scale rearrangements in the evolution of species barriers and/or the evolution of other key innovation traits in *Tillandsia possible*.

Our analyses reveal genomic changes of all scales between two members of an adaptive radiation representing a recent shift to constitutive CAM. However, in this recent shift between closely related species, differences in photosynthetic metabolism are brought about largely by temporal expression changes enabled by both existing and *de novo* gene duplication, rather than adaptive sequence evolution of existing gene copies, which may play a role at later stages of divergence (but see SI Note 12). Large scale rearrangements observed so far seem unlinked from functional divergence, more likely affecting reproductive isolation (de Vos et al., 2020; Faria and Navarro, 2010), and need further study. Our findings support an important role for gene family expansion in generating novel variation that fuels the evolution of the CAM continuum.

The two *de novo* assemblies presented in this study are the first tillandsioid and fourth bromeliad genomes published so far. Despite both genomes exhibiting one of the highest TE contents reported to date for a non-polyploid plant species (Pedro et al., 2021), the joint use of long-read sequencing and chromatin conformation capture successfully led to highly contiguous assemblies with high-quality gene sets (SI Note 13). Along with other recently developed resources for Bromeliaceae (Yardeni et al., 2021; Liu et al., 2021), these genomes will be crucial in future investigations of this highly diverse and species-rich plant family, and in further studies of CAM evolution.

## 5. Materials & Methods

### 5.1. Flow cytometry and cytogenetic experiments

#### 5.1.1. Genome size measurements

Approximately 25 mg of fresh leaf material was co-chopped according to the chopping method of (Galbraith et al., 1983) together with an appropriate reference standard (*Solanum pseudocapsicum*, 1.295 pg/1C) (Temsch, 2010; Temsch et al., 2022) in Ottós I buffer (Otto et al., 1981). After filtration through a 30 µm nylon mesh (Saatilene Hitech, Sericol GmbH, Germany) and incubation with RNase A (0.15mg/ml, Sigma-Aldrich, USA) at 37°C, Ottós II buffer (Otto et al., 1981) including propidium iodide (PI, 50mg/L, AppliChem, Germany) was added. Staining took place in the refrigerator for at least one hour or up to over-night. Measurement was conducted on a CyFlow ML or a CyFlow Space flow cytometer (Partec/Sysmex, Germany) both equipped with a green laser (532nm, 100mW, Cobolt AB, Sweden). The fluorescence intensity (FI) of 10,000 particles were measured per preparation and the 1C-value calculation for each sample followed the equation: 1C_Obj_ = (FI peak mean_G1 Obj_ / FI peak mean_G1 Std_) × 1C_Std_

#### 5.1.2. Karyotyping

Actively growing root meristems of genome assembly accessions (see Table S1) were harvested and pre-treated with 8-hydroxyquinoline for 2 hours at room temperature and 2 hours at 4°C. The roots were then fixed in Carnoy’s fixative (3 : 1 ethanol : glacial acetic acid) for 24 hours at room temperature and stored –20 °C until use. Chromosome preparations were made after enzymatic digestion of fixed root meristems as described in (Jang and Weiss-Schneeweiss, 2015). Chromosomes and nuclei were stained with 2 ng/µl DAPI (4’,6-diamidino-2-2phenylindole) in Vectashield antifade medium (Vector Laboratories, Burlingame, CA, USA). Preparations were analysed with an Axiolmager M2 epifluorescent microscope (Carl Zeiss) and images were captured with a CCD camera using AxioVision 4.8 software (Carl Zeiss). Chromosome number was established based on analyses of several preparations and at least five intact chromosome spreads. Selected images were contrasted using Corel PhotoPaint X8 with only those functions that applied equally to all pixels of the image and were then used to prepare karyotypes.

### 5.2. Genome Assembly

#### 5.2.1. Plant material selection and sequencing

Genome assemblies were constructed from the plant material of one accession per species (see Table S1). The accessions were placed in a dark room for a week to minimise chloroplast activity and recruitment, after which the youngest leaves were collected and flash frozen with liquid nitrogen. High molecular weight extraction for ultra-long reads, SMRTbell library preparation and PacBio Sequel sequencing was performed by Dovetail Genomics^TM^ (now Cantata Bio). Dovetail Genomics^TM^ also prepared Chicago (Putnam et al., 2016) and Hi-C (Lieberman-Aiden et al., 2009) libraries which were sequenced as paired-end 150bp reads on an Illumina HiSeq X instrument. Additional DNA libraries were prepared for polishing purposes using Illumina’s TruSeq PCR-free kit, which were sequenced on a HiSeq2500 as paired-end 125 bp reads at the Vienna BioCenter Core Facilities (VBCF), Austria.

RNA-seq data of *T. fasciculata* used for gene annotation was sampled, sequenced, and analysed in De La Harpe et al. 2020 under SRA BioProject PRJNA649109. For gene annotation of *T. leiboldiana*, we made use of RNA-seq data obtained during a similar experiment, where plants were kept under greenhouse conditions and sampled every 12 hours in a 24-hour cycle. Importantly, while the *T. fasciculata* RNA-seq dataset contained three different genotypes, only clonal accessions were used in the *T. leiboldiana* experiment. For *T. leiboldiana*, total RNA was extracted using a QIAGEN RNeasy® Mini Kit, and poly-A capture was performed at the Vienna Biocenter Core Facilities (VBCF) using a NEBNext kit to produce a stranded mRNA library. This library was sequenced on a NovaSeq SP as 150 bp paired end reads.

For both species, sequencing data from different time points and accessions were merged into one file for the purpose of gene annotation. Before mapping, the data was quality-trimmed using AdapterRemoval (Schubert et al., 2016) with default options (--trimns, --trimqualities). We allowed for overlapping pairs to be collapsed into longer reads.

#### 5.2.2. First draft assembly and polishing

We constructed a draft assembly using long-read PacBio data with CANU v1.8 (Koren et al., 2017) for both species. To mitigate the effects of a relatively low average PacBio coverage (33x), we ran two rounds of read error correction with high sensitivity settings (corMhapSensitivity=high corMinCoverage=0 corOutCoverage=200) for *T. fasciculata.* Additionally, we applied high heterozygosity (correctedErrorRate=0.105) settings, since K-mer based analyses pointed at an elevated heterozygosity in this species (See SI Note 2), and memory optimization settings (corMhapFilterThreshold=0.0000000002 corMhapOptions=“--repeat-idf-scale 50” mhapMemory=60g mhapBlockSize=500).

Given that the coverage of *T. leiboldiana* PacBio averaged 40x, we limited error correction for this species to only one round. CANU was run with additional settings accommodating for high frequency repeats (ovlMerThreshold=500) and high sensitivity settings as mentioned above. To minimise the retention of heterozygous sequences as haplotigs in *T. fasciculata* (see SI Note 2), we reassigned allelic contigs using the pipeline Purge Haplotigs (Roach et al., 2018). Raw PacBio data was mapped to the draft assembly produced in the previous step with minimap2 (Li, 2018), before using the Purge Haplotigs pipeline.

Since the size of the *T. leiboldiana* draft assembly indicates, together with previous analyses, that this species is largely homozygous (SI Note 2), we did not include a PurgeHaplotigs step. However, we did make use of the higher average coverage of the *T. leiboldiana* PacBio data to polish the assembly with two rounds of PBMM v.1.0 and Arrow v2.3.3 (Pacific Biosciences).

#### 5.2.3. Scaffolding and final polishing

Scaffolding of both assemblies was performed in-house by Dovetail Genomics^TM^ using Chicago and Hi-C data and the HiRise scaffolding pipeline (Putnam et al., 2016). To increase base quality and correct indel errors, we ran additional rounds of polishing with high-coverage Illumina data (See above, section 2.1.) using Pilon v1.22 (Walker et al., 2014). The Illumina data was aligned to the scaffolded assembly using BWA-MEM (Li, 2013), and then Pilon was run on these alignments. We evaluated the result of each round using BUSCO v.3 (Waterhouse et al., 2018) with the Liliopsida odb9 library and proceeded with the best version. For *T. fasciculata*, polishing was performed twice, fixing SNPs and indels. We did not fix small structural variation in this genome due to the relatively low coverage (35x) of Illumina data. For *T. leiboldiana*, one round of polishing on all fixes (SNPs, indels and small structural variants) resulted in the highest BUSCO scores.

### 5.3. Annotation

#### 5.3.1. TE annotation and repeat masking

*De novo* TE annotation of both genome assemblies was performed with EDTA v.1.8.5 (Ou et al., 2019) with option –sensitive. To filter out genes that have been wrongly assigned as TEs, *A. comosus* (pineapple) coding sequences (Ming et al., 2015) were used in the final steps of EDTA. Using the species-specific TE library obtained from EDTA, we masked both genomes using RepeatMasker v.4.0.7 (Smit et al., 2013-2015). Importantly, we excluded all TE annotations marked as “unknown” for masking to prevent potentially genic regions flagged as TEs to be masked during annotation. The search engine was set to NCBI (-e ncbi) and simple and low-complexity repeats were left unmasked (-nolow). We produced both hard-masked and soft-masked (--xsmall) genomes.

#### 5.3.2. Transcriptome assembly

We constructed transcriptome assemblies for both species using Trinity *de novo* assembler v.2.4.8. (Grabherr et al., 2011) using default parameters starting from the raw mRNA-seq data. These were evaluated with BUSCO. Additionally, before feeding the transcriptome assemblies to the gene annotation pipeline, we ran a round of masking of interspersed repeats to avoid an overestimation of gene models due to the presence of active transposases in the RNA-seq data.

#### 5.3.3. Gene prediction and functional annotation

Gene models were constructed using a combination of BRAKER v.2.1.5 (Hoff et al., 2019) and MAKER2 v.2.31.11 (Campbell et al., 2014). Starting with BRAKER, we obtained splicing information from RNA-seq alignments to the masked genome as extrinsic evidence using the *bam2hints* script of AUGUSTUS v.3.3.3 (Stanke et al., 2008). A second source of extrinsic evidence for BRAKER were single-copy protein sequences predicted by BUSCO when run on the masked genomes in genome mode with option --long. Predictions made by BRAKER were evaluated with BUSCO and with RNA-seq alignments.

Subsequently, we built our final gene predictions using MAKER2. As evidence, we used (1) the gene models predicted by BRAKER, (2) a transcriptome assembly of each respective species (see above section 3.2.), (3) a protein sequence database containing proteins of *Ananas comosus comosus* (F153) (Ming et al., 2015) and *Ananas comosus bracteatus* (CB5) (Chen et al., 2019) and manually curated SwissProt proteins from monocot species (64,748 sequences in total) and (4) a GFF file of complex repeats obtained from the masked genome (see above section 3.1.) and an extended repeat library containing both the EDTA-produced *Tillandsia*-specific repeats and the monocot repeat library from RepBase (7,857 sequences in total). By only providing masking information of complex repeats and setting the model organism to “simple” in the repeat masking options, hard-masking in MAKER2 was limited to complex repeats while simple repeats were soft-masked, which makes these available for gene prediction. MAKER2 predicts genes both *ab initio* and based on the given evidence using AUGUSTUS.

We evaluated the resulting set of predicted gene models by mapping the RNA-seq data (section 2.1.) back to both the transcript and full gene model sequences and running BUSCO in transcriptome mode. We also calculated the proportion of masked content in these gene models to ascertain that MAKER2 had not predicted TEs as genes. A second run of MAKER, which included training AUGUSTUS based on the predicted models from the first round, resulted in lower BUSCO scores and was not further used. We functionally annotated the final set of gene models in Blast2Go v.5.2.5 (Götz et al., 2008) using the Viridiplantae database.

### 5.4. Inferring gene orthology

Orthology between gene models of *T. fasciculata*, *T. leiboldiana* and *Ananas comosus* was inferred using Orthofinder v.2.4.0 (Emms and Kelly, 2019). Protein sequences produced by MAKER2 of inferred gene models were used for *T. fasciculata* and *T. leiboldiana*. For *A. comosus*, the publicly available gene models of F153 were used. The full Orthofinder pipeline was run without additional settings. Counts per orthogroup and the individual genes belonging to each orthogroup were extracted from the output file Phylogenetic_Hierarchical_Orthogroups/N0.tsv.

Orthofinder was run a second time on gene models present only on main contigs (See Results). For each gene model, the longest isoform was selected, and gene models with protein sequences shorter than 40 amino acids were removed. This resulted in 27,024, 30,091 and 31,194 input sequences for *A. comosus*, *T. fasciculata* and *T. leiboldiana* respectively. Then, the steps mentioned above were repeated.

### 5.5. Gene model assessment and curation

Gene model sets were assessed and curated using several criteria. Gene models with annotations indicating a repetitive nature (transposons and viral sequences) together with all their orthologs were marked with “NO_ORTHOLOGY” in the GFF file and excluded from downstream analyses. Using the per-exon expression data obtained in our mRNA-seq experiment (see below, section 11) and information gathered on the length of the CDS and the presence / absence of a start and stop codon, we further classified our gene models into ROBUST and NOT-ROBUST categories. A gene model was considered ROBUST (i) if all exons are expressed or, (ii) if both start and stop codons are present and the CDS has a minimum length of 50 amino-acids.

### 5.6. Analysing TE class abundances

By rerunning EDTA with step --anno, we obtained TE abundances and detailed annotation of repetitive content for the whole assembly. Per-contig abundances of each class were calculated with a custom python script (available at https://github.com/cgrootcrego/Tillandsia_Genomes). Using this curated TE library, the assemblies were masked again with RepeatMasker for downstream analyses. The resulting TE class abundances reported by RepeatMasker were then compared between species and reported.

### 5.7. Spatial distribution of repetitive, genic and GC content

The spatial distribution of genes, transposable elements and GC content as shown in Fig. 3a, was analysed on a per-window basis, using windows of 1 Mb. Gene counts were quantified as the number of genes starting in every window, based on genes with assigned orthology, including both single and multicopy gene models. Repetitive content was measured as the proportion of masked bases in each window, stemming from the hard-masked assembly using the curated TE library. Per-window gene counts and proportion of repetitive bases was then visualised using the R package circlize (Gu et al., 2014). GC content was calculated as the proportion of G and C bases per 1 Mb windows. Correlation between genic, repetitive and GC content was calculated and tested for significance using the Kendall Rank Correlation Coefficient, after testing for normality using the Shapiro-Wilk test.

Repetitive, GC and gene content as shown in Fig. 3b was estimated directly from the soft-masked reference genomes using 100 kb non-overlapping sliding windows as described in (Leroy et al., 2021). TE content corresponds to the proportion of soft-masked positions per window. For the *Tillandsia* genomes, the curated TE library (see above, section 6.) was used as a basis for soft-masking in RepeatMasker. For *A*. *comosus*, a soft-masked version of the genome was obtained from NCBI (https://www.ncbi.nlm.nih.gov/datasets/genome/GCA_902162155.2/). As compared to the version of Leroy et al. 2021, this script was modified to estimate GC content in repetitive regions (soft-masked regions only). In addition to this, we estimated the genic fraction by considering the total number of genomic positions falling in genes based on the GFF files (feature = “gene”) divided by the size of the window (100 kb). This estimate was derived for the same window boundaries as used for GC and TE content to be able to compare all statistics. The relative per-window proportion of genic bases corresponding to non-robust genes (see above, Section 5) was also estimated by dividing the number of non-robust gene positions with the total number of gene positions.

### 5.8. Synteny between T. fasciculata and T. leiboldiana

Synteny was inferred with GENESPACE v.0.8.5 (Lovell et al., 2022), using orthology information obtained with Orthofinder of the gene models from *A. comosus*, *T. fasciculata* and *T. leiboldiana*. This provided a first, visual graphical to detect large-scale rearrangements. We used GENESPACE with default parameters, except that we generated the syntenic map (riparian plot) using minGenes2plot=200. Other methods have also been used to confirm the chromosomal rearrangements and to identify the genomic breakpoints more precisely (see SI Note 5).

### 5.9. Gene family evolution

#### 5.9.1. Family size correction

Gene counts per orthogroup were evaluated using per-gene mean coverage to detect co-assembled heterozygous gene sequences that may have escaped Purge Haplotigs in the assembly step. To do this, whole-genome Illumina reads of both species (See Methods, section 2.1.) were aligned to their respective assemblies using Bowtie2 v.2.4.4. (Langmead and Salzberg, 2012) with the very-sensitive-local option. Bowtie2 specifically assigns multi-mapping reads randomly, allowing the detection of artificial gene models thanks to a decreased overall coverage across the orthogroup, as reads from one biological copy would be randomly distributed over two or more locations in the genome. Per-base coverage in genic regions was calculated using *samtools depth* and a bed-file specifying all locations of orthologous genes. We then calculated the average coverage per orthologous gene.

The distribution of per-gene mean coverage in each species’ gene model set was then visualised using ggplot2 (Wickham, 2016) for different categories of genes: single-copy (only one gene model assigned to the orthogroup in the species investigated), multi-copy (more than one gene assigned to the orthogroup in the species investigated), ancestral single-copy (only one gene model assigned to the orthogroup in all species used in the orthology analysis), ancestral multi-copy (multiple gene model assigned to the orthogroup in all species used in the orthology analysis and the number of gene models assigned is equal across species), and unique multi-copy (more than one gene assigned to the orthogroup in the species investigated and no genes assigned to the orthogroup in other species). This revealed that, while most categories of genes had a unimodal distribution centred around the average coverage across the genome, multi-copy and unique multi-copy families showed a bimodal or expanded distribution, especially in *T. fasciculata* (Fig. S12). This points at the presence of genes with multiple alleles per gene in the annotation. Hence, gene count sizes per orthogroup and species were corrected by the ratio of the total coverage across all genes of one species in the orthogroup and the expected coverage, which was calculated as the product of the total number of genes in the orthogroup and the average coverage of single-copy genes in that species.

Size corrections were only applied on orthogroups containing multicopy genes. Plastid and mitochondrial genes were excluded from this analysis. We detected plastid genes with BLASTn against the *A. comosus* chloroplast sequence and the *Oryza* IRSGP-1 mitochondrial sequence. Additionally, all genes annotated as “ribosomal” were also excluded from the downstream gene family evolution analyses. Originally, 9,210 genes in *T. fasciculata* and 6,257 genes in *T. leiboldiana* were assigned to orthogroups with multiple gene copies in at least one species. After correcting orthogroup sizes by coverage, we retained 6,261 and 4,693 gene models, respectively (Table S4).

#### 5.9.2. Analysis of multicopy orthogroups

The distribution of gene counts per multicopy orthogroup was compared between *T. fasciculata* and *T. leiboldiana* with a non-parametric test (Mann-Whitney U). Using the log-ratio of per-species gene count, we investigated which gene families experienced large changes in gene count compared to the background (SI Note 6).

Functional characterization of multicopy families was done with a GO term enrichment analysis of the underlying genes using the Fisher’s exact test in TopGo (Alexa and Rahnenführer, 2009). Enrichment analyses were done on all genes belonging to multicopy orthogroups, on a subset of genes belonging to families that are larger in *T. fasciculata* and on a subset of genes belonging to families that are larger in *T. leiboldiana*. The top 100 significantly enriched GO terms were then evaluated. GO terms putatively associated with key innovation traits were used to list multicopy gene families of interest.

Additionally, we searched for specific genes that are known to underlie CAM evolution in these multi-copy gene families. The IDs of candidate pineapple genes for CAM were obtained from (Yardeni et al., 2021), who compiled extensive lists of genes from a diverse set of studies. For CAM, we considered all genes listed in Table S1 in this study under the categories “Differentially expressed in CAM / C3 experiment” (186 genes) (De La Harpe et al., 2020), “Positive selection in CAM / C3 shifts” (22) (De La Harpe et al., 2020), Gene families associated with CAM/C3” (79) (De La Harpe et al., 2020), “CAM-related *A. comosus*” (29) (Ming et al., 2015), “stomatal function” (48) (Christin et al., 2014), “aquaporin regulation” (24) (Vera-Estrella et al., 2012), “drought resistance” (61) (Xiao et al., 2007), “circadian metabolism” (47) (Wai et al., 2017), “malate transferase” (28) (Cosentino et al., 2013) and “circadian clock” (3) (McClung, 2006), resulting in a total of 527 genes. A separate list was made for gluconeogenesis and starch metabolism genes (288 genes) (Cushman et al., 2008). After obtaining these lists of pineapple gene IDs, we searched for their orthologs in *T. fasciculata* and *T. leiboldiana*, and investigated their presence in multi-copy gene families.

### 5.10. d_N_/d_S_ analysis

#### 5.10.1. On single-copy orthologous pairs

One-to-one orthologous genes were subjected to a test of positive selection using the non-synonymous to synonymous substitution ratio (ω = d_N_/d_S_). Gene pairs where both genes were incomplete (missing start and/or stop codon) or where the difference in total length was more than 20 % of the length of either gene were removed. We performed codon-aware alignments using the alignSequences program from MACSE v.2.05 (Ranwez et al., 2018) with options - local_realign_init 1 -local_realign_dec 1 for optimization. Pairwise d_N_/d_S_ ratios were estimated with the codeML function of PAML v.4.9. (Yang, 2007). Using a single-ratio model across sites and branches (Nssites = 0, model = 0), we tested for a fixed ω = 1 as null hypothesis, against an unfixed ω as the alternative hypothesis. Automatisation of codeML was achieved with a modified script from AlignmentProcessor (https://github.com/WilsonSayresLab/AlignmentProcessor/). The results of codeML under both the null and alternative model were compiled and significance of the result was calculated with the likelihood-ratio test (Wong et al., 2004). Multiple-testing correction was applied with the Benjamini-Hochberg method and an FDR threshold of 0.05. Orthologous gene pairs with a d_N_/d_S_ ratio larger than one and an adjusted p-value under 0.05 were considered candidate genes under divergent selection.

The d_N_/d_S_ values of all orthologous gene pairs with five or more variant sites in the MACSE alignment were used to obtain per-scaffold distributions of d_N_/d_S_ values in both genomes. We visualised d_N_/d_S_ distributions of all main scaffolds in both assemblies with boxplots and used density plots to visualise the d_N_/d_S_ distribution in rearranged chromosomes compared to all non-rearranged chromosomes. To test whether these distributions were significantly different, we ran a non-parametric test (Mann-Whitney U) between the distribution of each single rearranged chromosome and that of all non-rearranged chromosomes in each assembly.

#### 5.10.2. On duplicated orthogroups

We also performed tests of selection using d_N_/d_S_ on all orthogroups that were consisted of a single gene in *A. comosus* and a duplicated gene in either T. *leiboldiana* (1:1:2), or *T. fasciculata* (1:2:1). Only orthogroups that maintained this conformation after size correction were used in this analysis. Pairwise alignments were performed between the ortholog of one species and either paralog of the other species using MACSE. Then, ω was estimated in the same way as mentioned above.

### 5.11. RNA-seq experiment capturing photosynthetic phenotypes and expression

#### 5.11.1. Experiment set-up and sampling

To capture gene expression patterns related to CAM, we designed an RNA-seq experiment where individuals of *T. fasciculata* and *T. leiboldiana* were sampled at six time points throughout a 24-hour cycle. Six plants of each species were placed in a PERCIVAL climatic cabinet at 22 °C and a relative humidity (rH) of 68 % for 4 weeks, with a 12-hour light cycle. Light was provided by fluorescent lamps with a spectrum ranging from 400 to 700 nm. The light intensity was set at 124 µmol/m^2^s. The plants acclimated to these conditions for 4 weeks prior to sampling, during which they were watered every second day.

Leaf material from each plant was sampled every 4 hours in a 24-hour cycle starting one hour after lights went off. One leaf was pulled out of the base at each time-point without cutting. The base and tip of the leaf were then removed, and the middle of the leaf immediately placed in liquid nitrogen, then stored at −80 °C.

#### 5.11.2. Targeted metabolite analyses

To corroborate the photosynthetic phenotypes of *T. fasciculata* and *T. leiboldiana*, we measured malate abundances in the leaf throughout a 24-hour cycle. An approximate amount of 20 mg of frozen leaf material collected at six timepoints during the above-mentioned experiment was collected and ground to a powder with a TissueLyser and metal beads. Subsequent steps were performed at the Vienna Metabolomics Center (VIME, Department of Ecogenomics and Systems Biology, Vienna, Austria).

Polar metabolites were extracted in three randomised batches by modifying the procedure of (Weckwerth et al., 2004). A weighed amount of deep frozen and ground plant tissues was combined with 750 µL of ice-cold extraction solvent, consisting of methanol (LC-MS grade, Merck), chloroform (anhydrous >99 %, Sigma Aldrich), and water (MilliQ) in a ratio of 2.5:1:0.5 (v/v). Additionally, 7 µL of a solution of 10 mmol of pentaerythritol (PE) and 10 mM phenyl-β-d-glucopyranoside (PGP) respectively in water (MilliQ) were added as an internal standard mix. After ultrasonication at 4 °C for 20 minutes and centrifugation (4 min, 4 °C, 14,000 g) the supernatant was transferred to a new 1.5 mL tube (polypropylene). Another 250 µL of extraction solvent was added to the remaining pellet and after another cycle of ultrasonication and centrifugation as described before, the supernatant was combined with the previous. To induce phase separation 350 µL of water (MilliQ) was added. After thorough mixing and consecutive centrifugation (4 min, 4 °C, 14,000 g) 900 µL of the upper phase were transferred to a new 1.5 mL tube. Approximately 100 µL of the remaining polar phase of all samples were combined. The 900 µL aliquotes of this mixed sample were used as quality control during measurements. The polar phases and the aliquotes of the sample mix were dried in a vacuum centrifuge for 5 hours at 30 °C and 0.1 mbar.

The dried extracts were derivatised as described earlier (Doerfler et al., 2013) by dissolving the metabolite pellet carefully in 20 µL of 40 mg of methoxyamine hydrochloride (Sigma Aldrich) in 1 mL pyridine (anhydrous >99,8 %, Sigma Aldrich). After incubation at 30 °C and 700 rpm for 1.5 hours on a thermoshaker, 80 µL of *N*-methyl-*N*-trimethylsilyl-trifluoroacetamid (Macherey-Nagel) were added. The samples were incubated for 30 min at 37 °C and 750 rpm and consecutively centrifuged for 4 min at room temperature and 14,000 g.

Metabolite analysis was performed on an Agilent 7890B gas chromatograph equipped with a LECO Pegasus® BT-TOF mass spectrometer (LECO Corporation). Derivatised metabolites were injected through a Split/Splitless inlet equipped with an ultra-inert single tapered glass liner with deactivated glass wool (5910-2293, Agilent Technologies), a split ratio of 1:25 was used and the temperature was set to 230 °C. Components were separated with helium as carrier gas on a Restek Rxi-5Sil MS column (length: 30 m, diameter: 0.25 mm, thickness of film: 0.25 µm). The initial oven temperature was set to 70 °C held for 1 minute and ramped with a rate of 9 °C per minute until reaching 340 °C held for 10 minutes. Collection of spectra started after an acquisition delay of 280 seconds with a detector voltage of 1692.5 V, a rate of 15 spectra per second and a mass range of 50 ̶ 500 *m/z*. Retention indices were calculated based on the retention times of the alkane mixture C_10_-C_40_ run within each of the 2 batches. Samples were measured in randomised order and randomly distributed across the batches. Within each batch, a mixture of standard metabolites was measured for MSI level I identification of metabolites. Deconvolution, annotation, and processing of chromatograms was performed according to (Zhang et al., 2023) using ChromaTOF® (Version 5.55.29.0.1187, LECO Cooperation) and MS-DIAL, version 4.7 (Tsugawa et al., 2015). Areas of derivatisation products of single metabolite were summed and normalised by the main targeted ion content of each sample.

#### 5.11.3. RNA extraction and sequencing

Using the same sampled leaf material as for targeted metabolite analyses, total RNA was extracted for each sample and timepoint in randomised batches of 4-6 samples, using the QIAGEN RNeasy® Mini Kit in an RNAse free laboratory. Samples were digested using the kit’s RLT buffer with 1 % Beta-mercaptoethanol. Elution was done in two steps. The purity and concentration of the extractions was measured using Nanodrop, and RIN and fragmentation profiles were obtained with a Fragment Analyzer™ system. RNA libraries were prepared by the Vienna Biocenter Core Facilities (VBCF) using a NEBNext stranded mRNA kit before sequencing 150-bp paired-end reads on one lane of Illumina NovaSeq S4.

#### 5.11.4. RNA-seq data processing

The raw RNA-seq data was evaluated with FastQC (https://www.bioinformatics.babraham.ac.uk/projects/fastqc/) and MultiQC (Ewels et al., 2016), then quality trimmed using AdapterRemoval v.2.3.1 (Schubert et al., 2016) with settings –trimns --trimqualities --minquality 20 --trimwindows 12 --minlength 36. The trimmed data was then aligned to both the *T. fasciculata* and *T. leiboldiana* genomes using STAR v.2.7.9 (Dobin et al., 2013) using GFF files to specify exonic regions. Because mapping bias was lowest when mapping to *T. fasciculata* (see SI Note 8, Fig. S13, S14), our main analyses have been performed on the reads mapped to this genome. However, the alignments to *T. leiboldiana* were used for verification or expansion of the main analysis (SI Note 9).

#### 5.11.5. Differential Gene Expression analysis

We quantified read counts per exon using FeatureCounts from the Subread package v.2.0.3. (Liao et al., 2014) for paired-end and reversely stranded reads (-p -s 2). The counts were then summed up across exons per gene to obtain gene-level counts. The composition of the count data was investigated with PCA in EdgeR (Robinson et al., 2009). Then, counts were normalised using the TMM method in EdgeR, and every gene with a mean cpm < 1 was removed. We ran a differential gene expression (DE) analysis in maSigPro (Conesa et al., 2006), which detects genes with differential diurnal expression profiles between species using a regression approach. *T. leiboldiana* was used as the baseline in this analysis. Significant DE genes were then clustered using the hclust algorithm into modules, with the number of modules being determined with the K-means algorithm. Expression curves were plotted by taking the average expression in TPM (Transcripts per Million) across all replicates per species at each time. We calculated TPM by dividing the raw read count by the exonic length of the gene (RPK), which we then divided by the total sum of RPK values. Expression curves for entire clusters (Fig. S8) were plotted by median-centring the log(TPM) of each gene and time point against the median of all genes at each time point, while expression curves for individual genes or gene families (Fig. 5c, S5, S7) report average TPM.

GO term enrichments were performed for each cluster using the R package TopGO (Alexa and Rahnenführer, 2009). Separately, known candidate genes underlying CAM and starch metabolism (See Methods, Section 5.9.) were searched among differentially expressed genes.

#### 5.11.6. Annotation and enrichment of circadian clock-related motifs in promoter sequences

We counted the occurrences of four known circadian clock-related motifs in the 2-kb upstream regions of DE genes: the Morning Element (MOE: CCACAC) (Michael et al., 2008), the Evening Element (EE: AAAATATC) (Hudson and Quail, 2003), the CCA1-binding site (CBS: AAAAATCT) (Franco-Zorrilla et al., 2014) and the G-box element (G-box: CACGTG) (Michael and McClung, 2002). The same was done for all other curated genes that were not DE, which we considered as background sequences. We calculated the per-kb frequency of each motif based on the counts and total promoter length (2000 x number of genes) for both sets of genes. The percentage of change in frequency was calculated between both sets for each motif. Significance of frequency changes of circadian motifs in promotor regions of DE genes compared to non-DE genes were calculated with the Wilcoxon Rank Sum Test.

For a small set of genes known to underly key CAM enzymes, we counted the occurrence of each motif in both the 2-kb upstream region of every homolog of that gene (including non-DE paralogs) in both species, to annotate and describe circadian motifs in promoter sequences in detail. The detection of motifs was extended to 3-kb regions to allow for a more distant presence of motifs.

### 5.12. Intersecting findings of gene family evolution, TE insertion ad differential gene expression

#### 5.12.1. Spatial distribution of DE genes

The previously calculated per 1 kb-window counts of robust genes was used to obtain the per-window proportion of DE genes. This was then visualised with circlize as described above. Correlations of total DE gene count per scaffold and scaffold size was calculated with Kendall’s rank correlation test after testing for normality with the Shapiro-Wilk test.

#### 5.12.2. Gene family evolution and differential gene expression

Orthogroups were split based on relative family size in *T. fasciculata* (F) versus *T. leiboldiana* (L) in the following categories: Single-copy orthogroups (F = 1 : L = 1), orthogroups with family size larger in *T. fasciculata* (F > L), orthogroups with family size smaller in *T. fasciculata* (F < L), orthogroups with equal family sizes that are larger than 1 (F = L). Orthogroups unique to one species (F:0 or 0:L) were not considered in this analysis. We counted the number of orthogroups belonging to each category for the full orthogroup set, for the subset of orthogroups containing DE genes (DE orthogroups) and for the subset of orthogroups containing DE genes that have been previously described as CAM-related (CAM-DE orthogroups). We then tested whether counts in each orthogroup category were enriched in DE orthogroups and CAM-DE orthogroups compared to all orthogroups. For comparisons of all orthogroups versus DE orthogroups, we used the chi-square test of independence in R. For the comparison of CAM-DE orthogroups versus all orthogroups, we used Fisher’s exact test due to small sample sizes with 2×2 contingency tables of each orthogroup category versus all other categories, and DE genes versus non-DE genes. To study the effect of the reference genome used on our findings on gene family evolution in DE genes, we performed the same analysis on read counts obtained from mapping to *T. leiboldiana* (SI Note 9).

#### 5.12.3. Transposable element insertions and differential gene expression

Intronic TE insertions were obtained using *bedtools intersect* on the GFF files of the TE and gene annotations of both species. We used the full transcript length of a gene (feature = “mRNA” in GFF file) for this analysis, and only applied “known” TE annotations and the set of curated genes. This resulted in a dataset reporting the number of TE insertions per gene. We also obtained TE counts for genic regions including the 3-kb upstream region, by using *bedtools slop* with options *-l 3000 -r 0 -s*. For analyses on specific TE classes, we calculated TE insertion counts for the following four TE categories: LTR-Copia, LTR-Gypsy, Helitron and DNA transposon.

We then performed two tests on the resulting TE counts per gene: (1) whether the proportion of genes with one or more TE insertions is elevated in DE genes compared to the full gene set (chi-square test), and (2) whether the rate of TE insertions per gene measured, as the total count of intersections for each gene annotation with a TE annotation, is elevated in DE genes compared to non-DE genes (Mann-Whitney U test).

The same test was also applied to a restricted set of DE genes previously described as CAM-related, or involved in starch metabolism and gluconeogenesis. Then, genes of interest with a TE insertion rate higher than twice the genome-wide average were selected and the difference in number of TE insertions between orthologs of *T. leiboldiana* and *T. fasciculata* was taken in case of a one-to-one relationship.

## 6. Accession Numbers

The genome assemblies and raw data used in this study are available at NCBI-SRA under BioProject <u>PRJNA927306</u>. Specifically, the *T. fasciculata* genome assembly TFas_v1 can be downloaded here: https://ncbi.nlm.nih.gov/datasets/genome/GCA_029168755.1/. The *T. leiboldiana* genome assembly TLei_v1 can be found at: https://www.ncbi.nlm.nih.gov/datasets/genome/GCA_029204045.1/. The annotation of both genomes is available on the Github repository at: https://github.com/cgrootcrego/Tillandsia_Genomes, together with the list of orthogroups, counts table used for RNA-seq analyses, full GO term enrichment results, and all scripts written for this manuscript. The *A. comosus* sequences used in this study stem from BioProject PRJNA371634 (F153): https://www.ncbi.nlm.nih.gov/bioproject/PRJNA371634/ and BioProject PRJNA747096 (CB5): https://www.ncbi.nlm.nih.gov/bioproject/PRJNA747096. Information on all accessions used in this study can be found in Supplemental Table 1.

### 7. Supplemental data

Supplemental Figure S1. Per-metabolite loadings of PC1 and PC2

Supplemental Figure S2. Distribution of scaffold sizes and orthologous gene counts for the top 100 largest scaffolds of both assemblies.

Supplemental Figure S3. TE, GC, and gene content at three examples of syntenic chromosome triplets.

Supplemental Figure S4. In-depth visualisation of large-scale rearrangements between *T. fasciculata* and *T. leiboldiana*.

Supplemental Figure S5. Genome-wide distribution of d_N_/d_S_ values between single-copy orthologous genes.

Supplemental Figure S6. Heatmap of z-score normalised expression values (log(TPM)) of CAM-related DE genes featured in Figure 6.

Supplemental Figure S7. Average expression curve of PEPC kinase (PPCK) in *T. fasciculata* and *T. leiboldiana*.

Supplemental Figure S8. Per-gene expression curves of all differentially expressed genes clustered in co-expression modules.

Supplemental Figure S9: Relationship between DE gene count per scaffold and scaffold size. Supplemental Figure S10. Distribution of DE genes across the genome.

Supplemental Figure S11. Average expression curve of Aquaporin 2-6 in *T. fasciculata* and *T. leiboldiana*.

Supplemental Figure S12. Distribution of mean per-gene coverage across different gene family categories.

Supplemental Figure S13. Unique mapping rates of RNA-seq reads to three different reference genomes.

Supplemental Figure S14. Multi-mapping rates of RNA-seq reads to three different reference genomes.

Supplemental Figure S15. Genome size measurement histograms.

Supplemental Figure S16. Mitotic metaphase chromosomes and karyotypes of *Tillandsia fasciculata* and *Tillandsia leiboldiana*.

Supplemental Figure S17. Heterozygosity and genome size estimation with a k-mer based approach.

Supplemental Figure S18. Distribution of heterozygous sites per 1000 mappable variants. Supplemental Table S1. List of accessions used in this study.

Supplemental Table S2. Assembly statistics.

Supplemental Table S3. TE Abundances.

Supplemental Table S4. Orthology statistics.

Supplemental Table S5. List of CAM-related expanded orthogroups.

Supplemental Table S6. Full list of candidate genes under positive selection.

Supplemental Table S7. Full list of co-expression modules.

Supplemental Table S8. Per-kb frequencies of four promoter motifs associated with circadian clock transcription factors in DE and non-DE genes.

Supplemental Table S9. Annotation of circadian promoter motifs in the upstream regions of CAM genes.

Supplemental Table S10. List of DE genes related to CAM, starch metabolism and gluconeogenesis with a high TE insertion rate.

Supplemental File 1. Supplementary Notes.

Supplemental Dataset 1. Metabolomic Compound Abundances reported as raw area read-outs from MS-Dial, including the level of identification according to the Metabolomics Standards Initiative.

## Acknowledgments

*In memory of Christian Lexer – we will treasure your enthusiasm, guidance, and memory always*. This work was financially supported by the Austrian Science Fund (FWF) through the doctoral programme (DK) grant W1225-B20 to a faculty team including C.L. and O.P., FWF grant P35275 to O.P., and by the professorship start-up grant of Christian Lexer at the University of Vienna BE772002. We thank Joachim Hermisson, Magnus Nordborg, Andrew Clark, Nicholas Barton, Virginie Courtier-Orgogozo, John Parsch, Andreas Futschik, Rui Borges, Aglaia Szukala, Florian Schwarz, Marta Pelizzolla and Ahmad Muhammad for insightful discussions, advice, and feedback. We thank Gert Bachmann and Eline de Vos for help with the setup of the RNA-seq experiment; Peter Bak, Andreas Franzke, Nils Köster and Helmut & Lieselotte Hromadnik for their generous donations of *Tillandsia* accessions; and Thelma Barbarà for assistance during RNA extractions. We also thank Barbara Knickmann, Viktor Vagovics and Manfred Speckmaier for the care of *Tillandsia* plants at the Botanical Garden of the University of Vienna. Lastly, we thank Alisa Tscherko for assistance with trichome photography. Computational resources were provided by the Life Science Computer Cluster (LiSC) of the University of Vienna and the Vienna Scientific Cluster.

## 8. Author Contributions

This study was conceived by CL, JH, OP, TL and CGC. Sampling was conducted by MHJB, WT, GY and MDH. Laboratory work was conducted by MHJB, SS, LACS and CGC. Cytogenetic work was performed by HWS and EMT. The RNA-Seq experiment and DE analysis was conducted by CGC under the guidance of KH and OP. Analyses were performed by CGC, JH, GY, TL, and FB. The manuscript was primarily written by CGC and amended following the dedicated reading and feedback of all co-authors, especially KH, TL, and OP.

## Notes

https://github.com/cgrootcrego/Tillandsia_Genomes

### Competing Interest Statement

The authors have declared no competing interest.

### Summary of Updates

New results added, figure 1 updated.

