## Supplementary Information for "CAM evolution is associated with gene family expansion in an explosive bromeliad radiation"

**1. SUPPLEMENTARY FIGURES 2**
**2. SUPPLEMENTARY TABLES 18**
**3. SUPPLEMENTARY NOTES 28**
**NOTE 1: GENOME SIZE AND KARYOTYPE OF T. FASCICULATA AND T. LEIBOLDIANA 28**
**NOTE 2: PRE-ASSEMBLY ESTIMATION OF PER-ACCESSION HETEROZYGOSITY 28**
**NOTE 3: IDENTIFYING MAIN SCAFFOLDS IN DE NOVO ASSEMBLY 29**
**NOTE 4: ON THE SPATIAL DISTRIBUTION OF GC AND TE CONTENT IN BROMELIAD GENOMES 30**
**NOTE 5: IDENTIFYING LARGE-SCALE REARRANGEMENTS BETWEEN T. FASCICULATA AND T. LEIBOLDIANA 31**
**NOTE 6: SELECTING RAPIDLY EVOLVING GENE FAMILIES 32**
**NOTE 7: DETAILED DESCRIPTION OF CANDIDATE GENES FOR POSITIVE SELECTION 33**
**NOTE 8: CHOICE OF REFERENCE GENOME, MAPPING BIAS AND MAPPING STATISTICS FOR RNA-SEQ ANALYSIS 35**
**NOTE 9: DIFFERENTIAL GENE EXPRESSION USING THE T. LEIBOLDIANA ASSEMBLY 36**
**NOTE 10: ON THE DIFFERENTIAL EXPRESSION AND GENE FAMILY EXPANSION OF CAM-RELATED GENES 37**
**NOTE 11: ON THE SUCCESS OF DE NOVO ASSEMBLY OF HIGHLY REPETITIVE GENOMES 40**
**4. REFERENCES 41**

### 1. Supplementary Figures

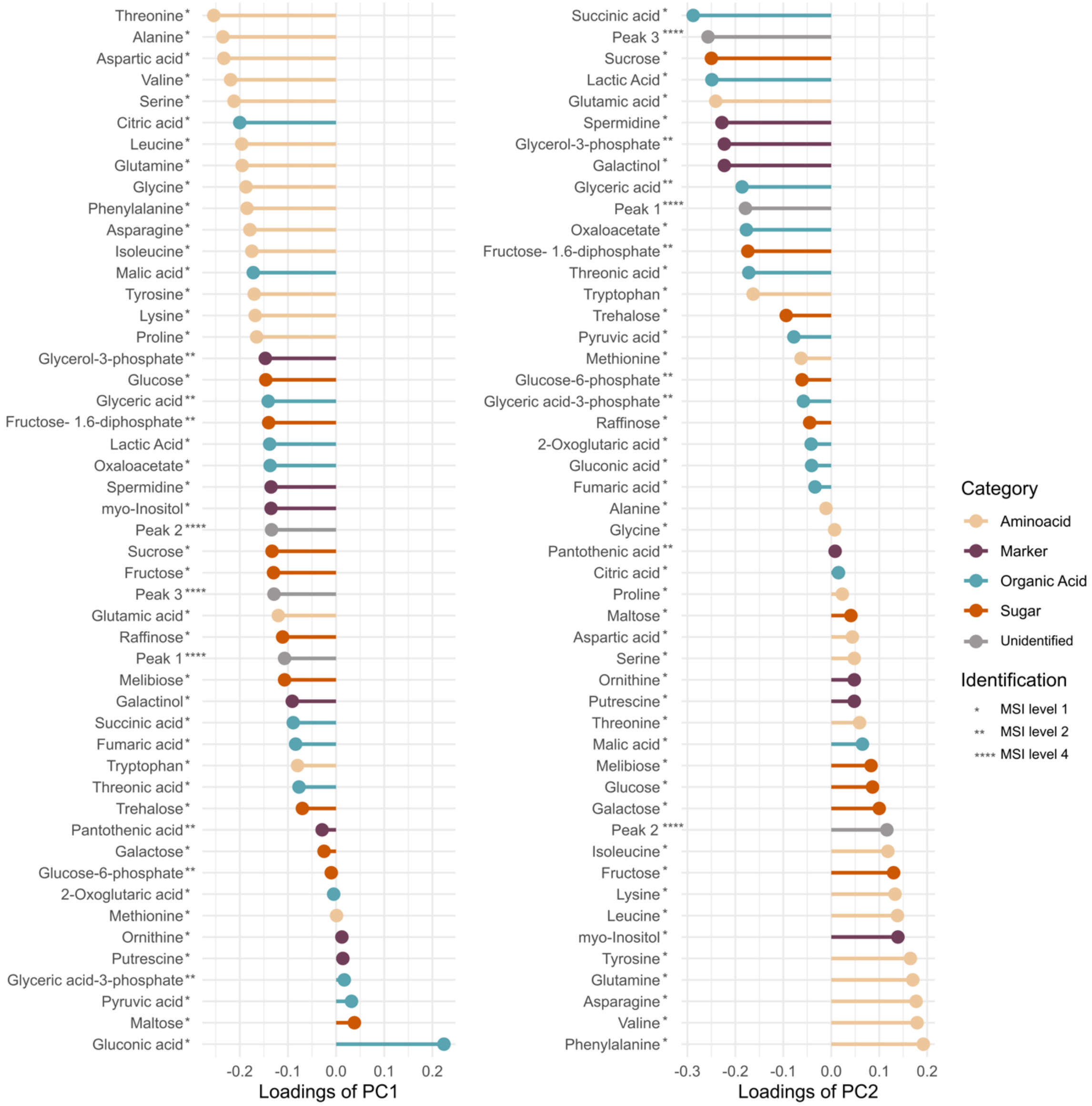


*Figure S1: Loadings of 45 individual metabolites on PC1 (left) and PC2 (right). Besides amino acids, the organic acids malate, citrate and gluconic acid contribute most to differentiation along PC1. Sugars appear as important contributors to differentiation along PC2. MSI Level 1: identified compound (mass spectra and retention index compared to reference compounds analyzed under same conditions). MSI level 2: Putatively annotated compound (EI mass spectra and retention index compared to in-house library). MSI level 4: Unidentified compound. Supports Figure 2.*

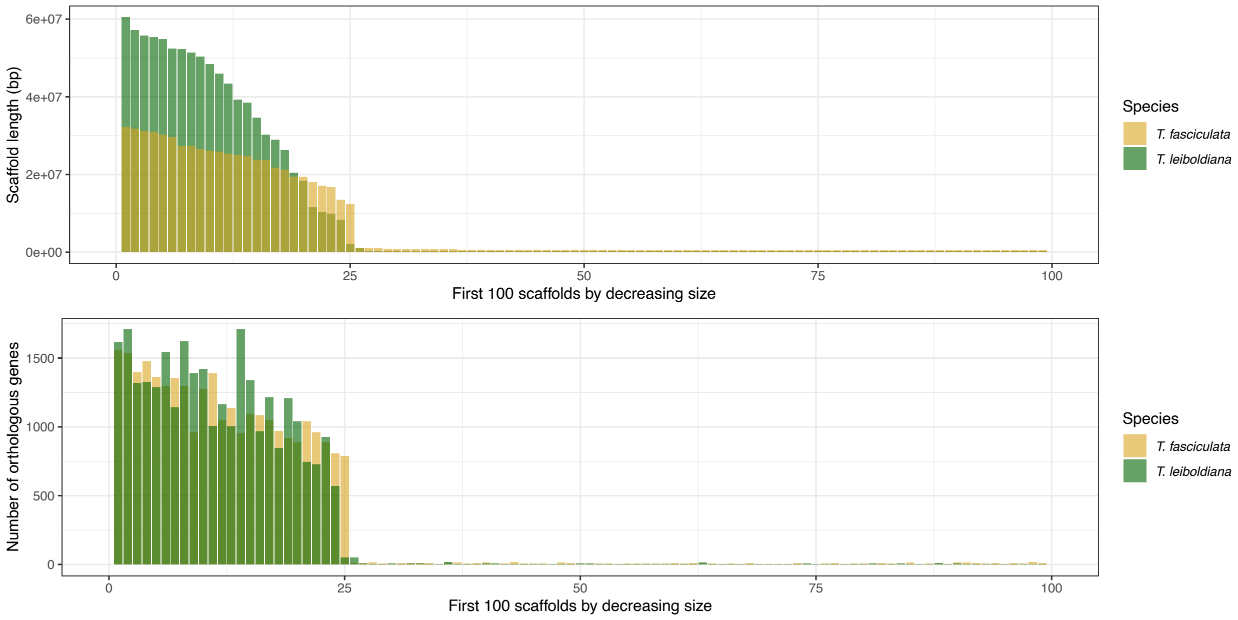


*Figure S2: Distribution of scaffold sizes (top) and count of orthologous genes per scaffold (bottom) for the top 100 largest scaffolds of both assemblies. Supports Figure 3.*

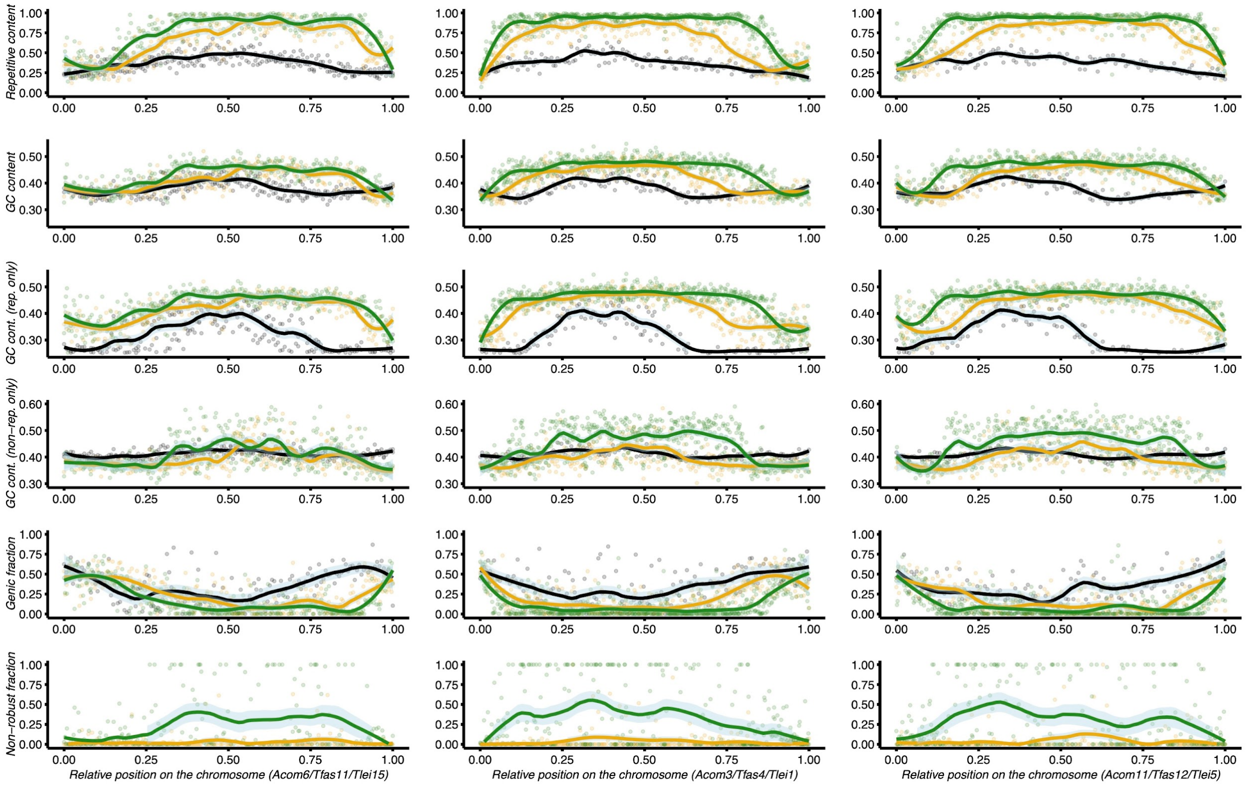


*Figure S3: TE, GC, and gene content at three examples of syntenic chromosome triplets of* A. comosus *(black),* T. fasciculata *(yellow) and* T. leiboldiana *(green). Each column represents a separate chromosome triplet. Each dot corresponds to an estimate in a non-overlapping 100 kb window. The line corresponds to the local regression (loess). Row-wise, from top to bottom, the plots show: (1) per-window proportion of soft-masked position in the assemblies (repetitive content), (2) GC content at all non-N positions (soft-masked or not), (3) GC content at soft-masked positions only, (4) GC content at positions that were not soft-masked,, (5) per-window proportion of bases falling in genes (genic fraction) and (6) the proportion of the genic fraction corresponding to non-robust genes (i.e. 1 minus this fraction corresponds to "robust" gene regions). This latter information is only provided for our two reference assemblies. Supports Figure 3.*

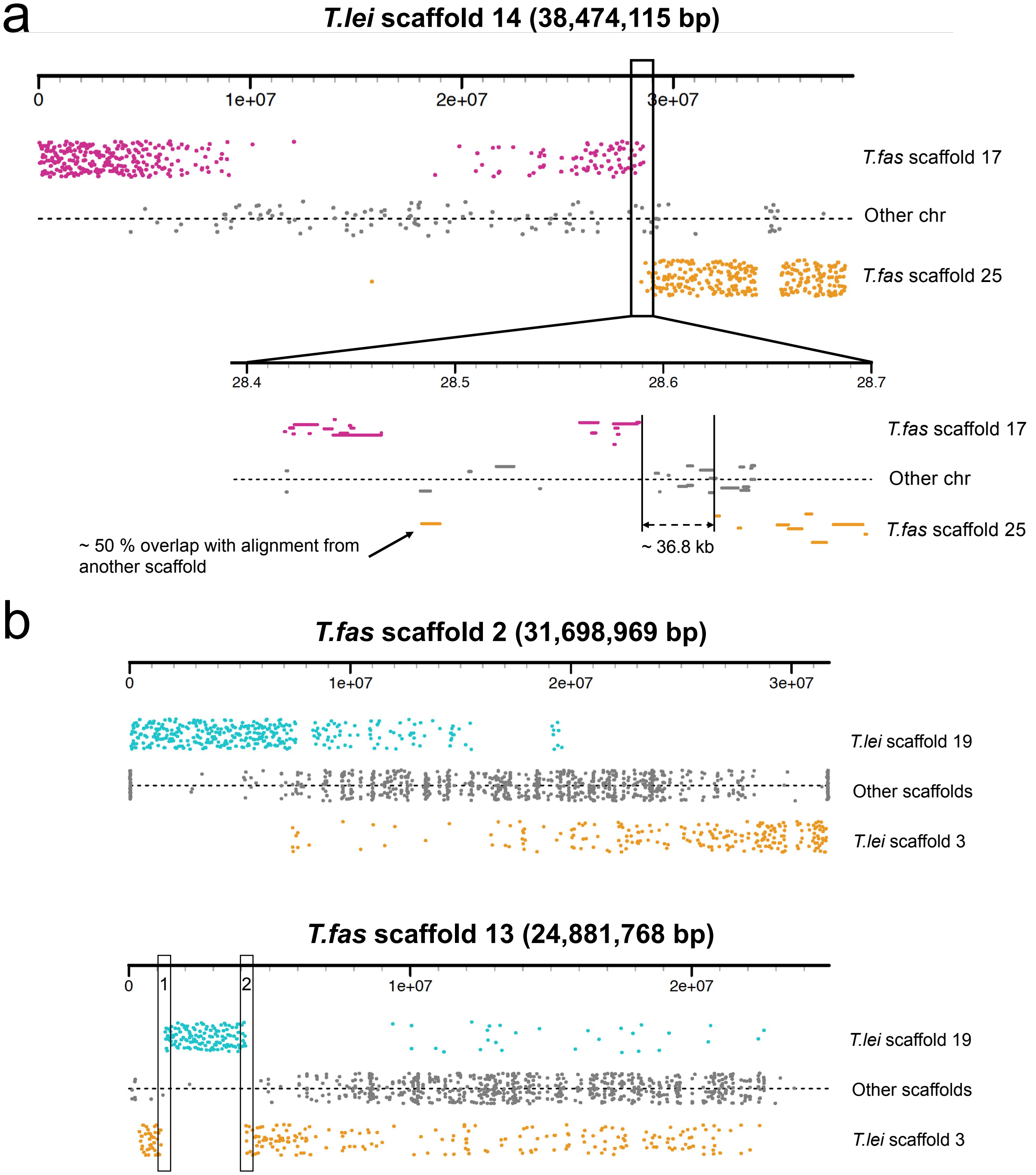
a
T.lei scaffold 14 (38,474,115 bp)
0
1e+07
2e+07
3e+07
T.fas scaffold 17
Other chr
T.fas scaffold 25
28.4
28.5
28.6
28.7
T.fas scaffold 17
Other chr
T.fas scaffold 25
~ 50 % overlap with alignment from another scaffold
~ 36.8 kb
b
T.fas scaffold 2 (31,698,969 bp)
0
1e+07
2e+07
3e+07
T.lei scaffold 19
Other scaffolds
T.lei scaffold 3
T.fas scaffold 13 (24,881,768 bp)
0
1
2
1e+07
2e+07
T.lei scaffold 19
Other scaffolds
T.lei scaffold 3

C

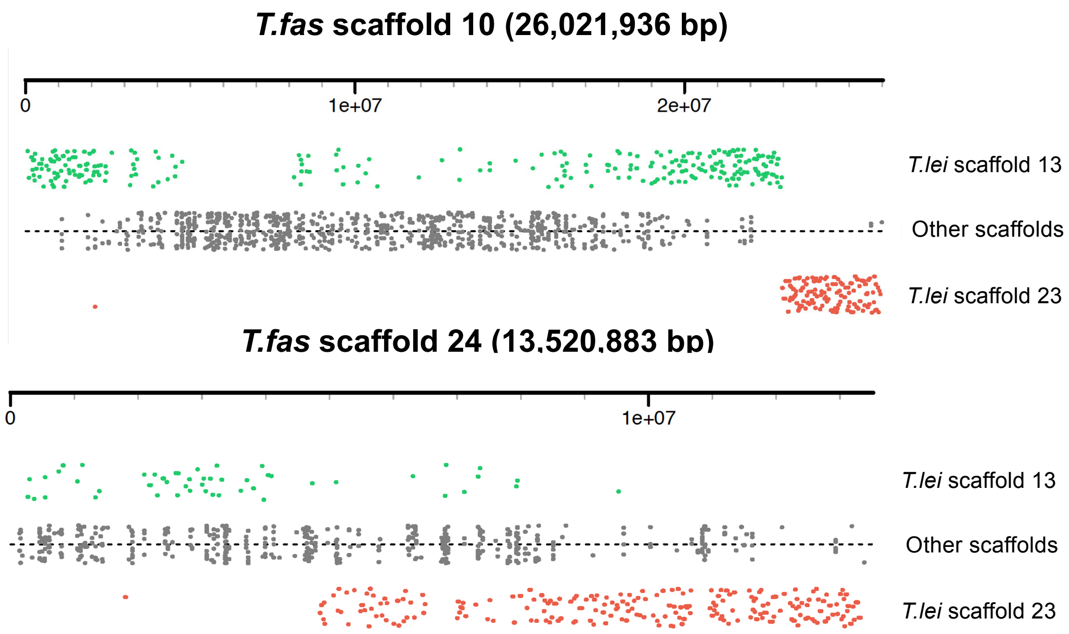


*Figure S4: In-depth visualisation of large-scale rearrangements between* T. fasciculata *and* T. leiboldiana *based on local alignments with less than a 90 % overlap with any other alignment. a) Potential fusion of scaffold 14 in* T. leiboldiana*, with enlargement of the breakpoint area. B) Translocation 1 – alignments were too sparse to determine a breakpoint on scaffold 2. Breakpoint 1 on scaffold 13 in* T. fasciculata *was not supported by raw PacBio alignments, however breakpoint 2 was. C) Translocation 2 – we find PacBio alignment support for the breakpoint on scaffold 10, but alignments were too sparse on scaffold 24 to determine a breakpoint. For more in-depth analysis and visualization of PacBio alignments, see the document 'Tfas_Tlei_rearrangements.pdf' on our github repository. Supports Figure 3.*

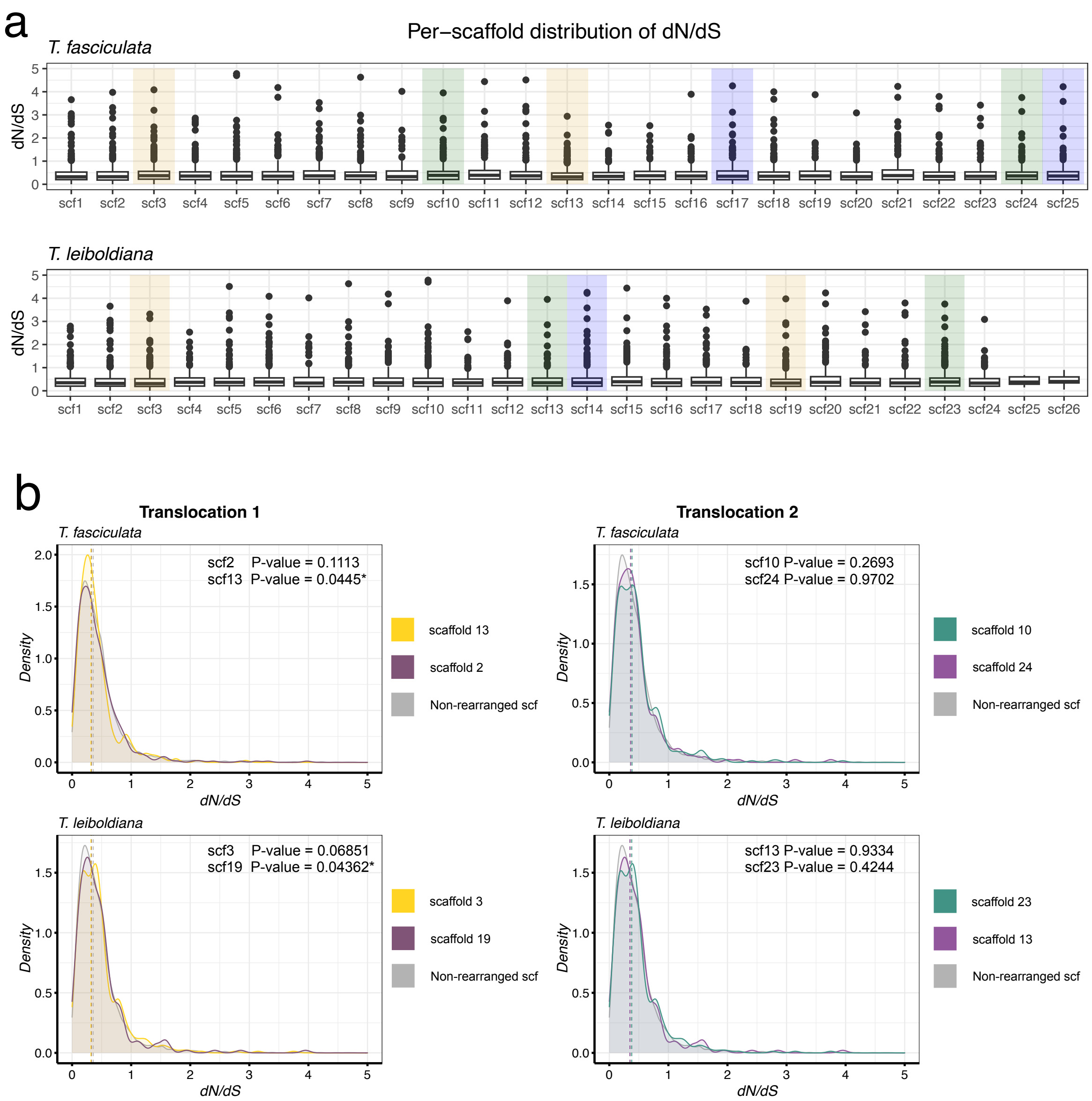


*Figure S5: Genome-wide distribution of $d_N/d_S$ values between single-copy orthologous genes. A) Boxplot of $d_N/d_S$ values in each scaffold of both assemblies. For ease of reading, the y-axis is cut-off at a $d_N/d_S$ value of five. Therefore, candidate genes with high values are not shown here. Scaffolds highlighted in colours are involved in the three reported large-scale rearrangements: (1) chromosomal fusion in* T. leiboldiana *(blue), (2) translocation 1 (yellow), and (3) translocation 2 (green) B) Distribution of $d_N/d_S$ values of all non-rearranged chromosomes versus chromosomes involved in translocations. P-values were obtained through the Mann Whitney U test, comparing dN/dS values of the respective scaffold against the background of non-rearranged scaffolds. Supports Figure 4.*

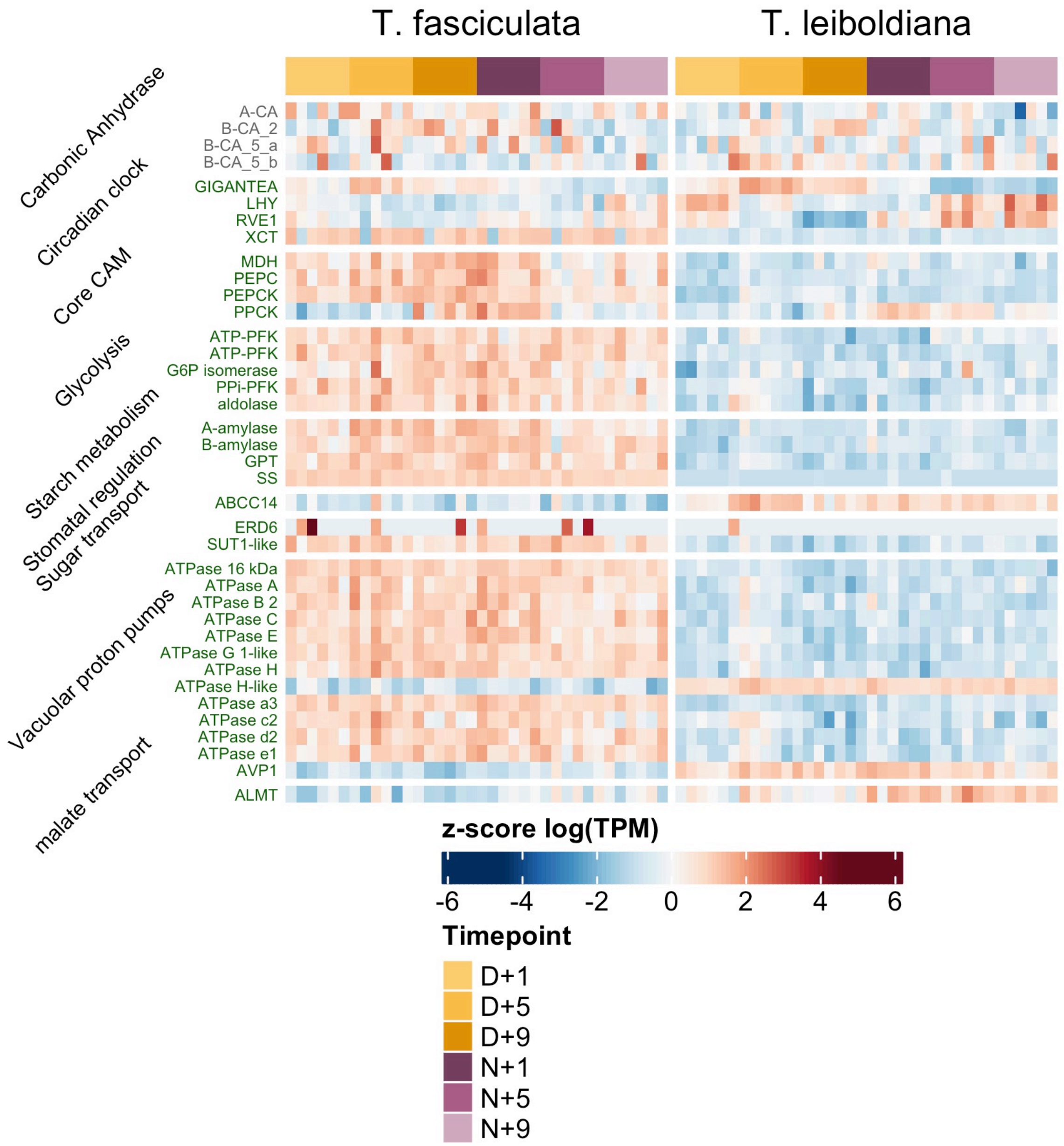


*Figure S6: Centred expression values (z-score of log(TPM)) for CAM-related DE genes featured in Figure 6. Genes in grey were not considered differentially expressed. SI Note X contains a detailed explanation and interpretation of the expression curves of these genes. Supports Figure 6.*

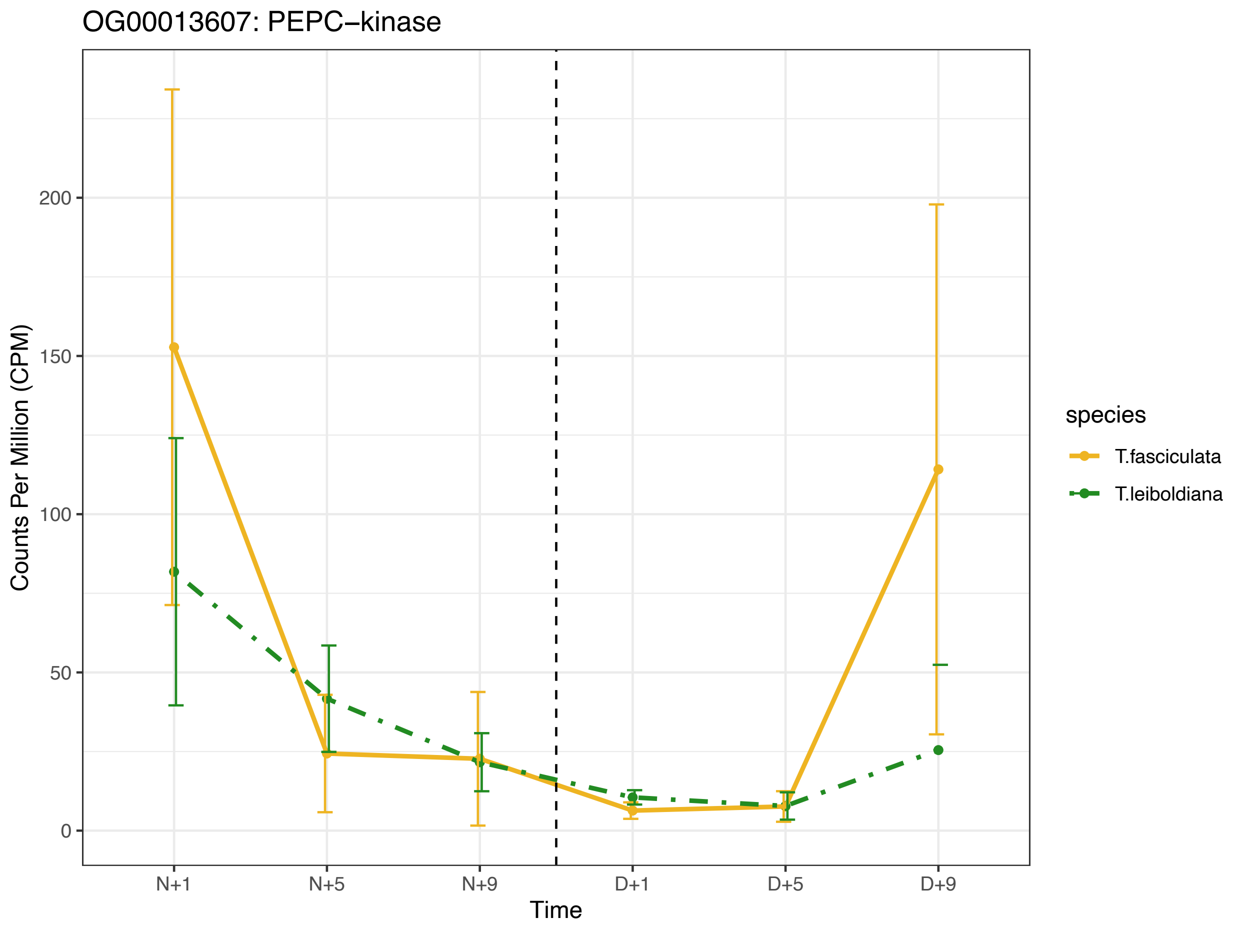


*Figure S7: Average expression curve of PEPC kinase (PPCK) in* T. fasciculata *and* T. leiboldiana *with standard deviation. The dashed vertical line marks the point where the light was switched on. Time is indicated in hours after the lights go off (N=Night) and after they go on (D=Day).Supports Figures 5 and 6.*

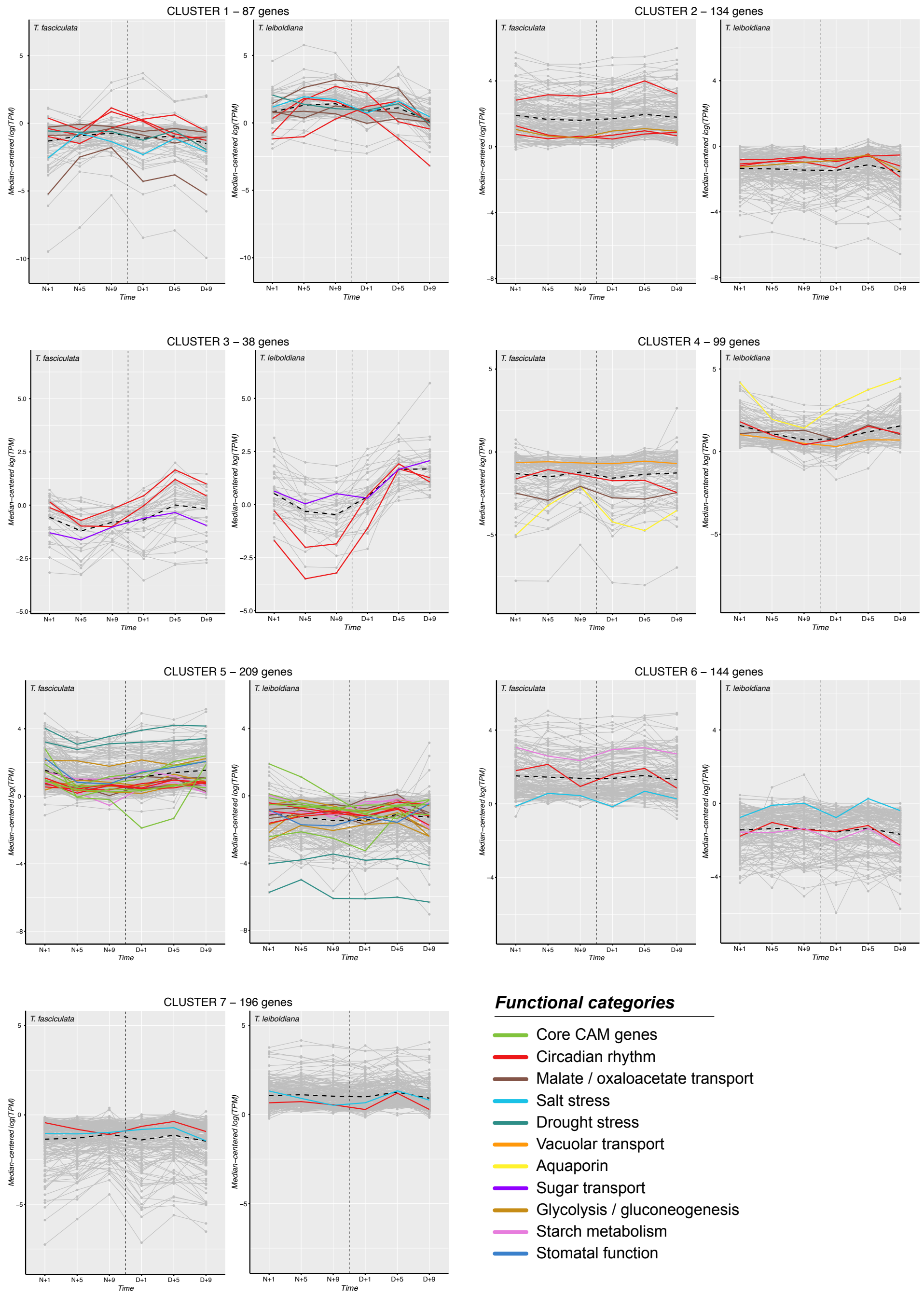


*Figure S8: Per-gene expression curves of all differentially expressed genes, spread over 7 co-expression clusters inferred with MaSigPro (and* T. fasciculata *as reference genome). The dashed vertical line marks the point where the light was switched on. Time is indicated in hours after the lights go off (N=Night) and after they go on (D=Day). Highlighted expression curves represent candidate genes underlying CAM-related functions. The colours correspond to specific subfunctions, laid out in the legend below. Supports Figures 5 and 6.*

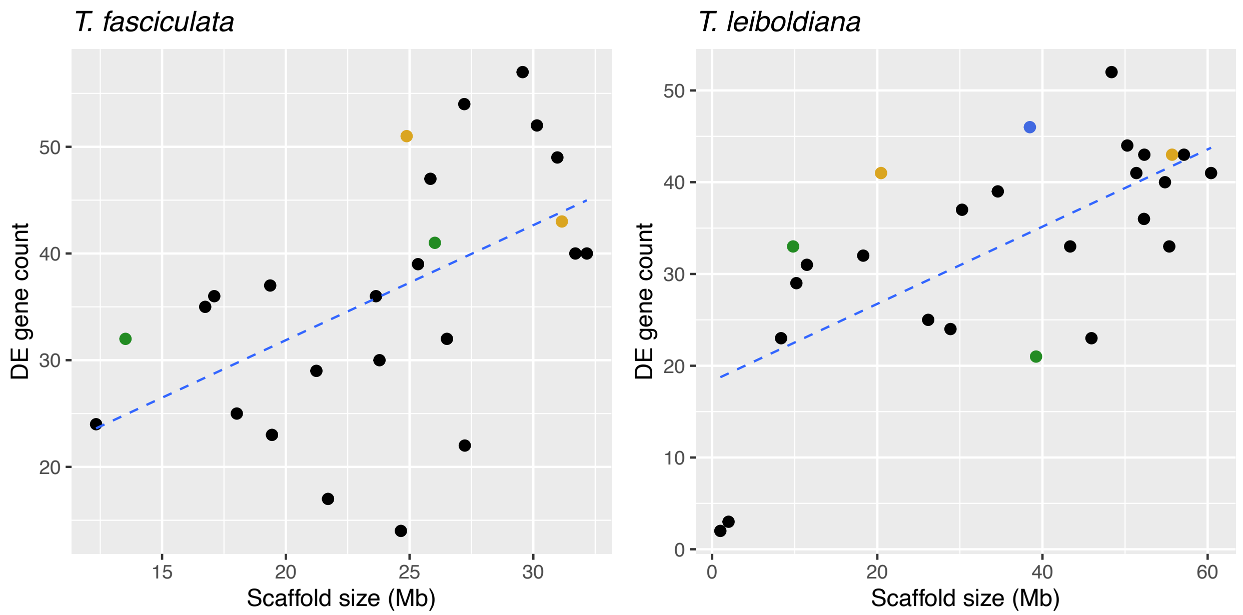


*Figure S9: Relationship between DE gene count per scaffold and scaffold size. Scaffolds involved in large-scale rearrangements are highlighted in colours; translocation 1 (yellow), translocation 2 (green), chromosomal fusion (blue). Dashed line computed with geom_smooth using a linear model. Supports Results section 3.9.1.*

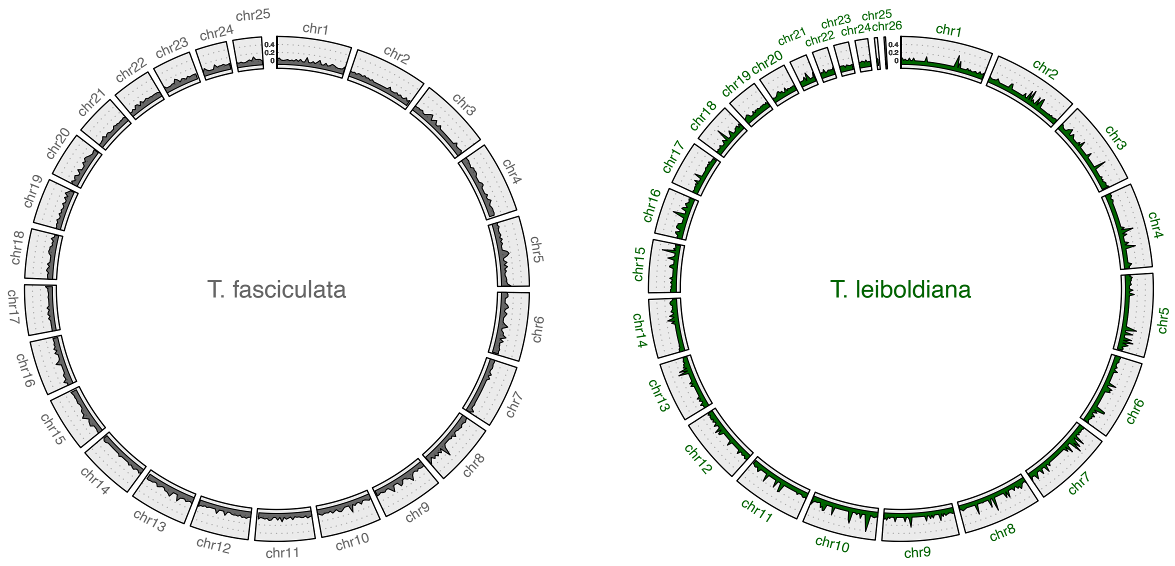


*Figure S10: Proportion of genes per 1 Mb window that are differentially expressed across the* T. fasciculata *(left) and* T. leiboldiana *(right) assembly. Supports Results section 3.9.1.*

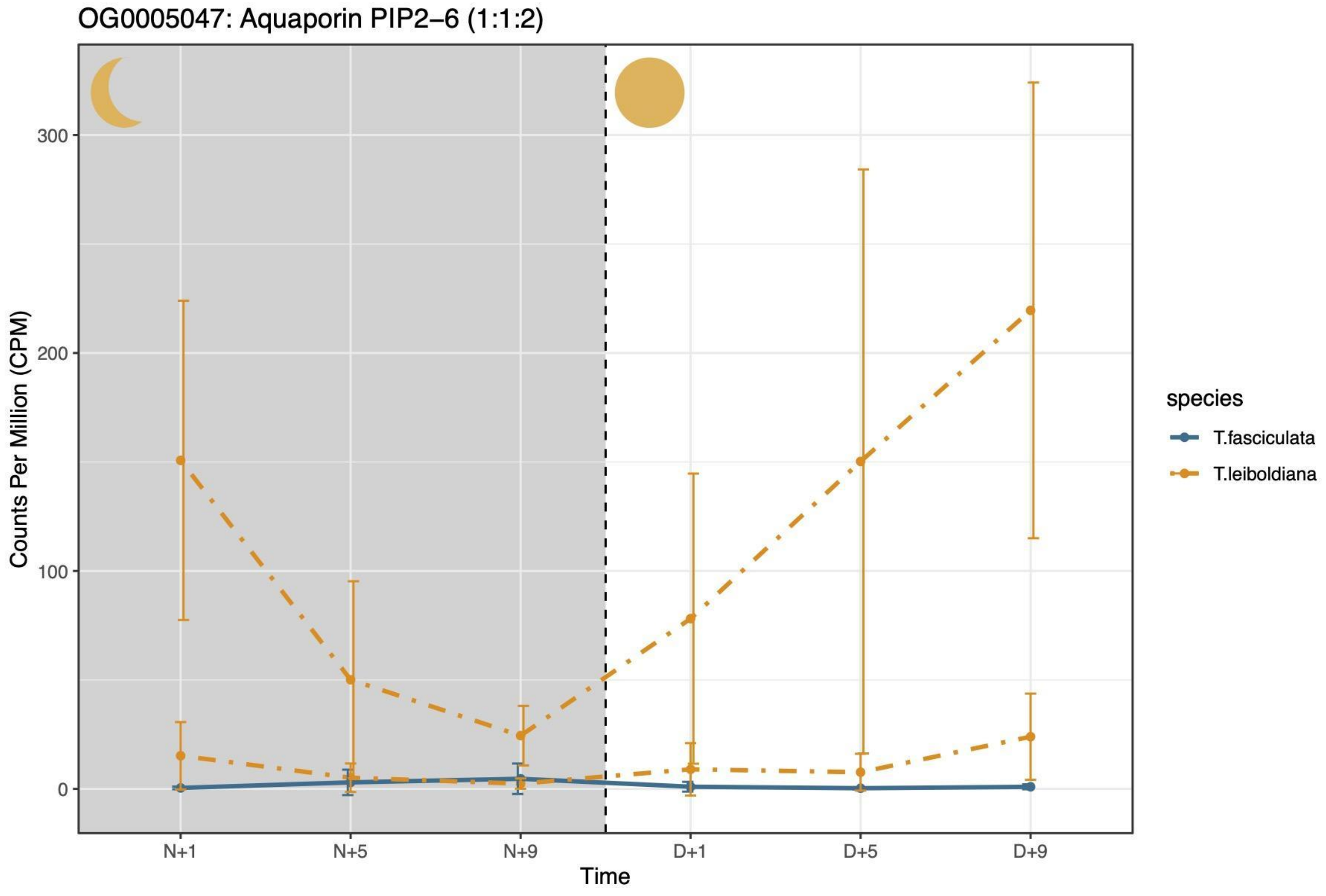


*Figure S11: Average expression curve of Aquaporin PIP2-6 in* T. fasciculata *and* T. leiboldiana *with standard deviation. The dashed vertical line marks the point where the light was switched on. Time is indicated in hours after the lights go off (N=Night) and after they go on (D=Day). Supports Figures 5 and 6.*

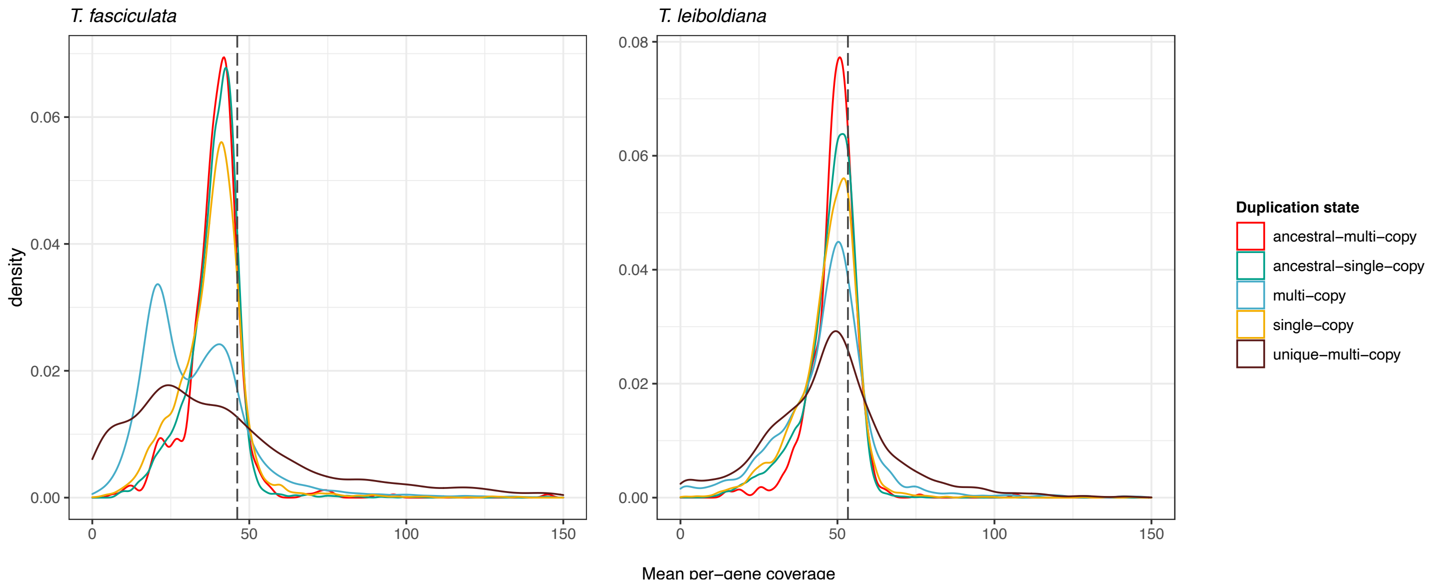


*Figure S12: Mean per-gene coverage distribution across different gene family categories in* T. fasciculata *(left) and* T. leiboldiana *(right). For further explanation of the different categories, see SI Note 7. Grey dashed lines indicate the whole-genome mean coverage. Supports Figure 5.*

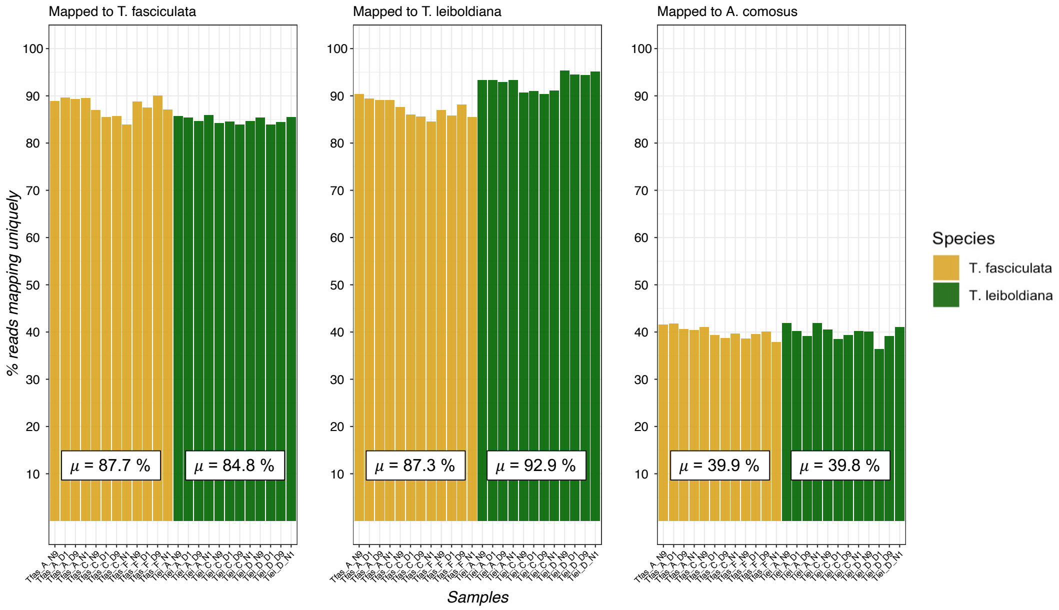


*Figure S13: Percentage of uniquely mapping RNA-seq reads for 24 samples to three different genome assemblies (*T. fasciculata*,* T. leiboldiana *and* A. comosus*). The mean unique mapping rate per species and genome is reported inside the white squares. Supports Figure 5.*

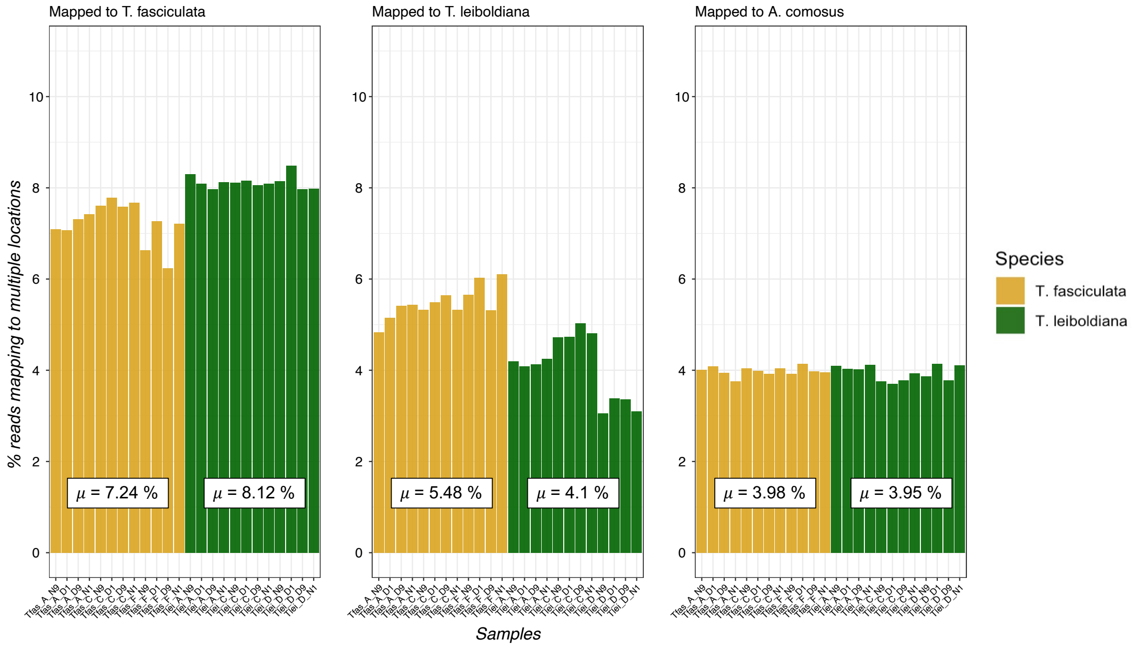


*Figure S14: Proportion of RNA-seq reads mapping in multiple genomic locations for 24 samples mapped to three different genome assemblies, with the mean mapping rate for each species and genome highlighted in white squares. Supports Figure 5.*

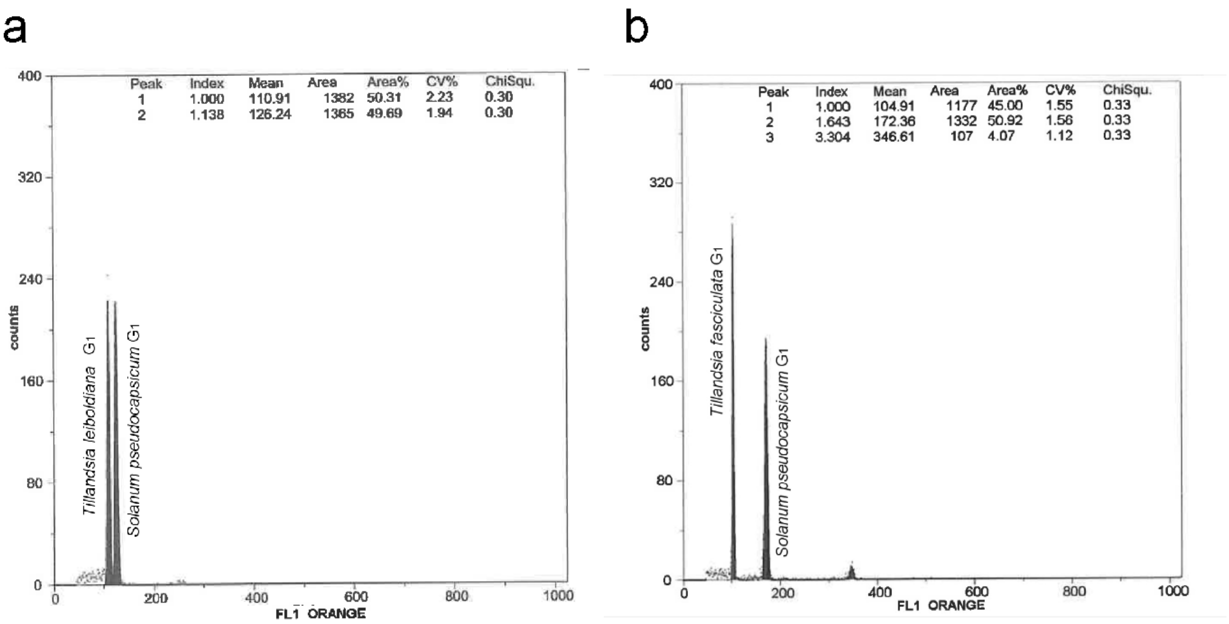


*Figure S15: Genome size measurement histograms of one exemplary run of* Tillandsia fasciculata *and* T. leiboldiana *showing the mean $G_1$ nuclei peak positions on the x-axis (fluorescence intensity) of the samples and the standard organism (Solanum pseudocapsicum, 1.295pg/1C).Supports Figure 3.*

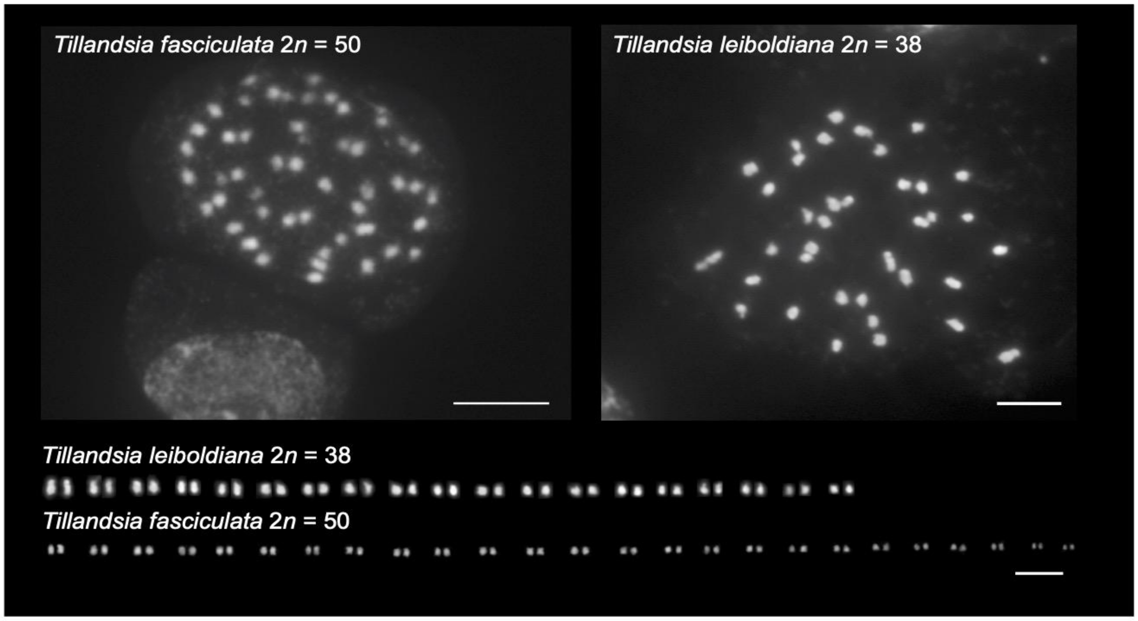


*Figure S16: Mitotic metaphase chromosomes and karyotypes of* Tillandsia fasciculata *and* Tillandsia leiboldiana*. Scale bar, 5µm. Supports Figure 3.*

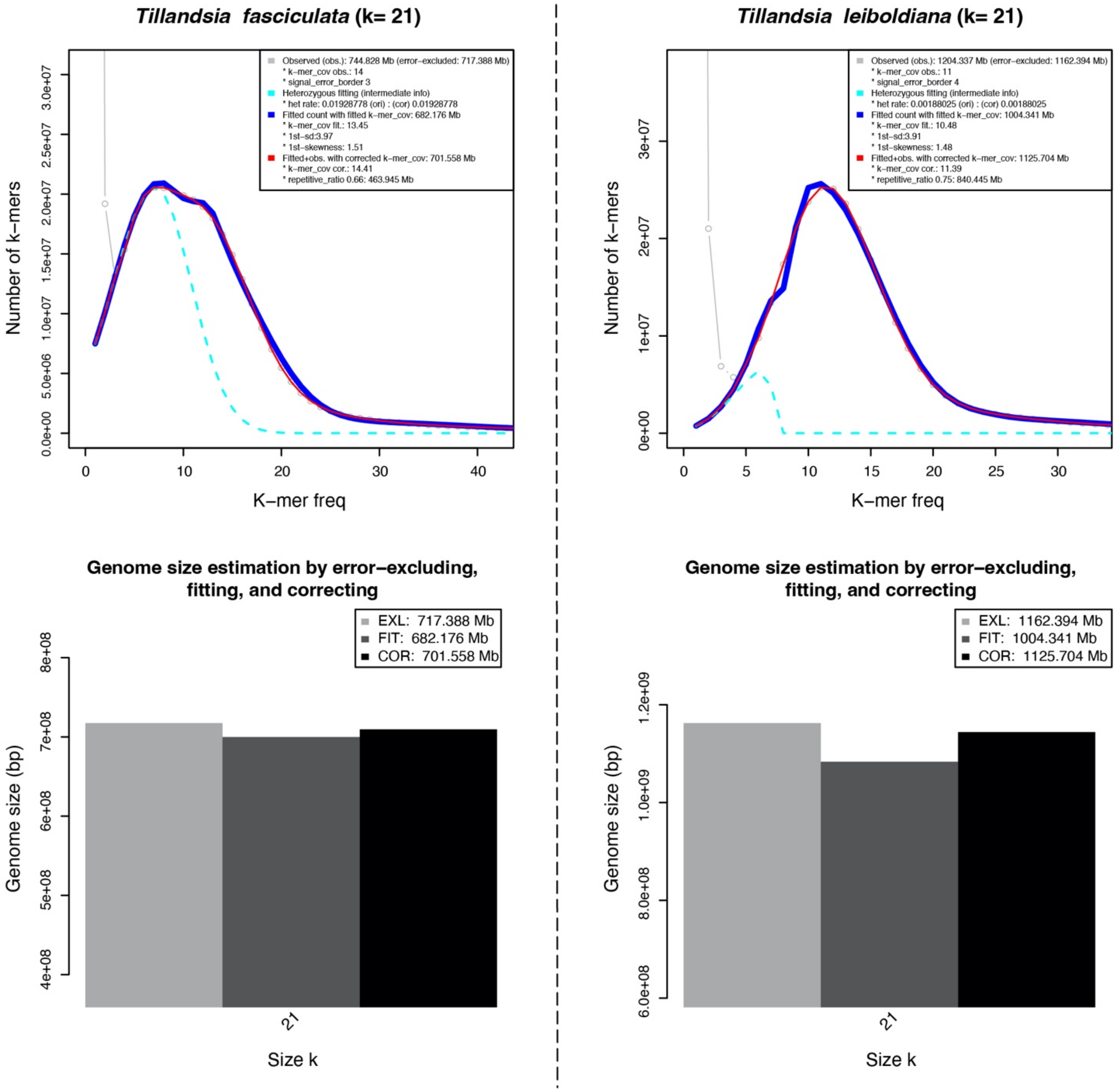


*Figure S17: Heterozygosity and genome size estimation with a k-mer based approach implemented in findGSE for* Tillandsia fasciculata *(left)* and T. leiboldiana *(right). Support Figure 3.*

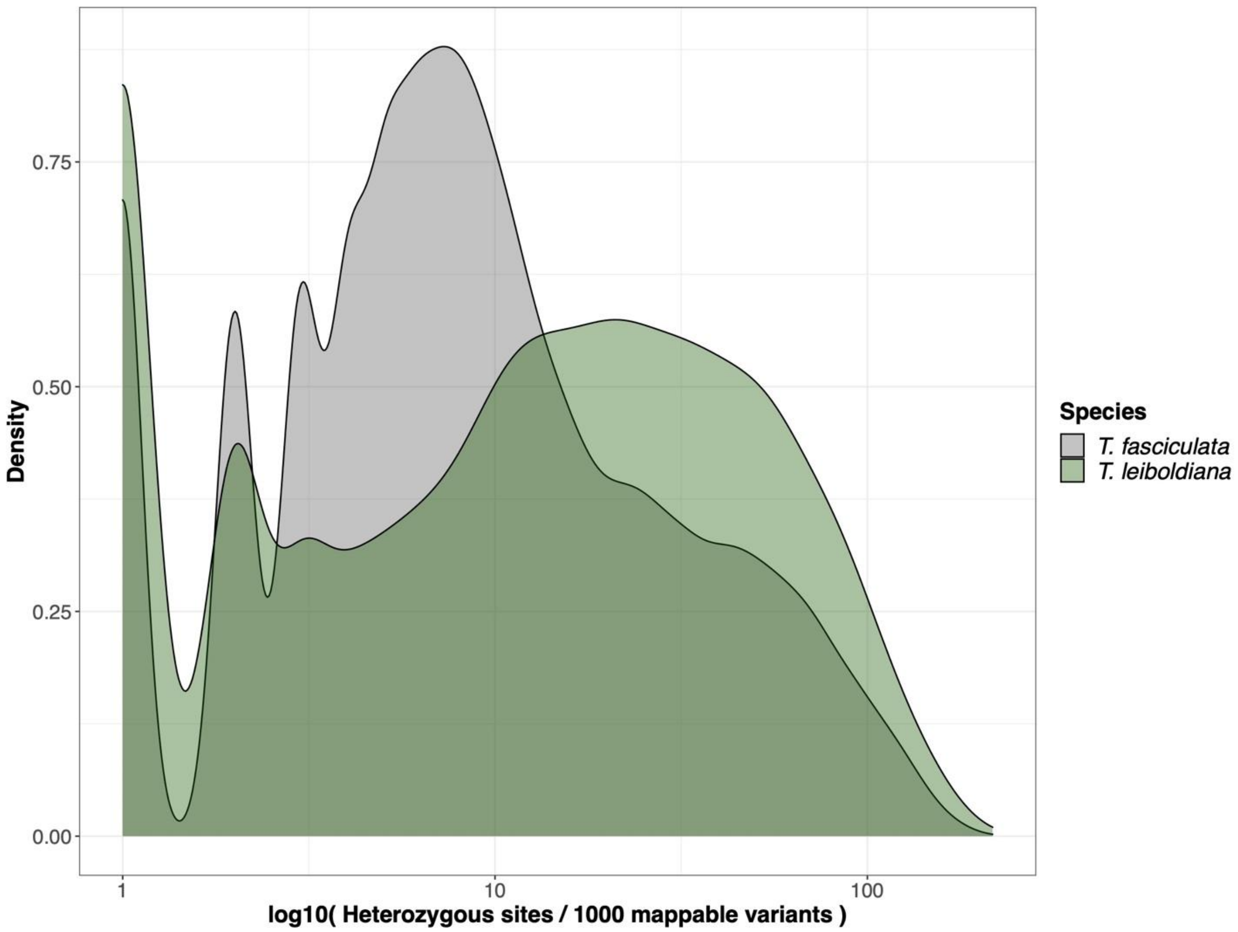


*Figure S18: Distribution of heterozygous sites per 1000 mappable variants on a logarithmic scale. Supports Figure 3.*

### 2. Supplementary tables

Table S1: List of accessions used in this study. Abbreviations used in this table: WU = Institutional Code (Wien Universität), HBV = Hortus Botanicus Vindobonensis (Botanical Garden of the University of Vienna), DBG = Deutsche Bromelien-Gesellschaft (German Bromeliad Society), s.n. = sin nomero (without number), s.d. = sin datos (without data), s.l. = sensu lato

| Source | Accession code | Herbarium accession | DNA № | Collector | Species | Country | Locality | Comment |
|---|---|---|---|---|---|---|---|---|
| Genome Assembly | | | | | | | | |
| Botanical Garden of the Univiersity of Vienna | HBV 0024657 (B179/91) | WU 0013642 | MHJB-B1840 | W. & S. Till 7116 | Tillandsia fasciculata s.l. | Costa Rica | Prov. Puntarenas, SW declivities of Cordillera de Tilaran, along the road from Sta. Elena to Rancho Grande. | |
| Botanical Garden of the University of Vienna | HBV 0024715 (B82/91) | WU 0003058 | MHJB-B1842 | W. & S. Till 7112 | Tillandsia leiboldiana s.l. | Costa Rica | Prov. Alajuela, 2 km N San Ranion | |
| RNA-SEQ for Gene Annotation | | | | | | | | |
| Botanical Garden of the University of Vienna | HBV 0026950 (B103/94) | WU 0006321 | MHJB-B1839 | G. Noller 9106 | Tillandsia fasciculata | Guatemala | Depto. Sololá, Lago de Atitlán | Originally sequenced for De La Harpe et. al. (2020) |
| Botanical Garden of the Univiersity of Vienna | HBV 0024656 (B108/94) | WU 0002124 | MHJB-B360 | K.-D. & R. Ehlers EM890701 | Tillandsia fasciculata | Mexico | Estdo. Chiapas, Sumidero Cañon bei Tuxtla Gutierrez | Originally sequenced for De La Harpe et. al. (2020) |
| Botanical Garden of the Univiersity of Vienna | HBV 0025322 (B100/91) | WU 0013632 | MHJB-B2295 | W. & S. Till 7005 | Tillandsia fasciculata | Costa Rica | Prov. Heredia, Barva north of Heredia | Originally sequenced for De La Harpe et. al. (2020) |
| Botanical Garden of the University of Vienna | HBV 0000663 (B84/91, BRO000613) | WU 0001725, WU 0003008 | MHJB-B323 | W. & S. Till 7043 | Tillandsia leiboldiana | Costa Rica | Prov. Cartago, Turrialba, Centro Agronomico Tropical de Investigacion y Enseñanza (CATIE) | |
| RNA-SEQ for Time Course Experiment | | | | | | | | |
| Botanical Garden of the University of Vienna | HBV 0025322 (B100/91) | s.d. | MHJB-B2295 | W. & S. Till 7005 | Tillandsia fasciculata | Costa Rica | Prov. Heredia, Barva north of Heredia | Tfas_E in RNA-seq analysis |
| Botanical Garden of the Univiersity of Vienna | HBV 0025194 (B293/96) | s.d. | MHJB-B2296 | S. Schatzl 77/59 | Tillandsia fasciculata | Mexico | Estdo. Jalisco, ca. 20 km S of Porto Vall | Tfas_B in RNA-seq analysis |
| Botanical Garden of the Univiersity of Vienna | HBV 0025334 (B99B53-1) | WU 0008562, WU 0013708 | MHJB-B1838 | E. Kamm s.n. | Tillandsia fasciculata | Honduras | s.d. | Tfas_C in RNA-seq analysis |
| Botanical Garden of the Univiersity of Vienna | HBV 0024655 (B99/91) (90/91) | WU 0013632, WU 0013754 | MHJB-B1841 | W. & S. Till 7004 | Tillandsia fasciculata | Costa Rica | Prov. Heredia, Barva north of Heredia | Tfas_D in RNA-seq analysis |
| Botanical Garden of the Univiersity of Vienna | HBV 0025326 (B90/91) | s.d. | MHJB-B2297 | W. & S. Till 7006 (7004) | Tillandsia fasciculata | Costa Rica | Prov. Heredia, Barva north of Heredia | Tfas_A in RNA-seq analysis |
| Botanical Garden of the Univiersity of Vienna | HBV 0024657 (B179/91) | WU 0013642 | MHJB-B1840 | W. & S. Till 7116 | Tillandsia fasciculata s.l. | Costa Rica | Prov. Puntarenas, SW declivities of Cordillera de Tilaran, along the road from Sta. Elena to Rancho Grande. | Tfas_F in RNA-seq analysis |
| Botanical Garden of the University of Vienna (Coll.Bak BV) | HBV 0032437 (Bak 126) | s.d. | MHJB-B1960 | s.d. (DBG Sept. 2011) | Tillandsia leiboldiana | Costa Rica | above Carteso | Tlei_A in RNA-seq analysis |
| Botanical Garden of the University of Vienna (Coll.Bak BV) | HBV 0032433 (Bak 27) | s.d. | MHJB-B1957 | P. Bak s.n. | Tillandsia leiboldiana | Costa Rica | s.d. | Tlei_C in RNA-seq analysis |
| Botanical Garden of the Univiersity of Vienna | HBV 0024715 (B82/91) | WU 0003058 | MHJB-B1842 | W. & S. Till 7112 | Tillandsia leiboldiana s.l. | Costa Rica | Prov. Alajuela, 2 km N San Ranion | Tlei_D in RNA-seq analysis |
| Botanical Garden of the University of Vienna (Coll.Bak BV) | HBV 0032436 (Bak 119) | s.d. | MHJB-B1959 | s.d. (DBG Sept. 2011) | Tillandsia leiboldiana | Mexico | Estdo. Puebla, Amixtlán | Tlei_E in RNA-seq analysis |
| Botanical Garden of the University of Vienna (Coll.Bak BV) | HBV 0032434 (Bak 37) | s.d. | MHJB-B1958 | P. Bak s.n. | Tillandsia leiboldiana | Costa Rica | s.d. | Tlei_F in RNA-seq analysis |
| Botanical Garden of the University of Vienna (Coll.Bak BV) | HBV 0032435 (Bak 45) | s.d. | MHJB-B1956 | P. Bak s.n. | Tillandsia leiboldiana | Honduras | s.d. | Tlei_G in RNA-seq analysis |

Table S2: Summary statistics of the *T. fasciculata* and *T. leiboldiana* assembly and gene annotation

| Assembly statistics | *T. fasciculata* | *T. leiboldiana* |
|---|---|---|
| Total length (bp) | 837,577,910 | 1,198,225,148 |
| Total scaffold count (> 1 kb) | 2,321 | 10,433 |
| Total contig count | 8,625 | 20,447 |
| N50 (Mb) | 23,642 | 43,365 |
| N90 (Kb) | 145,438 | 27,985 |
| L50 | 16 | 12 |
| L90 | 565 | 2,898 |
| GC content | 42.81 % | 44.73 % |
| Uniquely mapping RNA-seq reads | 69.13 % | 92.37 % |
| Complete BUSCO genes | 91.8 % | 88.1 % |
| Duplicated BUSCO genes | 6.2 % | 1.9 % |
| Fragmented BUSCO genes | 5.2 % | 5.4 % |
| **Gene model statistics** | *T. fasciculata* | *T. leiboldiana* |
| Gene model count | 34,886 | 38,180 |
| Average length (bp) | 4,090 | 4,225 |
| Complete BUSCO genes | 89.7 % | 85.3 % |
| Duplicated BUSCO genes | 11.6 % | 6.5 % |
| Fragmented BUSCO genes | 5.2 % | 7.9 % |
| Uniquely mapping RNA-seq reads | 64.61 % | 84.76 % |
| Gene models with AED-score > 0.5 | 93 % | 89.9 % |
| Scaffolds containing gene models | 1,191 | 2,621 |
| **Statistics of functional annotation** | *T. fasciculata* | *T. leiboldiana* |
| Gene models with BLAST | 31,883 | 33,971 |
| Gene models with GO terms | 26,505 | 27,148 |
| Gene models with Blast2Go annotation | 24,319 | 24,633 |

Table S3: Abundances of LTR, TIR and Helitron classes in main contigs of *T. fasciculata* and *T. leiboldiana*

| | | *Tillandsia fasciculata* | | | *Tillandsia leiboldiana* | | |
|---|---|---|---|---|---|---|---|
| Class | Type | Element count | Total length (bp) | Proportion of genome | Element count | Total length (bp) | Proportion of genome |
| | Total | 692,254 | 392,992,866 | 65.52 % | 1,268,380 | 697,969,745 | 77.07 % |
| LTR | | | | | | | |
| | Copia | 85,077 | 72,241,681 | 12.04 % | 226,304 | 117,239,901 | 12.95 % |
| | Gypsy | 136,750 | 130,003,938 | 21.67 % | 243,366 | 215,847,337 | 23.84 % |
| | Unknown | 179,371 | 110,637,301 | 18.45 % | 357,701 | 222,072,889 | 24.52 % |
| | Total | 401,198 | 312,882,920 | 52.16 % | 827,371 | 555,160,127 | 61.31 % |
| TIR | | | | | | | |
| | CACTA | 26,027 | 6,822,973 | 1.14 % | 22,788 | 6,785,096 | 0.75 % |
| | Mutator | 78,405 | 20,912,187 | 3.49 % | 227,557 | 87,286,075 | 9.64 % |
| | PIF_Harbinger | 13,059 | 3,335,951 | 0.56 % | 14,124 | 3,580,678 | 0.4 % |
| | Tc1_Mariner | 3,435 | 648,896 | 0.11 % | 1,589 | 372,253 | 0.04 % |
| | hAT | 31,871 | 8,365,061 | 1.39 % | 21,951 | 6,587,992 | 0.73 % |
| | Total | 152,797 | 40,085,068 | 6.69 % | 288,009 | 104,612,094 | 11.56 % |
| Helitron | | | | | | | |
| | Helitron | 118,069 | 35,165,625 | 5.86 % | 130,790 | 33,461,880 | 3.7 % |

Table S4: Orthology statistics before and after curation (removal of plastid genes and size correction). Size correction was not applied to *A. comosus,* however, unique genes to this species were removed from the dataset.

| | Before curation | | | After curation | | |
|---|---|---|---|---|---|---|
| General and single-copy statistics | *A. comosus* | *T. fasciculata* | *T. leiboldiana* | *A. comosus* | *T. fasciculata* | *T. leiboldiana* |
| Number of genes assigned to orthogroups | 21,045 | 26,325 | 23,584 | 20,416 | 24,397 | 22,968 |
| Proportion of input sequences assigned | 78 % | 87.5 % | 75 % | - | - | - |
| Number of single-copy genes in a given species | 12,794 | 14,311 | 15,537 | 12,602 | 15,236 | 15,479 |
| Proportion | 52.5 % | 54.36 % | 65.88 % | 61.73 % | 62.45 % | 67.39 % |
| Number of single-copy orthologues (1:1:1) | 10,012 | 10,012 | 10,012 | 10,707 | 10,707 | 10,707 |
| Proportion | 47.57 % | 38.03 % | 42.45 % | 52.44 % | 43.89 % | 46.62 % |
| Number of single-copy orthologues in *Tillandsia* | - | 13,128 | 13,128 | - | 14,086 | 14,086 |
| Proportion | - | 49.87 % | 55.66 % | - | 57.74 % | 61.33 % |
| Multi-copy statistics | | | | | | |
| Number of multi-copy genes in a given species | 8,260 | 12,014 | 8,011 | 7,809 | 9,161 | 7,489 |
| Proportion | 39.25 % | 45.64 % | 33.97 | 38.25 % | 37.55 % | 32.6 % |
| Number of genes in orthogroups with family size *T. fas* > *T. lei* | 2,594 | 6,976 | 2,709 | 1,178 | 4,170 | 1,356 |
| Proportion | 12.33 % | 26.50 % | 11.49 % | 5.77 % | 17.09 % | 5.9 % |
| Number of genes in orthogroups with family size T. fas < T. lei | 817 | 905 | 2,222 | 725 | 833 | 2,079 |
| Proportion | 3.88 % | 3.44 % | 9.42 % | 35.51 % | 34.14 % | 9.05 % |
| Number of genes in orthogroups with family size T. fas = T. lei | 1,244 | 1,265 | 1,265 | 1,291 | 1,258 | 1,258 |
| Proportion | 5.91 % | 4.81 % | 5,36 % | 6.32 % | 4.6 % | 4.6 % |
| Unique gene statistics | | | | | | |
| Number of unique genes | - | 3,101 | 3,192 | - | 3,051 | 3,154 |
| Proportion | - | 14.74 % | 13.53 % | - | 12.51 % | 13.73 % |

Table S5: List of CAM-related expanded orthogroups in either *T. fasciculata* or *T. leiboldiana.* Orthogroups marked by an asterisk (*) contain genes recovered in DGE analyses.

| Orthogroup | Conformation of gene family size | Function | Description |
|---|---|---|---|
| Expanded in *T. fasciculata* | | | |
| OG0000267* | 6 : 5 :4 | Pyrophosphate-energized vacuolar membrane | Proton pump located on the tonoplast that energizes the night-time accumulation of malate in |
| OG0000469 | 0 : 15 : 1 | Cytosolic enolase 3 | Catalyses the conversion of 2-phosphoglycerate (2-PG) to phosphoenolpyruvate (PEP) in the |
| OG0000539*, OG0004427 | 2 : 8 : 3, 1 : 2 :1 | Vacuolar-type proton ATPase subunit H | Subunit of a V-ATPase proton pump, which energizes the night-time accumulation of malate |
| OG0000555 | 0 : 7 : 1 | Enolase | Catalyses the conversion of 2-phosphoglycerate (2-PG) to phosphoenolpyruvate (PEP) in the |
| OG0000601* | 3 : 3 : 2 | V-type proton ATPase 16 kDa proteolipid | Subunit of a V-ATPase proton pump, which energizes the night-time accumulation of malate |
| OG0001440 | 2 : 2 : 1 | Succinate dehydrogenase [ubiquinone] | Subunit of the Succinate-ubiquinone oxidoreductase complex (complex II), which is |
| OG0001710 | 1 : 2 : 1 | Acidic beta-fructofuranosidase 1 | Vacuolar acid invertase involved in the accumulation of hexoses in the vacuole during the |
| OG0002059 | 2 : 2 : 1 | Malate dehydrogenase (MDH) | Converts oxaloacetate to malate as part of the carbon fixation module of CAM. |
| OG0002712, OG0003619 | 2 : 2 : 1, 1 : 2 : 1 | Late embryogenesis abundant (LEA) protein | Involved in drought stress response [4]. |
| OG0003207 | 1 : 3 : 1 | Beta carbonic anhydrase 5 | Involved in carbon fixation by catalyzing the conversion of CO2 to bicarbonate, which is the |
| OG0003437* | 1 : 2 : 1 | succinate dehydrogenase subunit 6, | Plant-specific [6] subunit of the succinate dehydrogenase complex involved in anchoring the |
| OG0005044* | 1 : 2 : 1 | Protein XAP5 CIRCADIAN TIMEKEEPER | Involved in the regulation of light response [8], the circadian clock [9], and disease resistance |
| OG0005172* | 1 : 2 : 1 | mitochondrial NAD-dependent isocitrate | Catalyzes the third step of the tricarboxylic acid cycle and is a putative alternative carbon |
| OG0000580* | 4 : 3 : 2 | Pyrophosphate--fructose 6-phosphate 1- | Catalyzes the second step of the glycolysis in the cytoplasm, and can generally mostly access |
| OG0001599* | 3 : 2 : 1 | granule-bound starch synthase | Involved in starch synthesis by catalyzing the production of amylose and the structure of |
| Expanded in *T. leiboldiana* | | | |
| OG0000288 | 2 : 1 : 13 | Histidine kinase 3 | Regulator of stomatal opening in *Arabidopsis thaliana* in response to light and stress [13] |
| OG0000320 | 4 : 4 : 5 | glyceraldehyde-3-phosphate dehydrogenase 2, | Catalyzes the sixth step of the glycolysis. |
| OG0000323 | 6 : 2 : 4 | Aquaporin NIP1-1 | Membrane transporter of water and small solutes, which shows diel expression in pineapple |
| OG0000507 | 7 : 1 : 2 | NADP-dependent isocitrate dehydrogenase | Catalyzes the third step of the tricarboxylic acid cycle. |
| OG0001563 | 3 : 1 : 2 | Enolase | Catalyses the conversion of 2-phosphoglycerate (2-PG) to phosphoenolpyruvate (PEP) in the |
| OG0001933 | 2 : 1 : 2 | Phosphoglycerate kinase | Catalyses the first energy-producing step of the glycolysis, and also the reversed step in |
| OG0004307 | 1 : 1 : 2 | Late embryogenesis abundant (LEA) protein | Involved in drought stress response [4]. |
| OG0005047* | 1 : 1 : 2 | Aquaporin PIP2-6 | Membrane transporter of water and small solutes, which shows diel expression in pineapple |


Citations

| | |
|---|---|
| [1] | Wai, C.M., VanBuren, R., Zhang, J., Huang, L., Miao, W., Edger, P.P., Yim, W.C., Priest, H.D., Meyers, B.C., Mockler, T., Smith, J.A.C., Cushman, J.C. and Ming, R. (2017), Temporal and spatial transcriptomic and microRNA dynamics of CAM photosynthesis in pineapple. Plant J, 92: 19-30. https://doi.org/10.1111/tpj.13630 |
| [2] | Araújo WL, Nunes-Nesi A, Osorio S, et al. Antisense inhibition of the iron-sulphur subunit of succinate dehydrogenase enhances photosynthesis and growth in tomato via an organic acid-mediated effect on stomatal aperture. Plant Cell. 2011;23(2):600-627. doi:10.1105/tpc.110.081224 |
| [3] | J. T. Christopher , JAM. Holtum, Patterns of Carbon Partitioning in Leaves of Crassulacean Acid Metabolism Species during Deacidification, Plant Physiology, Volume 112, Issue 1, September 1996, Pages 393–399, https://doi.org/10.1104/pp.112.1.393 |
| [4] | Xiao B, Huang Y, Tang N, Xiong L. Over-expression of a LEA gene in rice improves drought resistance under the field conditions. Theor Appl Genet. 2007;115(1):35-46. doi:10.1007/s00122-007-0538-9 |
| [5] | Ming, R., VanBuren, R., Wai, C. et al. The pineapple genome and the evolution of CAM photosynthesis. Nat Genet 47, 1435–1442 (2015). https://doi.org/10.1038/ng.3435 |
| [6] | Millar AH, Eubel H, Jänsch L, Kruft V, Heazlewood JL, Braun HP. Mitochondrial cytochrome c oxidase and succinate dehydrogenase complexes contain plant specific subunits. Plant Mol Biol. 2004;56(1):77-90. doi:10.1007/s11103-004-2316-2 |
| [7] | Christine Schikowsky, Jennifer Senkler, Hans-Peter Braun, SDH6 and SDH7 Contribute to Anchoring Succinate Dehydrogenase to the Inner Mitochondrial Membrane in Arabidopsis thaliana , Plant Physiology, Volume 173, Issue 2, February 2017, Pages 1094–1108, https://doi.org/10.1104/pp.16.01675 |
| [8] | Ellen L. Martin-Tryon, Stacey L. Harmer, XAP5 CIRCADIAN TIMEKEEPER Coordinates Light Signals for Proper Timing of Photomorphogenesis and the Circadian Clock in Arabidopsis , The Plant Cell, Volume 20, Issue 5, May 2008, Pages 1244–1259, https://doi.org/10.1105/tpc.107.056655 |
| [9] | Liu, Lei, Xiaoyun Li, Li Yuan, Guofang Zhang, Hui Gao, Xiaodong Xu, and Hongtao Zhao. "XAP5 CIRCADIAN TIMEKEEPER specifically modulates 3'splice site recognition and is important for circadian clock regulation partly by alternative splicing of LHY and TIC." Plant Physiology and Biochemistry 172 (2022): 151-157. |
| [10] | Xu, Yong-Ju, Yang Lei, Ran Li, Ling-Li Zhang, Zhi-Xue Zhao, Jing-Hao Zhao, Jing Fan et al. "XAP5 CIRCADIAN TIMEKEEPER Positively Regulates RESISTANCE TO POWDERY MILDEW8. 1–Mediated Immunity in Arabidopsis." Frontiers in Plant Science 8 (2017): 2044. |
| [11] | Nadine Töpfer and others, Alternative Crassulacean Acid Metabolism Modes Provide Environment-Specific Water-Saving Benefits in a Leaf Metabolic Model, The Plant Cell, Volume 32, Issue 12, December 2020, Pages 3689–3705, https://doi.org/10.1105/tpc.20.00132 |
| [12] | Nancy Wieland Carnal , Clanton C. Black, Soluble Sugars as the Carbohydrate Reserve for CAM in Pineapple Leaves : Implications for the Role of Pyrophosphate:6-Phosphofructokinase in Glycolysis, Plant Physiology, Volume 90, Issue 1, May 1989, Pages 91–100, https://doi.org/10.1104/pp.90.1.91 |
| [13] | Marchadier E, Hetherington AM. Involvement of two-component signalling systems in the regulation of stomatal aperture by light in Arabidopsis thaliana. New Phytol. 2014;203(2):462-468. doi:10.1111/nph.12813 |
| [14] | Zhu, F., Ming, R. Global identification and expression analysis of pineapple aquaporins revealed their roles in CAM photosynthesis, boron uptake and fruit domestication. Euphytica 215, 132 (2 |

Table S6: Full list of candidate genes for adaptive sequence evolution.

| Orthogroup | Genes | dN/dS | dN | dS | adj p-value | Function |
|---|---|---|---|---|---|---|
| **One-to-one orthologues** | | | | | | |
| OG0002972 | Tfasc_v1.01130-RA, Tlei_v1.02341-RA | Inf | 0,0416 | 0,0004 | 0,00162 | jacalin-related lectin 3-like |
| OG0005000 | Tfasc_v1.24851-RA, Tlei_v1.26186-RA | Inf | 0,0247 | 0,0002 | 0,00628 | cucumber peeling cupredoxin-like |
| OG0006253 | Tfasc_v1.06027-RA, Tlei_v1.01410-RA | Inf | 0,0221 | 0,0002 | 0,00066 | mitochondrial prohibitin-3 |
| OG0009278 | Tfasc_v1.29917-RA, Tlei_v1.20972-RA | Inf | 0,0214 | 0,0002 | 0,00155 | chloroplastic Peroxiredoxin-2E-2 |
| OG0012770 | Tfasc_v1.15761-RA, Tlei_v1.22494-RA | Inf | 0,0318 | 0,0003 | 0,01374 | metallo-hydrolase/oxidoreductase superfamily protein |
| OG0011786 | Tfasc_v1.12421-RA, Tlei_v1.10008-RA | 9,2985 | 0,0189 | 0,002 | 0,00008 | U-box_domain-containing_protein |
| OG0008528 | Tfasc_v1.04577-RA, Tlei_v1.08412-RA | 8,17 | 0,0322 | 0,0039 | 0,01292 | hypothetical protein ACMD2_08159 |
| OG0015603 | Tfasc_v1.14595-RA, Tlei_v1.21364-RA | 8,0697 | 0,0228 | 0,0028 | 0,01575 | anaphase-promoting complex subunit CDC27 |
| OG0008124 | Tfasc_v1.15280-RB, Tlei_v1.22017-RA | 3,1536 | 0,0238 | 0,0075 | 0,01368 | uncharacterized protein LOC109710130 isoform X1 |
| OG0009004 | Tfasc_v1.16390-RA, Tlei_v1.06962-RA | 3,1379 | 0,0271 | 0,0087 | 0,01148 | Hydroquinone glycosyltransferase |
| OG0019176 | Tfasc_v1.13382-RA, Tlei_v1.18537-RA | 2,8534 | 0,0466 | 0,0163 | 0,00561 | uncharacterized protein LOC109718296 |
| OG0008977 | Tfasc_v1.16327-RA, Tlei_v1.06894-RA | 2,6204 | 0,0281 | 0,0107 | 0,00512 | glutamate receptor 2.8-like |
| OG0010014 | Tfasc_v1.26649-RA, Tlei_v1.28282-RA | 2,0042 | 0,0701 | 0,035 | 0,00603 | Glycerophosphodiester phosphodiesterase GDPDL7 |
| **1:1:2 orthologues** | | | | | | |
| OG0003849 | Tfasc_v1.03397-RA, Tlei_v1.09369-RA | Inf | 0,0093 | 0,0001 | 0,00697 | kinesin-like protein KIN-10C |
| | Tfasc_v1.03397-RA, Tlei_v1.09368-RA | 4,1684 | 0,0093 | 0,0022 | 0,09607 | |
| **1:2:1 orthologues** | | | | | | |
| OG0003795 | Tfasc_v1.16138-RA, Tlei_v1.06694-RA | 6,2573 | 0,0648 | 0,0104 | 0,02611 | Ubiquitin-conjugating enzyme E2 |
| | Tfasc_v1.16140-RA, Tlei_v1.06694-RA | 0,2438 | 0,0334 | 0,1368 | 0,03386 | |
| OG0004404 | Tfasc_v1.08653-RA, Tlei_v1.17093-RA | 3,9913 | 0,0945 | 0,0237 | 0,02787 | protein ECERIFERUM 1-like |
| | Tfasc_v1.21655-RA, Tlei_v1.17093-RA | 0,1733 | 0,0051 | 0,0297 | 0,00059 | |

Table S7: Description of gene clusters inferred in co-expression analyses from maSigPro

| Cluster | Number of genes | Genes of interest | GO terms of interest |
|---|---|---|---|
| 1 | 87 | Tfasc_v1.23066: protein LNK1-like;<br>Tfasc_v1.19779: Dicarboxylate transporter 1, chloroplastic;<br>Tfasc_v1.08739: mitochondrial uncoupling protein 5-like;<br>Tfasc_v1.00158: ABC transporter G family member 5-like;<br>Tfasc_v1.01823: Protein REVEILLE 1 [1];<br>Tfasc_v1.03880: Transcription factor PCF2 [2];<br>Tfasc_v1.06881*: protein LHY-like isoform X1 [2];<br>Tfasc_v1.09028: aluminum-activated malate transporter 9-like isoform X1 [1] | GO:1902356: oxaloacetate(2-) transmembrane transport;<br>GO:0071423: malate transmembrane transport;<br>GO:1902074: response to salt;<br>GO:0071472: cellular response to salt stress;<br>GO:1902584: positive regulation of response to water deprivation;<br>GO:1901002: positive regulation of response to salt stress;<br>GO:0015131: oxaloacetate transmembrane transporter activity;<br>GO:0015140: malate transmembrane transporter activity |
| 2 | 134 | Tfasc_v1.25154: V-type proton ATPase catalytic subunit A [3];<br>Tfasc_v1.11797: protein XAP5 CIRCADIAN TIMEKEEPER [3];<br>Tfasc_v1.21051: V-type proton ATPase subunit e1;<br>Tfasc_v1.15469*: Acyl-coenzyme A thioesterase [2];<br>Tfasc_v1.09221: Pyrophosphate--fructose 6-phosphate 1-phosphotransferase subunit alpha;<br>Tfasc_v1.12690*: pentatricopeptide repeat-containing protein At1g09900-like [2]; | GO:0033179: proton-transporting V-type ATPase, V0 domain;<br>GO:0047334: diphosphate-fructose-6-phosphate 1-phosphotransferase activity |
| 3 | 38 | Tfasc_v1.09150: protein HOMOLOG OF MAMMALIAN LYST-INTERACTING PROTEIN 5;<br>Tfasc_v1.03774: probable aquaporin PIP2-6 [4];<br>Tfasc_v1.14176*: long chain acyl-CoA synthetase 4-like [2];<br>Tfasc_v1.24696: V-type proton ATPase subunit H-like [5];<br>Tfasc_v1.25341: pyrophosphate-energized vacuolar membrane proton pump [3];<br>Tfasc_v1.28097: F-box/kelch-repeat protein SKIP25 [2];<br>Tfasc_v1.14652-RA: protein GIGANTEA [3] | GO:0007623: circadian rhythm;<br>GO:0010378: temperature compensation of the circadian clock;<br>GO:0046323: glucose import;<br>GO:0009637: response to blue light;<br>GO:0048578: positive regulation of long-day photoperiodism, flowering;<br>GO:0071482: cellular response to light stimulus |
| 4 | 99 | Tfasc_v1.09150: protein HOMOLOG OF MAMMALIAN LYST-INTERACTING PROTEIN 5;<br>Tfasc_v1.03774: probable aquaporin PIP2-6 [4];<br>Tfasc_v1.14176: long chain acyl-CoA synthetase 4-like [2];<br>Tfasc_v1.24696: V-type proton ATPase subunit H-like [5];<br>Tfasc_v1.25341: pyrophosphate-energized vacuolar membrane proton pump [3];<br>Tfasc_v1.28097: F-box/kelch-repeat protein SKIP25 [2] | GO:1903335: regulation of vacuolar transport;<br>GO:0000221: vacuolar proton-transporting V-type ATPase, V1 domain |
| 5 | 209 | Tfasc_v1.01733: V-type proton ATPase 16 kDa proteolipid subunit [2];<br>Tfasc_v1.16595: soluble starch synthase I;<br>Tfasc_v1.27353: beta-amylase 3 [10]<br>Tfasc_v1.26941: glucose-6-phosphate/phosphate translocator [6];<br>Tfasc_v1.17464: fructose-bisphosphate aldolase 1, cytoplasmic [6]<br>Tfasc_v1.21461: Pyruvate kinase, cytosolic isozyme [2][6];<br>Tfasc_v1.16311: Phosphoenolpyruvate carboxylase (PEPC) [7];<br>Tfasc_v1.03126: malate dehydrogenase [7];<br>Tfasc_v1.03128: PEPC kinase [7];<br>Tfasc_v1.06749: phosphoglucan phosphatase DSP4, amyloplastic isoform X1;<br>Tfasc_v1.09378: V-type proton ATPase subunit d2 [2][5];<br>Tfasc_v1.20370: ATP-dependent 6-phosphofructokinase 5, chloroplastic;<br>Tfasc_v1.24086: ATP-dependent 6-phosphofructokinase 3;<br>Tfasc_v1.17156: aconitate hydratase;<br>Tfasc_v1.04514: glucose-6-phosphate isomerase, cytosolic;<br>Tfasc_v1.07899: phosphoenolpyruvate carboxykinase (ATP);<br>Tfasc_v1.26280: protein MAEA homolog;<br>Tfasc_v1.27863: pyruvate decarboxylase 1;<br>Tfasc_v1.03307: V-type proton ATPase subunit a3 [5];<br>Tfasc_v1.00598: nuclear pore complex protein NUP50A-like [2];<br>Tfasc_v1.01511: late embryogenesis abundant protein Lea5-like [8];<br>Tfasc_v1.03613*: probable pyruvate, phosphate dikinase regulatory protein, chloroplastic [2][7];<br>Tfasc_v1.14299: late embryogenesis abundant protein group 8 protein [8];<br>Tfasc_v1.14724: V-type proton ATPase subunit B 2 [3];<br>Tfasc_v1.15525: protein YLS7-like [2];<br>Tfasc_v1.16080: V-type proton ATPase subunit c''2 [2][5];<br>Tfasc_v1.17712: V-type proton ATPase subunit C [3];<br>Tfasc_v1.18209: V-type proton ATPase subunit G 1-like [3];<br>Tfasc_v1.19718: F-box protein At1g55000 [2];<br>Tfasc_v1.19742: V-type proton ATPase subunit H [5]<br>Tfasc_v1.26860: gamma-aminobutyrate transaminase 1, mitochondrial [2];<br>Tfasc_v1.28537: protein FD [9];<br>Tfasc_v1.28610: uncharacterized protein LOC109727440 [9] | GO:0005983: starch catabolic process;<br>GO:0061615: glycolytic process through fructose-6-phosphate;<br>GO:0006099: tricarboxylic acid cycle;<br>GO:0006094: gluconeogenesis;<br>GO:0006002: fructose 6-phosphate metabolic process;<br>GO:0045721: negative regulation of gluconeogenesis;<br>GO:0007035: vacuolar acidification;<br>GO:0006107: oxaloacetate metabolic process;<br>GO:0004737: pyruvate decarboxylase activity;<br>GO:0004612: phosphoenolpyruvate carboxykinase (ATP) activity;<br>GO:0008964: phosphoenolpyruvate carboxylase activity;<br>GO:0046961: proton-transporting ATPase activity, rotational mechanism;<br>GO:0047780: citrate dehydratase activity;<br>GO:0004347: glucose-6-phosphate isomerase activity |
| 6 | 144 | Tfasc_v1.20354: sucrose transport protein SUT2-like [3];<br>Tfasc_v1.25107: VMA21-like domain-containing protein;<br>Tfasc_v1.06152*: Protein EDS1L [2];<br>Tfasc_v1.08627: WAT1-related protein [2];<br>Tfasc_v1.16433: arabinogalactan peptide 16-like [2];<br>Tfasc_v1.24231*: replication stress response regulator SDE2 [2];<br>Tfasc_v1.30364*: protein FLX-like 4 [2];<br>Tfasc_v1.25107: VMA21-like domain-containing protein;<br>Tfasc_v1.26899: obg-like ATPase 1;<br>Tfasc_v1.25528: granule-bound starch synthase | GO:0015770: sucrose transport;<br>GO:0015768: maltose transport;<br>GO:0070072: vacuolar proton-transporting V-type ATPase complex assembly;<br>GO:1901001: negative regulation of response to salt stress;<br>GO:0005364: maltose:proton symporter activity;<br>GO:0003985: acetyl-CoA C-acetyltransferase activity;<br>GO:0004373: glycogen (starch) synthase activity |
| 7 | 196 | Tfasc_v1.30467: isocitrate dehydrogenase [NAD] catalytic subunit 5, mitochondrial;<br>Tfasc_v1.21339: catalase isozyme 1;<br>Tfasc_v1.12645: flavonoid 3',5'-hydroxylase 2-like [2];<br>Tfasc_v1.25055*: berberine bridge enzyme-like 18 [2];<br>Tfasc_v1.29220: ABC transporter C family member 4-like [3] | GO:1902074: response to salt;<br>GO:1900034: regulation of cellular response to heat;<br>GO:0004449: isocitrate dehydrogenase (NAD+) activity |
| | * | Candidate gene for adaptive sequence evolution in CAM/C3 shifts reported in [2] | |

Citations


| | |
|---|---|
| [1] | Wai, Ching M. et al. 2017. “Temporal and Spatial Transcriptomic and MicroRNA Dynamics of CAM Photosynthesis in Pineapple.” The Plant Journal 92(1): 19–30. |
| [2] | De La Harpe, M. et al. 2020. "Genomic footprints of repeated evolution of CAM photosynthesis in a Neotropical species radiation." Plant, Cell and Environment 43(12): 2987-3001. |
| [3] | McClung, C. Robertson. 2006. “Plant Circadian Rhythms.” The Plant Cell 18(4): 792–803. |
| [4] | Vera-Estrella, R. et al. 2012. “Day/Night Regulation of Aquaporins during the CAM Cycle in Mesembryanthemum Crystallinum.” Plant, Cell & Environment 35(3): 485–501. |
| [5] | Cosentino, Cristian et al. 2013. “Proteomic Analysis of Mesembryanthemum Crystallinum Leaf Microsomal Fractions Finds an Imbalance in V-ATPase Stoichiometry during the Salt-Induced Transition from C3 to CAM.” Biochemical Journal 450(2): 407–15. |
| [6] | Cushman, John C. et al. 2008. “Large-Scale MRNA Expression Profiling in the Common Ice Plant, Mesembryanthemum Crystallinum, Performing C3 Photosynthesis and Crassulacean Acid Metabolism (CAM).” Journal of Experimental Botany 59(7): 1875–94. |
| [7] | Ming, Ray et al. 2015. “The Pineapple Genome and the Evolution of CAM Photosynthesis.” Nature Genetics 47(12): 1435–42. |
| [8] | Xiao, B., et al. 2007. “Over-Expression of a LEA Gene in Rice Improves Drought Resistance under the Field Conditions.” Theoretical and Applied Genetics 115(1): 35–46. |
| [9] | Christin, Pascal-Antoine et al. 2014. “Shared Origins of a Key Enzyme during the Evolution of C4 and CAM Metabolism.” Journal of Experimental Botany 65(13): 3609–21. |
| [10] | Sean E. Weise and others, The role of transitory starch in C3, CAM, and C4 metabolism and opportunities for engineering leaf starch accumulation, Journal of Experimental Botany, Volume 62, Issue 9, May 2011, Pages 3109–3118, https://doi.org/10.1093/jxb/err035 |

Table S8: Per-kb frequency of four promotor motifs associated with transcription factors of the circadian clock: The Morning Element (MOE), the Evening Element (EE), the CCA1-Binding Site (CBS) and the G-box motif. The difference in frequency reflects the change in the per-kb frequency of a motif in the target group compared to the background per-kb frequency (in percentage). Negative values reflect a decreased frequency of a motive compared to the background, where as positive values show an increase. Motifs with > 10 % frequency increase in a target group are shown in bold. Significance of per-gene motif frequency difference was tested with the Wilcoxon-Rank Test: *p-value < 0.05, **p-value < 0.01, ***p-value < 1E-03.

| Background per-kb frequency of circadian motifs | | | | | |
|---|---|---|---|---|---|
| Group | Species | MOE | EE | CBS | G-box |
| Non-DE genes | *T. leiboldiana* | 0,172479 | 0,117555 | 0,105135 | 0,193086 |
| | *T. fasciculata* | 0,161376 | 0,093558 | 0,105319 | 0,184369 |
| **Difference in per-kb frequency of circadian motifs in percentage (%)** | | | | | |
| | Species | MOE | EE | CBS | G-box |
| All DE genes | *T. leiboldiana* | -4,18 | **15,78** | 9,65 | -3,25 |
| | *T. fasciculata* | -2,93 | **18,54** | **18,39** | 0,93 |
| cluster 1 | *T. fasciculata* | -13,72 | 8,24 | 14,18 | **81,94**** |
| cluster 2 | *T. fasciculata* | 0,49 | 5,92 | -18,74 | -26,70 |
| cluster 3 | *T. fasciculata* | -15,50 | **207,70**** | **72,64** | -1,38 |
| cluster 4 | *T. fasciculata* | **25,98** | -0,16 | **35,64** | -19,54 |
| cluster 5 | *T. fasciculata* | -10,10 | **49,53** | **45,13** | -4,45 |
| cluster 6 | *T. fasciculata* | 1,59 | 0,75 | -49,41 | -17,75 |
| cluster 7 | *T. fasciculata* | -8,33 | -17,78 | **43,27*** | 12,33 |

Table S9: Occurrence of circadian TF-binding motifs in 2-kb (* and 3-kb) upstream regions of core CAM genes and their related homologs in *T. fasciculata* and *T. leiboldiana*. Genes marked with an ✦ were differentially expressed.

| Gene_ID | MOE | EE | CBS | G-Box | Orthogroup | Function |
|---|---|---|---|---|---|---|
| Tfasc_v1.29951-RA | 0 | 2 | 0 | 2 | OG0009306 | α-Carbonic anhydrase |
| Tlei_v1.20941-RA | 0 | 1 | 0 | 1 | | |
| Tfasc_v1.09713-RA | 0 | 1 | 0 | 0 | OG0008211 | β-Carbonic anhydrase 2 |
| Tlei_v1.24345-RA | 0 | 1 | 0 | 0 | | |
| Tfasc_v1.04038-RA | 0 | 1* | 0 | 0 | OG0003207 | β-Carbonic anhydrase 5 |
| Tfasc_v1.08735-RA | 0 | 0 | 0 | 1 | | |
| Tlei_v1.13295-RA | 0 | 0 | 0 | 1 | | |
| Tfasc_v1.07635-RA | 1 | 0 | 1* | 0 | OG0001548 | Malate dehydrogenase |
| Tfasc_v1.03126-RA ✦ | 1 | 0 | 0 | 0 | | |
| Tlei_v1.15644-RA | 1* | 0 | 1* | 0 | | |
| Tlei_v1.03837-RA ✦ | 1 | 0 | 0 | 0 | | |
| Tfasc_v1.16311-RA ✦ | 0 | 0 | 0 | 0 | OG0001504 | PEPC 1 |
| Tlei_v1.06877-RA ✦ | 0 | 0 | 0 | 0 | | |
| Tfasc_v1.09872-RA | 0 | 0 | 0 | 0 | | |
| Tlei_v1.24540-RA | 1 | 0 | 0 | 0 | | |
| Tfasc_v1.17028-RA | 0 | 1* | 0 | 1 | OG0014303 | PEPC 2 |
| Tlei_v1.07896-RA | 0 | 1 | 0 | 1 | | |
| Tfasc_v1.07899-RA ✦ | 3 | 0 | 0 | 0 | OG0019125 | PEPCK |
| Tlei_v1.15895-RA ✦ | 1* | 0 | 0 | 0 | | |
| Tfasc_v1.21696-RA | 1 | 0 | 0 | 0 | OG0004399 | PPCK 1 |
| Tlei_v1.17058-RA | 1 | 1 | 0 | 0 | | |
| Tfasc_v1.03128-RA ✦ | 0 | 0 | 1* | 0 | OG0013607 | PPCK 2 |
| Tlei_v1.03835-RA ✦ | 0 | 0 | 1 | 2 | | |

Table S10: List of DE genes related to CAM, starch metabolism and gluconeogenesis with a TE insertion rate higher than twice the genome-wide average in *T. leiboldiana* and *T. fasciculata.* Genes of the two species belonging to the same orthogroup are marked with the same symbol in the first column. When only one orthologous copy of the gene exists in the other species, we report the difference in the number of TE insertions between species as (# insertion in *T. fasciculata* - # insertion in *T. leiboldiana*). Therefore, a positive value reflects a gene with more TE insertions in *T. fasciculata* than *T. leiboldiana*, while a negative value shows the opposite.

| Gene ID | TE insertion count (intronic / intronic + 3 kb upstream) | Orthogroup | # copies in *A. comosus* | # copies in *T.fasciculata* | # copies in *T.leiboldiana* | ΔTE insertions (F - L) | Description |
|---|---|---|---|---|---|---|---|
| In T. fasciculata | | | | | | | |
| Tfasc_v1.24696-RA | 9 / 19 | OG0000539 | 2 | 8 | 3 | NA | V-type proton ATPase subunit H-like (proton pump, vacuolar transport) |
| Tfasc_v1.29220-RA | 6 / 6 | OG0001699 | 2 | 2 | 2 | NA | ABC transporter C family member 4-like (stomatal movement, circadian rhythm) [1][2] |
| Tfasc_v1.22969-RA ✦ | 10 / 10 | OG0011563 | 1 | 1 | 1 | 3 | glucose-1-phosphate adenylyltransferase large subunit 3, chloroplastic/amyloplastic (starch synthesis) |
| Tfasc_v1.30467-RA | 7 / 9 | OG0005172 | 1 | 2 | 1 | 4 | isocitrate dehydrogenase [NAD] catalytic subunit 5, mitochondrial (alternative $CO_2$ fixator in CAM) [3] |
| Tfasc_v1.08627-RA | 19 / 24 | OG0007038 | 1 | 1 | 1 | 19 | WAT1-related protein (secondary cell wall growth) [4] |
| Tfasc_v1.09028-RA | 10 / 11 | OG0007222 | 1 | 1 | 1 | 4 | aluminum-activated malate transporter 9-like isoform X1 (malate transporter) |
| Tfasc_v1.17712-RA ✣ | 7 / 19 | OG0004832 | 2 | 1 | 1 | -1 | V-type proton ATPase subunit C (proton pump, vacuolar transport) |
| Tfasc_v1.12645-RA ╳ | 7 / 10 | OG0016000 | 1 | 1 | 1 | 0 | flavonoid 3',5'-hydroxylase 2-like (anthocyanin pathway) [5] |
| Tfasc_v1.14652-RA ✭ | 12 / 16 | OG0014957 | 1 | 1 | 1 | -2 | protein GIGANTEA (circadian rhythm, stomatal opening) [7][8] |
| In T. leiboldiana | | | | | | | |
| Tlei_v1.10209-RA ╳ | 7 / 19 | OG0016000 | 1 | 1 | 1 | 0 | flavonoid 3',5'-hydroxylase 2-like (anthocyanin pathway) [5] |
| Tlei_v1.21413-RA ✭ | 14 / 16 | OG0014957 | 1 | 1 | 1 | -2 | protein GIGANTEA (circadian rhythm, stomatal opening) [7][8] |
| Tlei_v1.16158-RA | 23 / 36 | OG0015868 | 1 | 1 | 1 | -14 | Beta-fructofuranosidase, insoluble isoenzyme 3 (sugar metabolism) |
| Tlei_v1.04787-RA ✣ | 8 / 18 | OG0004832 | 2 | 1 | 1 | -1 | V-type proton ATPase subunit C (proton pump, vacuolar transport) |
| Tlei_v1.02834-RA | 15 / 25 | OG0002339 | 2 | 1 | 1 | 1 | sugar transporter ERD6-like |
| Tlei_v1.23795-RA ✦ | 7 / 7 | OG0011563 | 1 | 1 | 1 | 3 | glucose-1-phosphate adenylyltransferase large subunit 3, chloroplastic/amyloplastic (starch synthesis) |

### 3. Supplementary Notes

#### Note 1: Genome size and karyotype of *T. fasciculata* and *T. leiboldiana*

The genome size of both specimens used for *de novo* assembly was measured with flow cytometry (See Methods, section 5.1.1.), estimating genome sizes of approximately 790 and 1,130 Mb for *T. fasciculata* and *T. leiboldiana* respectively (Fig. S15). These estimates are slightly higher than those obtained computationally with a kmer-based approach implemented in findGSE (Sun et al., 2018) (k = 21 ,701 and 1,125 Mb respectively, Fig. S17); but deviations of genome size in computational approaches have been reported frequently (Al-Qurainy et al., 2021; Pflug et al., 2020; Elliott and Gregory, 2015). We also obtained a karyotype for both species using root material (See Methods, section 5.1.2.). We observe a change in karyotype between the two species resulting in a reduction by six chromosome pairs in *T. leiboldiana* compared to *T. fasciculata*, which carries the base karyotype of 2n = 50 encountered in *Tillandsioideae* (Brown and Gilmartin, 1989) (Fig. S16). This is in accordance with chromosome counts reported by Brown and Gilmartin (1989).

#### Note 2: Pre-assembly estimation of per-accession heterozygosity

Heterozygosity estimates of the chosen accessions along with several other candidate accessions from the Botanical Gardens of the University of Vienna were obtained with short-read Illumina data before *de novo* assembly, with the aim to select accessions with lowest heterozygosity and to make adjustments during *de novo* assembly to account for potentially elevated rates of heterozygosity. This short-read data was later used for polishing purposes and the sequencing details can be found in the Methods section 5.2.1.

Heterozygosity estimates were obtained by two approaches: a k-mer based approach and a reference-based approach. For the k-mer based approach, findGSE (Sun et al., 2018) was used with a k = 21 to obtain k-mer peaks of heterozygosity. For the reference-based approach, reads were trimmed with TrimmOmatic (Bolger et al., 2014) and mapped with GSNAP (Wu et al., 2016) to the *Tillandsia adpressiflora* pseudoreference built by De la Harpe and colleagues (2020). After filtering for low mapping quality and marking duplicates, variants were called for all accessions using *freebayes* (Garrison and Marth, 2012). Variants with an individual depth under 5 and above 45 were removed and no missing data was retained. Then, heterozygous sites per 1000 mappable sites were counted with a custom-made python script, filtering for allele balance between 0.25 and 0.75. This yielded between 51,000 and 58,000 windows, translating to roughly a 50 Mb portion of the genome. We expect these windows to be enriched for genic and other conserved regions, therefore resulting in an underestimate, though for relative comparisons, we regard this approach as valid.

Both the k-mer (Fig. S17) and reference-based approach (Fig. S18) showed that heterozygosity is elevated in *T. fasciculata* compared to *T. leiboldiana*. This result is consistent with the nucleotide diversity estimates reported by (Yardeni et al., 2021) based on sequencing of 1776 targeted loci in several *Tillandsia* species, including *T. leiboldiana* ($\pi_S$=5.7x10$^{-3}$) and *T. fasciculata* ($\pi_S$=8.1x10$^{-3}$). The relatively moderate levels of diversity in these species helped us to obtain assemblies with such a remarkable contiguity (Table S2), despite their very high repetitive content (see Supplementary Note 4).

**Note 3: Identifying main scaffolds in de novo assembly**

After scaffolding with Hi-C data, the resulting *de novo* assemblies contained a total of 2,321 and 10,443 scaffolds (> 1 kb) for *T. fasciculata* and *T. leiboldiana* respectively. However, more than 99 % of one-to-one orthologous gene pairs are located on the 25 and 26 largest scaffolds of both assemblies respectively, while 90.7 % and 87.6 % of all gene models are on these scaffolds. Therefore, the remaining scaffolds mainly consist of repetitive content, virtually corresponding to short, duplicated regions in the assembly. The mean proportion of repetitive content in the remaining scaffolds is indeed much higher than in main scaffolds (94.9 % and 91 % in small scaffolds of *T. fasciculata* and *T. leiboldiana*, versus 65.5 and 77.1% in the "main" scaffolds (see main text)). In addition, these "main" scaffolds contain the vast majority of the assembly (72 % and 75.5 % of the total assembly length). While the sizes of the longest 25 and 24 scaffolds are over 1 Mb, other scaffold sizes steeply decline afterwards (Fig. S2). Given this, we decided to regard these scaffolds as representative for the respective genomes and excluded all secondary scaffolds from downstream analyses from this point onwards. Though scaffolds 26 and 25 in *T.*

*leiboldiana* are smaller than 1 Mb, they contain a substantial number of orthologous genes (Fig. S2) and were therefore maintained in all analyses.

**Note 4: On the spatial distribution of GC and TE content in bromeliad genomes**

After finding that GC and genic content were negatively correlated in both *Tillandsia* genomes (See Results), we decided to study the link between GC content and repetitive content in all bromeliad genomes available to us at the time (*A. comosus*, *T. fasciculata* and *T. leiboldiana*) Using softmasked versions for all three genomes (see Materials and Methods), we computed the proportion of soft-masked bases across 100 kb windows. We also computed the overall GC content (considering both softmasked and non-softmasked positions), the GC content for soft-masked bases only, and the GC content for non-softmasked bases in the same windows. Based on this, the average TE content was estimated to be of 36.7 %, 67.9 % and 79.1% for *A. comosus*, *T. fasciculata* and *T. leiboldiana*, respectively. These three genomes therefore represent a gradient regarding the amount of TEs. After having reported the relatively well conserved synteny (Fig. 3c), we were able to estimate the evolution of the repetitive, GC and genic chromosomal landscapes across syntenic chromosomes.

We selected three examples of syntenic triplets considering scaffolds with no main chromosomal rearrangements: triplets A.com 3 / T.fas 4 / T.lei 1 (Fig. 3b), A.com 6 / T.fas 11 / Tlei. 15 and A.com 11 / T.fas 12 / T.lei 5 (Fig. S3). We then considered a relative position of each window on the scaffold (window position/scaffold length) to account for the difference in length of the syntenic chromosomes in the three assemblies. From this visualisation, it became clear that the GC content landscape is largely shaped by TE dynamics in the three genomes, since the GC% at non-repetitive content show no to little variation across the scaffold, except in *T. leiboldiana*. This contributed to building large GC-rich isochores in centromeric regions. Note here that our *de novo* TE libraries are not necessarily exhaustive and therefore a part of the repetitive content may have remained non-softmasked, which could partly or completely explain the pattern observed for GC at non-softmasked position in *T. leiboldiana*. Since this pattern is observed in all three species, regardless of the difference in repetitive content, this may be a family-wide phenomenon.

### Note 5: Identifying large-scale rearrangements between *T. fasciculata* and *T. leiboldiana*

Large-scale rearrangements were first perceived in the synteny analysis of the two assemblies. These were further investigated by whole-genome alignment of the two assemblies to each other, and to *A. comosus*, using nucmer (Delcher et al., 2002) and visualised with Dot (https://github.com/dnanexus/dot).

Large rearrangements that were identified between *T. fasciculata* and *T. leiboldiana* were further investigated by performing LastZ alignments (Harris, 2007) of soft-masked genomes. LastZ was run with the following settings: --notransition, --step=10, --gapped, --chain, --gfextend, --format=maf. Local alignments were filtered by the 95th identity and length quartile as implemented in (Leroy et al., 2021). Additionally, alignments were filtered by uniqueness with a custom-made python script, by removing all alignments with more than a 90 % overlap. Final local alignments were visualised for each scaffold as in Leroy et al (2021). All scripts are available at: https://shorturl.at/xLS15.

The breakpoint area of confirmed rearrangements was defined as the region between the last alignment of a given scaffold and the first alignment of another scaffold. Rearrangements were then finally confirmed by investigating the alignment of long-read PacBio data to the assembly of the same species. Whenever no clear break could be identified in the long-read alignment within the breakpoint area, the rearrangement was considered as confirmed.

With these methods, we described three potential large-scale rearrangements. Scaffold 14 in *T. leiboldiana* could be a fusion of scaffolds 17 and 25 in *T. fasciculata*, or scaffolds 17 and 25 could be the result of a fission in a reversed scenario (Fig. S4a). We also detected two potential translocations (Fig. S4b for Translocation 1 and Fig. S4c for Translocation 2). All breakpoints were confirmed by alignment of raw PacBio reads, except for breakpoint 1 on scaffold 13 of *T. fasciculata* of Translocation 1 (Fig. S4b). However, breakpoint 2 on this scaffold was confirmed, which led us to maintain the translocation as a candidate rearrangement. For several scaffolds, local alignments were too sparse to determine a clear breakpoint (See Fig. S4). To see the PacBio alignment at each breakpoint, see the supplementary PDF file "Tfas_Tlei_rearrangements.pdf" on our github repository.

We studied the effects of large-scale rearrangement on the genomic distribution of $d_N/d_S$ values and, separately, on DE genes (See SI Note 11) to understand if there is a link between chromosomal and functional evolution in *Tillandsia*. We did this by testing whether the distribution of $d_N/d_S$ values in any of the rearranged chromosomes deviated from that of non-rearranged chromosomes (See Methods, section 9.1). Of the nine scaffolds involved in the three reported rearrangements, only two had a $d_N/d_S$ distribution significantly deviating from that of non-rearranged chromosomes (scaffold 13 in *T. fasciculata* and scaffold 19 in *T. leiboldiana*, see Fig. S5b). The $d_N/d_S$ values in these scaffolds, which are both involved in Translocation 1, were ever so slightly reduced compared to non-rearranged scaffolds (median scaffold 13 = 0.3197, other scaffolds in *T. fasciculata* = 0.3569; median scaffold 19 = 0.3326, other scaffolds in *T. leiboldiana* = 0.3591). An overall reduction in chromosome-wide $d_N/d_S$ values can be expected in rearranged chromosomes due to an increase in linkage disequilibrium resulting from recombination suppression, which in turn increases background selection (Mérot et al., 2020; Cicconardi et al., 2021). This can have important implications for both adaptation and speciation, as functional loci may be under increased selective constraint, and selection against introgression may also become stronger (Cicconardi et al., 2021). However, the significance of this reduction in $d_N/d_S$ is very slight and only visible in two of nine rearranged scaffolds. Combined with our results on DE gene distribution across the genome, which show no signal of rearrangements playing a role in spatial distribution of ecologically relevant genes (SI Note 12), we are cautious in heralding large-scale rearrangement as a key driving force of ecological diversification in *Tillandsia* until additional supporting evidence becomes available (See SI Note 14).

**Note 6: Selecting rapidly evolving gene families**

To better understand the distribution of gene size differences, the log-ratio was taken of *T. fasciculata* to *T. leiboldiana* gene counts, and the overall mean log-ratio was subtracted to correct for background rates of gene loss or duplication. Orthogroups were ranked by corrected log-ratios and the top and bottom 2 % were then selected for further analysis. Due to the relatively large proportion of one-to-one relationships (79 %) among orthogroups, all orthogroups with a family size change between *T. fasciculata* and *T. leiboldiana* were included in the top 2 % of gene changes and therefore selected for GO term enrichment, which was performed separately for orthogroups with gene count larger in *T. fasciculata* (916 orthogroups) and larger in *T. leiboldiana* (583 orthogroups).

**Note 7: Detailed description of candidate genes for positive selection**

Our $d_N/d_S$ calculations pointed at 13 single-copy and 3 multi-copy genes exhibiting signatures of divergent selection between *T. fasciculata* and *T. leiboldiana*. The most relevant genes have been described in the main text (fig. 4c) and in Table S6, but here we provide more information on all candidate genes, which could be interesting for future work to investigate speciation genes in *Tillandsia*.

Among single-copy candidates, we found a Jacalin-related lectin (JLR, OG0002972), which are often associated with biotic and abiotic stimuli, though their biological function is largely unknown. In wheat, a mannose-specific JLR has been identified as a component of the defence system (Xiang et al., 2011). In rice, a JLR has been described as playing a role in salt stress response (Zhang et al., 2000).

Orthogroup OG0005000 codes for a cupredoxin (cupredoxin cucumber peeling-like). Cupredoxins are small proteins containing a copper centre which function as electron transfer shuttles between redox partners, but their more specific biological function is largely unknown (Guss et al., 1996). However, they tend to play a role in respiration, photosynthesis and metabolism (Choi and Davidson, 2011), and therefore represent another interesting candidate for further investigations.

Another candidate for adaptive sequence evolution was a mitochondrial prohibitin-3 (OG0006253), a subunit of the prohibitin complex. While the exact mechanism of prohibitin is unknown, it has been associated with mitochondrial biogenesis in *Nicotiana benthamiana* (Ahn et al., 2006) and more specifically protection against salt stress in *Arabidopsis thaliana* (Wang et al., 2012).

We recovered a hydroquinone glycosyltransferase (OG0009004), a broad-spectrum glycosyltransferase involved in the secondary metabolism of many phenolic compounds and xenobiotics(Hefner et al., 2002).

Orthogroup OG0009278 codes for chloroplastic Peroxiredoxin-2E-2, which is a member of the thiol peroxidase family. These enzymes play an important role in regulating Reactive

Oxygen Species (ROS) by reducing hydroperoxides. Peroxiredoxin-2E-2 is present in chloroplasts, especially in reproductive tissues, and its expression is sensitive to light and salt levels(Rouhier and Jacquot, 2005).

The remaining single-copy orthogroups that are candidates for adaptive sequence evolution are involved in cell replication (OG0015603) or members of a broad gene superfamily (OG0012770).

In addition to single copy genes, we tested for adaptive sequence evolution in orthogroups with a 1:1:2 or 1:2:1 relationship, *i.e.* a single gene in *A. comosus* and a duplicated gene either in *T. leiboldiana* (1:1:2 , 108 genes), or *T. fasciculata* (1:2:1, 190). We recovered a gene family in 1:2:1 conformation coding for a protein ECIFERUM 1-like, where one of two copies in *T. fasciculata* had $\omega > 1$. In *A. thaliana,* protein ECIFERUM-1 is involved in the biosynthesis of alkanes, which form hydrophobic cuticular waxes that protect the plant from desiccation. It has been shown that changes in expression of ECIFERUM-1 affect susceptibility to water stress and pathogens, therefore linking the protein with responses to biotic and abiotic stress(Bourdenx et al., 2011).

We also recovered one candidate orthogroup in 1:1:2 conformation coding for a kinesin-like protein KIN-10C. Members of the kinesin superfamily are molecular motors playing important roles in intracellular transport of vesicles and organelles, spindle formation and elongation, chromosome segregation, morphogenesis, and signal transduction (Li et al., 2012). Both *T. leiboldiana* gene copies showed elevated dN/dS ratios (See Table 1), though only one ratio was significant, suggesting that both gene copies have undergone significant evolution in *T. leiboldiana*.

Another candidate orthogroup in 1:2:1 conformation codes for a Ubiquitin-conjugating (UBC) enzyme E2, which plays an important role in the targeting of proteins by ubiquination for the proteasome. In mung bean, a UBC E2 enhances osmotic stress tolerance (Chung et al., 2013), and in Arabidopsis the overexpression of a soybean (Zhou et al., 2010) and peanut (Wan et al., 2011) UBC E2 protein increases drought and salt tolerance.

**Note 8: Choice of reference genome, mapping bias and mapping statistics for RNA-seq analysis**

Our RNA-seq dataset of 72 samples (2 species x 6 timepoints x 6 accessions) consists on average of 68,856,922 ± 15,234,149 reads per sample. Prior to conducting DGE analysis on the full set of RNA-seq reads, we tested for a potential mapping bias due to the choice of reference genome by mapping a random subset of reads consisting of 10 % of the original set of reads per sample, for a subset of 24 samples (3 samples per species, 4 time points per sample). Subsets were mapped to the *T. fasciculata*, *T. leiboldiana* and *A. comosus* (ASM154086v1) genome assemblies with STAR as described in Section 5.11.3 of the main text. Mapping statistics were collected from STAR's final log file (Log.final.out) and visualised in R with *ggplot2* (Fig. S13 and S14).

Unique mapping rates differ the least between species when mapping to A. comosus, however, a very large proportion (> 60 %) of reads are lost (Fig S13). When mapping to *T. fasciculata* reference genome, the difference in average mapping rate between species is 2,9 %, while being 5.6 % when mapping to *T. leiboldiana*. Also the difference between species in average multimapping rate is smaller for the *T. fasciculata* reference (0.88 % versus 1.38 % for the *T. leiboldiana* reference), despite having overall higher multimapping rates (Fig. S14). Given the small difference in mapping rate differences between species when mapping to the *T. fasciculata* genome, we decided to conduct our DGE analyses on this genome primarily. Interestingly, *T. fasciculata* samples map equally well to both the *T. fasciculata* and *T. leiboldiana* genome in terms of uniquely mapping reads, while the reverse is not true (*T. leiboldiana* unique mapping rates decrease when mapped to *T. fasciculata*). For multimapping reads, rates are typically lower when samples are mapped to their conspecific reference genome.

Mapping of the full dataset (72 samples) to the *T. fasciculata* genome resulted in an average uniquely mapping rate of 85.7 % ± 5.25 % for *T. fasciculata* samples and 85.19 % ± 0.61 % for *T. leiboldiana* samples. When mapping to the *T. leiboldiana* genome assembly, 85.4 % ± 5.25 % of *T. fasciculata* reads mapped uniquely, while *T. leiboldiana* samples had an average unique mapping rate of 93.4 % ± 1.46 %.

### Note 9: Differential gene expression using the *T. leiboldiana* assembly

In addition to the DE analysis using the *T. fasciculata* genome as reference, we performed a second time-dependent DE analysis in maSigPro with the *T. leiboldiana* genome as reference to test whether the one-directional enrichment of multi-copy gene families is the result of a technical bias when using the *T. fasciculata* genome for DE analysis. It is possible that differential gene expression in additional copies of *T. leiboldiana* are missed, as these are not present in the *T. fasciculata* genome and may be too divergent to map onto a different copy.

The DE analysis on the *T. leiboldiana* reference genome recovered 863 genes with differential expression profiles in a 24-hour period between species. These belonged to 714 orthogroups, of which 458 overlapped with the set of DE orthogroups recovered in the DE analysis using *T. fasciculata* as reference genome. This is equivalent to 66 % of non-unique DE orthogroups called with *T. fasciculata* as reference and 70 % called with *T. leiboldiana* as reference. The overlap in DE orthogroups is lowest for expanded gene families in the respective genome, with only 31 % and 33 % of orthogroups larger in *T. fasciculata* and *T. leiboldiana* respectively containing DE genes in both analyses. This highlights the importance of performing timewise DE analyses on both genome assemblies.

We indeed find enrichment for gene families with gene counts higher in *T. leiboldiana*, which occur twice as often in the DE subset compared to the whole genome (Chi-square P = $1.011568e^{-33}$, Fig. 5b). We also find a small increase of multicopy families with higher gene counts in *T. fasciculata* compared to the whole genome when using mapped reads to *T. leiboldiana* (See Results). 59 known genes related to CAM, gluconeogenesis and starch metabolism were part of the subset of DE genes detected using *T. leiboldiana* as reference genome. 49 out of the 57 orthogroups that these genes belonged to were also recovered as DE orthogroups in the DE analysis using *T. fasciculata* as reference genome. The remaining non-overlapping eight orthogroups consist of seven one-to-one orthogroups and one multi-copy orthogroup with equal counts (3:3) with one DE gene. The 14 CAM-related DE orthogroups that were only called when using *T. fasciculata* as an orthogroup consist of four one-to-one orthogroups, eight orthogroups expanded in *T. fasciculata* and two multi-copy orthogroups of equal size. Fisher's exact test on each category of DE orthogroups related to CAM, starch metabolism and gluconeogenesis found in the *T. leiboldiana* analysis shows that, unlike for all DE orthogroups, there is no significant enrichment of gene families expanded in *T. leiboldiana*. This suggests that gene family expansion in CAM-related genes is specifically found in the CAM lineage.

**Note 10: On the differential expression and gene family expansion of CAM-related genes**

We summarised our findings on evolutionary processes affecting CAM-related genes in figure 6, by highlighting if a gene was affected by differential gene expression, gene family expansion, high TE insertion rates or adaptive sequence evolution in or between *T. fasciculata* and/or *T. leiboldiana*. Here we give a more detailed explanation of this figure and discuss expression profiles of affected genes, which can be found in a compiled PDF here: https://github.com/cgrootcrego/Tillandsia_Genomes.

Starting with the night-time carbon fixating module of CAM, we find gene family expansion in β-carbonic anhydrase (CA) 5 in *T. fasciculata*, though no significant differential gene expression. The pineapple ortholog of this family showed diel expression and circadian regulation (Wai et al., 2017). CA converts $CO_2$ to $HCO_3^-$, which is then added to phosphoenolpyruvate (PEP) by Phosphoenolpyruvate carboxylase (PEPC). PEPC underwent an ancient duplication shared by pineapple and the two *Tillandsia* species, but only one copy (Tfas_v1.16311) shows a diel pattern in *T. fasciculata* and not in *T. leiboldiana* (Fig. 5c). The other copy is lowly expressed in both species.

An important regulator of PEPC is PEPC kinase (PPCK), which reduces PEPC's sensitivity to malate inhibition and renders it more active by phorsphorylation during the night (Nimmo et al., 2001) . PPCK (Tfasc_v1.03128-RA) shows accentuated diel expression in *T. fasciculata* compared to *T. leiboldiana* (Fig. S7). PPCK also shows an increase in night-time expression in *T. leiboldiana*, which has been reported previously in other C3 *Tillandsia* (De La Harpe et al., 2020) and in *Yucca* (Heyduk et al., 2022), suggesting a conserved time-structured expression of PPCK.

The product of PEPC's catalysed reaction is oxaloacetate, which is converted to malate by malate dehydrogenase (MDH). We detect a MDH gene family (OG0001548) with differential expression between *T. fasciculata* and *T. leiboldiana* which also shares a duplication across all three species. Similar as with PEPC, one copy shows a clear diel pattern in *T. fasciculata* while the other copy is lowly expressed (Fig. 5c). Another MDH gene family that is expanded in *T.*

*fasciculata* (OG0002059) shows expression in both copies with a slight phase shift from *T. leiboldiana*, peaking in the late night.

The resulting malate is then transported to the vacuole, putatively by aluminium-activated malate transporters (ALMT) (Kovermann et al., 2007; Wai et al., 2017). The single-copy gene Tfasc_v1.09028-RA, which codes for aluminum-activated malate transporter 9 and is orthologous to the candidate CAM malate transporter in pineapple (Wai et al., 2017), was differentially expressed – though it shows diel expression in both *Tillandsias* and is more highly expressed in *T. leiboldiana*. This gene also shows elevated TE insertion rates in *T. fasciculata*.

Night-time malate transport into the vacuole is energized by two vacuolar pumps: vacuolar $H^{+-}$ATPase pump (V-ATPase) and $H^{+}$-PPiase (V-PPiase, AVP1), which help maintain the ionic gradient of the tonoplast (Wai et al., 2017; McRae et al., 2002). Several subunits of V-ATPase are differentially expressed, showing a variety of expression patterns, as they are spread out across four different expression clusters (cluster 2, 3, 4 and 5, table S7), including elevated expression in *T. fasciculata* and in *T. leiboldiana*, suggesting a complex regulatory structure. Subunit A and B, which show high expression in CAM-performing tissues in pineapple, were highly expressed in *T. fasciculata* compared to *T. leiboldiana* and showed a clear diel pattern with elevated expression at night. Interestingly, a few DE subunits have undergone gene family expansion in *T. fasciculata* (See Table S5). Subunit H (OG0000539) contains two DE gene copies and eight gene copies in total in *T. fasciculata*, as opposed to three in *T. leiboldiana*. Expression patterns of these two gene copies show elevated and diel expression in one (Tfasc_v1.19742) and reduced, non-diel expression in the other one (Tfasc_v1.24696) compared to *T. leiboldiana*. The latter gene also has an elevated TE insertion count in *T. fasciculata*, suggesting the pseudogenization of a gene that is expressed in the C3 plant, and differential co-opting of gene copies between a C3 and CAM plant. Subunit C (Tfasc_v1.17712-RA) also shows elevated TE insertion rates in both species and has an elevated, diel expression profile in *T. fasciculata*.

We also detect differential expression in a copy of AVP1, a secondary vacuolar pump, which has more gene copies in *T. fasciculata* than in *T. leiboldiana*. Two copies show diel cycling in *T. fasciculata*, peaking during the night, but one *T. leiboldiana* copy is much more highly expressed and shows a phase shift as it peaks in the late day. As found in pineapple, AVP1 is much more lowly expressed than V-ATPase subunits, however its expression profile differs from that of pineapple (peaking at midday) (Wai et al., 2017).

During daytime, malate is transported out of the vacuole and reconverted to PEP. There are two possible metabolic routes known in CAM plants, either by converting malate to oxaloacetate by MDH and decarboxylation to PEP by PEP carboxykinase (PEPCK), or by decarboxylating malate into pyruvate by malic enzyme (ME) and converting pyruvate into PEP with pyruvate orthophosphate dikinase (PPDK). In pineapple, the main carboxylase is PEPCK (Dittrich et al., 1973), and indeed we do not recover differential temporal expression in ME or PPDK, while PEPCK shows a strong diel pattern peaking during the day and dropping at night. This indicates that the daytime CAM metabolism is conserved between pineapple and *Tillandsia*.

During the day, CAM plants also accumulate transitory starch in the chloroplast and/or soluble sugars in the vacuole, which become a source of PEP at night. In Bromeliaceae, the relative importance of starch versus soluble sugars as sources of PEP is variable across species, with some species accumulating only starch, others, only soluble sugars, and others both (Christopher and Holtum, 1998). We find differentially expressed genes involved in both pathways: vacuolar transporters which bring soluble sugars into the vacuole (SUT2 and ERD6) on the one hand, and starch synthase, α- and β-amylase and glucose-6-phosphate/phosphate translocator (GPT) for starch accumulation and phosphorylitic degradation on the other hand. Two families of vacuolar invertases, which convert sucrose to hexoses in the vacuole, show gene family expansion in *T. fasciculata* (OG0001710) and elevated TE insertion rates in *T. leiboldiana* (OG0015868). A starch synthase family that is expanded in *T. fasciculata* (OG0001599) is also differentially expressed, showing diel expression peaking during the day in *T. fasciculata* while being very lowly expressed in *T. leiboldiana* with no diel cycling.

To maintain the supply of PEP for night-time carbon fixation, vacuolar hexoses and/or transitory starch is broken down through glycolysis (Christopher and Holtum, 1996). Indeed, several glycolytic enzymes are differentially expressed and clustering together with PEPC in cluster 5, which shows diel expression peaking in the late afternoon to the early night in *T. fasciculata*: glucose-6-phosphate isomerase (Tfasc_v1.04514-RA), phosphofructokinase (Tfasc_v1.20370-RA, Tfasc_v1.24086-RA) and aldolase (Tfasc_v1.17464-RA). ATP-dependent phosphofructokinase however peaks in expression during the day, while an isoform of the expanded PPi-dependent phosphofructokinase family OG0000580 shows elevated expression in the early night. This could indicate an important role for soluble sugars stored in the vacuole as a source for night-time PEP, given that PPi-dependent phosphofructokinase cannot access fructose-

6-phosphate precedent from starch accumulated in the chloroplast (Carnal and Black, 1989). A hexokinase was also called as differentially expressed in the *T. leiboldiana* genome with elevated expression in *T. fasciculata*. We also find two expanded cytosolic enolase families in *T. fasciculata* (OG0000469 and OG0000555) and one in *T. leiboldiana* (OG0001563), and an expanded glyceraldehyde-3-phosphate dehydrogenase in *T. leiboldiana* (OG0000320). The enolase family OG0000555 contains two copies with elevated night-time expression in *T. fasciculata*, while GAPDH shows one copy in each species with high expression and a phase shift between species, with expression peaking at night.

While exact regulators of CAM are not as well described as the pathway itself, we do encounter some differentially expressed circadian clock regulators which have been described previously in the context of CAM. Three DE circadian genes, GIGANTEA (GI), REVEILLE 1 (RVE1) and LATE ELONGATED HYPOCOTYL (LHY) show a similar expression pattern with reduced diel oscillation in *T. fasciculata* compared to *T. leiboldiana*. None of these genes showed phase displacement between the two species, which differs from findings between *Agave* and *Arabidopsis* (Yin et al., 2018), particularly for RVE1. While none of these genes show gene family evolution, GI has a high number of TE insertions in both species (Table S10).

Lastly, we also recover differentially expressed regulators of stomatal movement such as ATP-binding cassette C14 (ABCC14) and succinate dehydrogenase (SDH) which exist in a multi-copy gene family. In both enzymes, one gene copy in *T. leiboldiana* shows stronger diel cycling than the copies in *T. fasciculata*. While ABC-C14 peaks during the day, SDH peaks in the late night in *T. leiboldiana*. The highest expressed copy of ABC-C14 in *T. fasciculata*, which peaks in activity in early night, also has an increased number of TE insertions.

### Note 11: On the success of *de novo* assembly of highly repetitive genomes

The highly repetitive content observed in many plant genomes often causes fragmented genome assemblies. Despite considerable progress thanks to long-read sequencing technologies, little to no genomic resources are available yet for some plant clades, particularly for species with the remaining challenge of a highly repetitive content (Marks et al., 2021). The availability of long-read sequencing and chromatin conformation capture technologies have now enabled the assembly of particularly complex genomes with little fragmentation, in the best case at the chromosome-level. Our project was therefore launched and made possible because of these recent technological advances.

The kmer-based approach implemented in findGSE (k = 21, see also SI Note 1 and 2) estimates the TE content directly from raw reads, *i.e.,* prior to generating *de novo* assemblies. We estimated that the repetitive content of *T. fasciculata* and *T. leiboldiana* was around 66 % and 75 %, respectively. Based on *de novo* generated assemblies and TE annotations performed with EDTA (Ou et al., 2019), we observed remarkably consistent estimates (65.5 % and 77.1 % in *T. fasciculata* and *T. leiboldiana*, Table S3) on the main scaffolds (SI Note 3). Such values are one of the most elevated among plant genomes assembled at chromosome scale (Pedro et al., 2021).

Additionally, our analyses of spatial distribution of TE content highlight the extreme local levels of repetitive content in these genomes, especially in *T. leiboldiana*. In non-telomeric regions, the observed repetitive content most often reaches values above 80% for *T. fasciculata* and nearly 100% for *T. leiboldiana* (Fig. 3b, Fig. S3).

Considering the extremely high repetitive content in centromeric regions, especially for *T. leiboldiana*, the limited fragmentation of our assembly can be considered as a success and therefore represent empirical evidence of the progress made in plant genomics, only twenty years after the release of the first plant genome. At the age of long-read sequencing and chromatin conformation capture technologies, the *de novo* sequencing of plant species associated with a highly repeated genomic content is becoming more and more feasible.
